# Diverse Protein Architectures and α-*N*-Methylation Patterns Define Split Borosin RiPP Biosynthetic Gene Clusters

**DOI:** 10.1101/2021.12.24.474128

**Authors:** Aman S. Imani, Aileen R. Lee, Nisha Vishwanathan, Floris de Waal, Michael F. Freeman

## Abstract

Borosins are ribosomally synthesized and post-translationally modified peptides (RiPPs) with α-*N*-methylations installed on the peptide backbone that impart unique properties like proteolytic stability to these natural products. The borosin RiPP family was initially reported only in fungi until our recent discovery and characterization of a Type IV split borosin system in the metal-respiring bacterium *Shewanella oneidensis*. Here, we used hidden Markov models and sequence similarity networks to identify over 1,600 putative pathways that show split borosin biosynthetic gene clusters are widespread in bacteria. Noteworthy differences in precursor and α-*N*-methyltransferase open reading frame sizes, architectures, and core peptide properties allow further subdivision of the borosin family into six additional discrete structural types, of which five have been validated in this study.

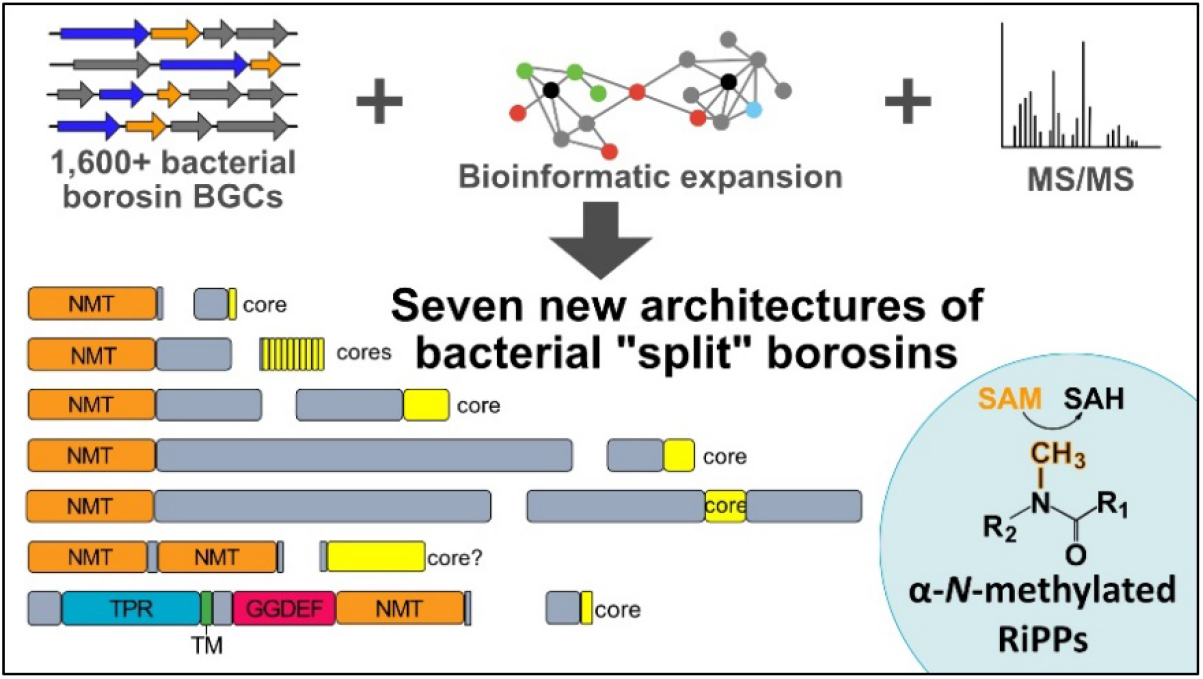

## Introduction

Early top-down approaches to natural product discovery relied heavily on culturing microbes and performing bioactivity-guided screens. Scores of important and potent antibiotics, anticancer agents, and immunosuppressants were discovered using this approach. Over time, as much of the “low-hanging fruit” was picked, rediscovery rates increased and the identification of new bioactive scaffolds plummeted.^1^ However, rapid advances in DNA sequencing technologies and bioinformatic tools in the 21^st^ century has reinvigorated the field of natural products. The plethora of accessible (meta)genomic data has enabled a genomics-driven approach to natural product discovery that offers insight into the true biosynthetic potential of microbes through culture-independent methods.^2^

One class of natural products that has benefited from *in silico* approaches is ribosomally synthesized and post-translationally modified peptides (RiPPs).^3^,^4^ RiPPs are produced as genomically encoded precursor peptides composed of a leader and core peptide (Figure 1). Tailoring enzymes, typically encoded alongside the precursor gene in a biosynthetic gene cluster (BGC), recognize the leader peptide and modify the core peptide. The final steps in RiPP biosynthesis usually involve proteolytic cleavage of the core peptide and export of the mature natural product. While RiPPs are limited to the 20 canonical amino acids as building blocks, exquisite chemical diversity is generated by hypervariable core sequences and extensive post-translational modifications (PTMs) that enhance their stability and activity.^5^

**Figure 1.**
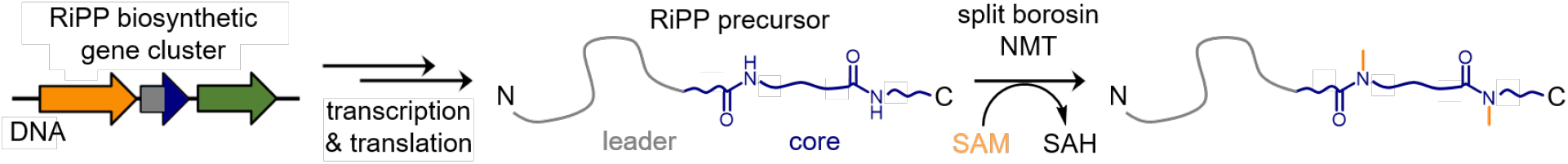
RiPP precursor peptide maturation. Borosin precursor peptides are α-*N*-methylated on the peptide backbone through an *S*-adenosylmethionine (SAM)-dependent process. SAH = *S*-adenosylhomocysteine.

Along with our collaborators, we and others identified the fungal nematicide omphalotin A as the first RiPP natural product with amide backbone α-*N*-methylations.^6,7^ Once thought to be an exclusive feature of non-ribosomal peptides, amide backbone α*-N*-methylation imparts peptides with properties that include enhanced backbone rigidity, membrane permeability and resistance to proteases, making these modifications attractive for pharmacological development. In omphalotin biosynthesis, an α-*N*-methyltransferase domain (NMT) in the RiPP precursor OphMA is fused to the core peptide substrate and iteratively α-*N*-methylates the omphalotin A sequence in an N- to C-terminal fashion.^6^ We named this family of α-*N*-methylated RiPPs the borosins, with the omphalotins as their founding members. Soon after this discovery, the cytotoxic gymnopeptides^8^ and the omphalotin-related lentinulin A and dentrothelin A^9^ were also found to be from the borosin RiPP family. A more thorough analysis of the fungi-derived borosin pathways revealed different protein architectures of the fused methyltransferase-encoding precursors. Based on differences found in the leader and core peptides, the precursors in the putative borosin pathways were categorized into three structural types (Figure 2A).^8^

**Figure 2.**
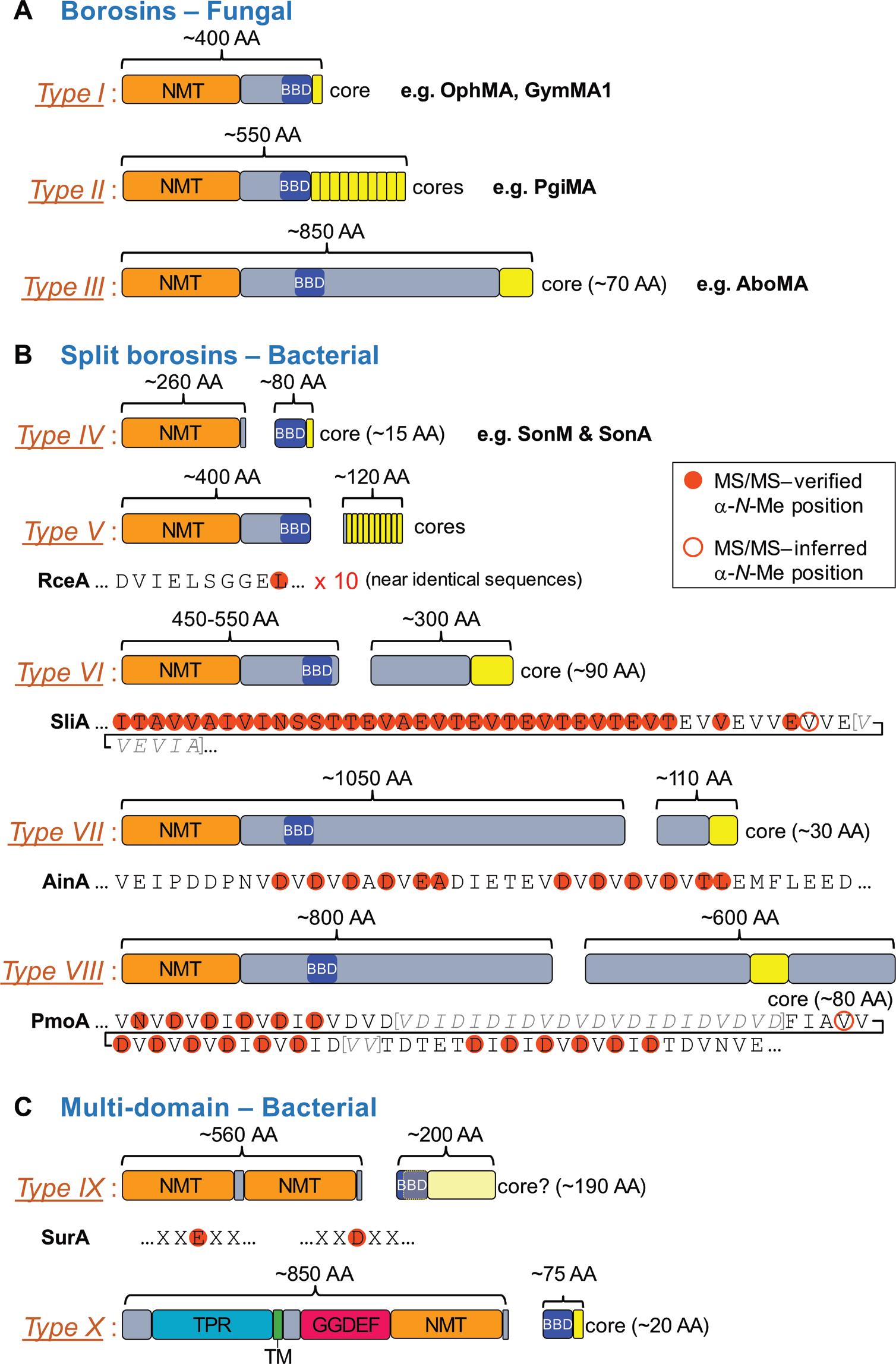
Borosin α-*N*-methyltransferases and precursor peptides are subdivided into Types I-X based on overall protein architecture and core peptide composition. Cartoon representations are displayed for borosin α-*N*-methyltransferases (NMTs) and/or precursor peptide structural types. Borosin binding domains (BBDs) as predicted by HHpred^10^ are depicted as blue domains, core peptides are displayed as yellow segments, and sequences with no predicted function are displayed in grey. **(a)** Previously characterized Type I-III fused borosin systems from fungi listed alongside previously verified examples. **(b)** ‘Split’ borosins Types IV-VIII identified in bacteria. Types V-VIII are newly described in this study. Putative core peptide sequences show α-*N*-methylation patterns as inferred from global LC-MS/MS analysis of precursor peptides from in vivo *E. coli* expressions with their cognate NMTs: RceA/M, *Rhodospirillum centenum* SW; SliA/M, *Spirosoma linguale* DSM 74; AinA/M, *Achromobacter insuavis* AXX-A; PmoA/M, *Pseudomonas mosselii* CIP 105259. Sequences below protein architectures shown in black text were identified by LC-MS/MS; sequences in bracketed grey italics were not identified by LC-MS/MS. **(c)** Multi-domain-encoding bacterial ‘split’ borosins Types IX and X (new to this study). Diffuse methylation patterns of Asp and Glu residues are observed with the Type IX borosin system SurA/M1 from *Streptomyces ureilyticus*. As the core peptide is not well defined, the entire precursor is depicted in pale yellow. TPR = tetra-trico-peptide repeat domain; TM = trans-membrane domain; GGDEF = diguanylate cyclase domain.

Crystallographic interrogation of the Type I omphalotin-encoding precursor OphMA spurred a proposal for the molecular mechanism of catalysis.^11^ Briefly, water-mediated proton abstraction from the target amide forms an imidate stabilized by an oxyanion hole. Subsequent nucleophilic attack on the methyl-donating cofactor *S*-adenosylmethionine (SAM) yields an α-*N*-methylated amide and *S*-adenosylhomocysteine (SAH) as byproducts. This process occurs iteratively in an N-to-C direction on the core peptide to create the omphalotin A α-*N*-methylated backbone.

The diversity of substrate-fused NMT architectures across fungi raised the intriguing question of whether bacteria also harbor borosin BGCs. Iterative PSI-BLAST searches of the OphMA NMT domain yielded putative hits that included one from the metal-respiring bacterium *Shewanella oneidensis* MR-1.^12^ Unlike the canonical substrate-fused NMT, this putative borosin BGC followed traditional RiPP biosynthetic logic with discrete NMT and precursor peptide open reading frames. Mass spectrometric and kinetics analysis revealed α-*N*-methylation of two residues in the core peptide, indicating methylation occurred truly *in trans* unlike previously identified substrate-fused Type I-III borosin systems. Crystal structures of fully resolved NMT-precursor complexes illustrate a five-helix bundle termed the borosin binding domain (BBD) as the dominant interaction domain of the precursor peptide with the NMT. In addition, stark conformational changes observed amongst different NMT-precursor mutants sparked a hypothesis that core peptide secondary structure changes direct iterative N-to-C α-*N*-methylation. Kinetic analysis revealed multiple substrate turnover and an efficient system with an apparent *k*_cat_ of 0.52 min^−1^ compared to 0.0053 min^−1^ for single peptide turnover observed with OphMA. Divergence of the *S. oneidensis-type* borosin protein architectures from the canonical fused Type I-III fungal systems warranted the designation of Type IV borosins informally referred to as split borosins.^12^

In this work, we perform extensive analyses and verification of split borosin pathways to reveal a diverse family of RiPPs widespread across bacteria. We validate five putative bacterial split borosin NMT and precursor pairs with distinct methylation patterns amongst six additional borosin structural types (Types V-X). Bioinformatic analysis of split borosins from major bacterial groups detected in BLASTP searches uncovered >1600 putative split borosins BGCs amongst many of the architectural types. The combination of *in vivo* and *in silico* data analysis allowed us to develop a set of rules using hidden Markov models (HMMs) from conserved borosin NMT regions to allow researchers to detect and mine putative borosin gene clusters in programs such as antiSMASH.^13^ The discovery and detailed analyses of borosin BGCs open the door to many putative bioactive metabolites and biotechnology applications for amide backbone α-*N*-methylated peptides.

## Results and Discussion

### Split borosin α-*N*-methyltransferase and precursor architecture variability

Due to the variable architectures of fused RiPP precursors observed in fungal borosin structural Types I-III, we sought to determine whether the newly identified bacterial split borosin NMTs and precursors were similarly diverse in size and sequence. Manual inspection was initially performed on bacterial genetic loci surrounding PSI-BLAST hits in bacterial genomes using the omphalotin methyltransferase domain in OphMA as a query. Interestingly, putative NMTs ranged in length from ~250 amino acids (AAs) to over 1000 AAs (Figure 2B). Heterologous expression of select bacterial NMTs alone followed by high-resolution, high-pressure liquid chromatography-mass spectrometric (LC-MS/MS) analysis did not yield observable autocatalytic self-methylation as in the fungal borosin systems.

Like with many RiPP classes, identification of split borosin precursors was not straightforward, albeit for different reasons than the typically hard-to-identify, short hypervariable precursor peptides.^4^ While some putative borosin pathways encoded relatively short peptides (50-100 AAs) homologous to SonA and were easily pinpointed, many BGCs did not. Careful analysis of the encoded surrounding genes revealed several large proteins of unknown function, some in excess of 600 AAs. Luckily, a subset of these putative proteins contained sequence repeats reminiscent of the cores in fungal borosin Type II and Type III pathways^8^ and were subsequently flagged as possible precursors.

In order to validate putative split borosin BGCs, NMTs and putative precursors were categorized into additional structural types based on their overall protein sizes and compositions (Figure 2B&C). Precursor-NMT pairs from each new structural type were then heterologously expressed in *E. coli* and purified via nickel-chelate chromatography. Precursors from *in vivo* coexpressions with their cognate NMTs were subsequently digested and analyzed by LC-MS/MS. Through extensive manual analysis and use of the proteomics software MaxQuant,^14^ a wide variety of split borosin core peptides in structural Types V-IX were found to be extensively α-*N*-methylated (Figure 2B, Figure S1).

### Split borosin Types V-IX and their diverse methylation patterns

Ordering of the split borosin architectural types was based on the collective length of NMT and precursor, similarly to what was reported for the fused borosin fungal systems Types I-III, and continuing from the first Type IV split borosins identified in bacteria.^12^ Type V split borosins were verified through expression of RceM & RceA from *Rhodospirillum centenum* SW. Similarly to the fused fungal Type II borosins, Type V split borosins are signified by a multi-core precursor peptide, where a single methylation is incorporated in each near-identical core repeat. Ten near-identical core peptides sequences of ‘DVIELSSGGEL’ are found in precursor RceA. Due to proteolytic limitations for MS/MS analysis, the mutant RceA S78F was also created and analyzed by LC-MS/MS to provide evidence for methylation in all ten copies of the core sequence repeat (Figures S1&S2). Curiously, the borosin binding domain (BBD), shown crucial for precursor peptide docking with the NMT of Type IV borosins, was predicted by HHpred^10^ to be encoded in the RceM NMT domain and not within the short RceA leader peptide (Figure 2B). Consequently, the structural determinants required for precursor—NMT interactions in Type V split borosins may differ from those observed in Type IV borosins.

For split borosin Types VI-VIII, we identified putative precursor genes adjacent to the NMTs encoding repetitive sequences reminiscent of the methylated ‘DVDVT’ repeats found in the fungal Type III borosin AboMA.^8^ Each of Types VI-VIII have distinct architectures and methylation patterns. For example, SliA (Type VI), a putative precursor protein encoded in a BGC from *Spirosoma linguale* DSM 74, is a 300-AA protein containing several TEVX repeats (X = Thr, Val, Ala). A defining feature of Type VI split borosins is the successive methylation pattern observed within its core peptide. Combining the results of several LC-MS/MS verified peptide fragments, we observed 32 consecutive α-*N*-methylated amino acids (Figure 2B, Figure S1). Due to the difficulties in overexpression, purification, as well as a high incidence of soluble aggregate formation with many of these new split borosin NMT-precursor pairs (Figure S3), in-vitro characterization has proven cumbersome and complete methylation of the core peptide has not been achieved for Types VI and VIII. For instance, we hypothesize up to 48 successive methylations are incorporated into SliA by SliM.

A relatively short precursor protein (~110 AA) encoding two small sets of ‘DVDV’ repeats and an unusually large NMT in excess of 1000 AA signifies Type VII split borosin pairs. Type VII split borosins were verified through coexpression of AinA and AinM from *Achromobacter insuavis* AXX-A (formerly identified as *A. xylosoxidans*).^15^ Interestingly, methylation of the acidic residues displays an opposite pattern as compared to the AboMA Type III fused borosin, where methylation occurs on Val and Thr residues (Figure S1). Despite substantial mutational analyses and core peptide replacements, the ‘rules’ for α-*N*-methylation incorporation into borosins are still enigmatic.^6,16^ A combination of sequence and secondary structural features of the core peptide likely contribute to the fidelity of borosin methylation in each RiPP system.^12^

Similarly to Type VII, the Type VIII split borosin systems also incorporate methylations of acidic residues in ‘D[V/I]D[V/I]’ repeats. However, the substantial size of the precursor peptide (~600 AA) along with the location of the core peptide in the middle of the protein makes this RiPP precursor quite an unusual example and justifies its own division among the split borosin BGCs (Figure 2B).

The affixed domains found in the split borosin precursors and larger α-*N*-methyltransferases garnered interest as to whether additional catalytic domains might be encoded in these genes. Beyond the autocatalytic fungal borosin methyltransferases, RiPP precursors have been recently found in plants to encode repetitive core sequences and a C-terminal BURP domain that functions as a macrocyclase.^17^ However, no additional catalytic domains were identified in NMTs or precursors in Types VI-VIII by programs such as HHpred.^10^ As an alternative method, we ran all of our newly identified NMT-precursor pairs for structural predictions through AlphaFold.^18^ While AlphaFold did give predicted folds for several of the additional domains (Figure S4), the predictions are based off very few homologous sequences and the resulting low-confidence structures have yet to reveal additional catalytic functions. Despite our best efforts, we have not yet observed PTMs beyond α-*N*-methylation in Type VI-VIII split borosin NMT-precursor coexpressions.

In contrast, a subset of split borosin α-*N*-methyltransferases do appear to encode additional catalytic domains. Borosin NMTs can be found fused to the C-terminus of GGDEF-containing proteins, similar to what is found in the *S. oneidensis* Type IV borosin BGC (Figure 2C).^12^ GGDEF proteins produce cyclic di-GMP, a secondary messenger in bacteria that has been linked to the lifestyle decision between motility and biofilm formation.^19^ These proteins are predicted to span the inner membrane with a tetracopeptide repeat (TPR) domain^20^ displayed in the periplasm. Another multi-domain example (Type IX) found in *Streptomyces ureilyticus* appears to have a duplicated and fused second borosin methyltransferase domain (Figure 2C). As expected, heterologous expression and purification revealed that Type IX SurM1 from *S. ureilyticus* is monomeric in solution as observed by size exclusion chromatography unlike all other borosin NMTs analyzed to date (Figure S3). Coexpression with its putative precursor protein, SurA, with or without an additional borosin NMT encoded in the BGC, SurM2 (a Type IV borosin NMT with only one catalytic domain), resulted in methylations distributed throughout SurA on Asp and Glu residues, even within the predicted BBD (Figure S1). However, repeated expressions of SurM1 and SurM2 alone or together with the precursor SurA resulted in a subset of different Asp and Glu methylated residues. As such, more information is needed to define the maturation of the precursor in this system.

### Sequence Similarity Network Analysis

To determine the abundance of different borosin architectures distributed in nature, a sequence similarity network was constructed of OphMA homologs identified through BLASTP. The BLASTP search was conducted by querying the α-*N*-methyltransferase domain (residues 1-250) of OphMA against the non-redundant protein database (April 28, 2021). Because an extensive overview of borosins found in fungal genomes has been previously explored,^8^ only non-fungal sequences were queried. The search returned 1804 sequences with E-values ranging from 4.00 × 10^−94^ to 0.037 (median of 3.0 × 10^−45^) and 21.4–58.2% identity (median of 39.2%) with the longest sequence comprising 1151 AA (Supplemental Dataset S1). The 1804 sequences are derived from sequenced genomes of 948 distinct organismal strains: 934 bacteria, 11 archaea, and 3 eukaryota. Within the bacterial representatives, the vast majority are classified as Gammaproteobacteria (66.1%), in particular the classes of Alteromonadales (23.4% of all BLASTP hits) and Xanthomonadales (22.5% of all BLASTP hits). Comparatively, available Gammaproteobacteria, Alteromonadales, and Xanthamonadales represent only 42.9%, 0.5%, and 1.3% of available non-fungal RefSeq genome assemblies, respectively, (as of April 28, 2021). Of the archaea, eight of the 11 (72.7%) representatives are Halobacteria, though Halobacteria comprise only 41.5% of Archaeal RefSeq assemblies. Interestingly, the eukaryotic sequences are from the sea anemones *Actinia tenebrosa* and *Exaiptasia diaphana*, and the mollusc *Pecten maximus*, and have E-values between 1.0 × 10^−79^ and 3.0 × 10^−58^. Inspection of the surrounding genes support the notion that the protein sequences are encoded in non-fungal eukaryotes due to the presence of introns, and the most closely related homologs of the surrounding genes also belonging to non-fungal eukaryotes.

A sequence similarity network was constructed by running an all vs. all analysis of the sequences using the Needleman-Wunsch global alignment scoring algorithm.^21,22^ Sequences included in the network are comprised of the BLASTP sequences containing a complete methyltransferase domain (1704 sequences), previously identified borosin α-*N*-methyltransferases found in fungi (55 sequences),^8^ and representative sequences of the YabN_N_like domain sub-family (cd11723)^23^ of tetrapyrrole methylases (76 sequences), for a total of 1834 sequences, with one sequence appearing in the BLASTP hits as well as the YabN group (Supplemental Dataset S2). The YabN_N_like domains were used as an outgroup as it is a subfamily of the TP_methylase domain family (cd09815) that was identified to be closely related to the borosin methyltransferase domains (cd11724).^23^

The functionally verified borosin architectural types are easily distinguishable from one another in the sequence similarity network at a cutoff value of 1.6 (Figure 3). To reduce redundancy, non-representative sequences were identified and removed from the network, resulting in 947 unique sequences. This removal was performed post-clustering to ensure taxonomic diversity within each group was not obscured. In the network, nine of the BLASTP sequences clustered with the outgroup and manual inspection confirmed these sequences to be members of the YabN_like outgroup.

**Figure 3.**
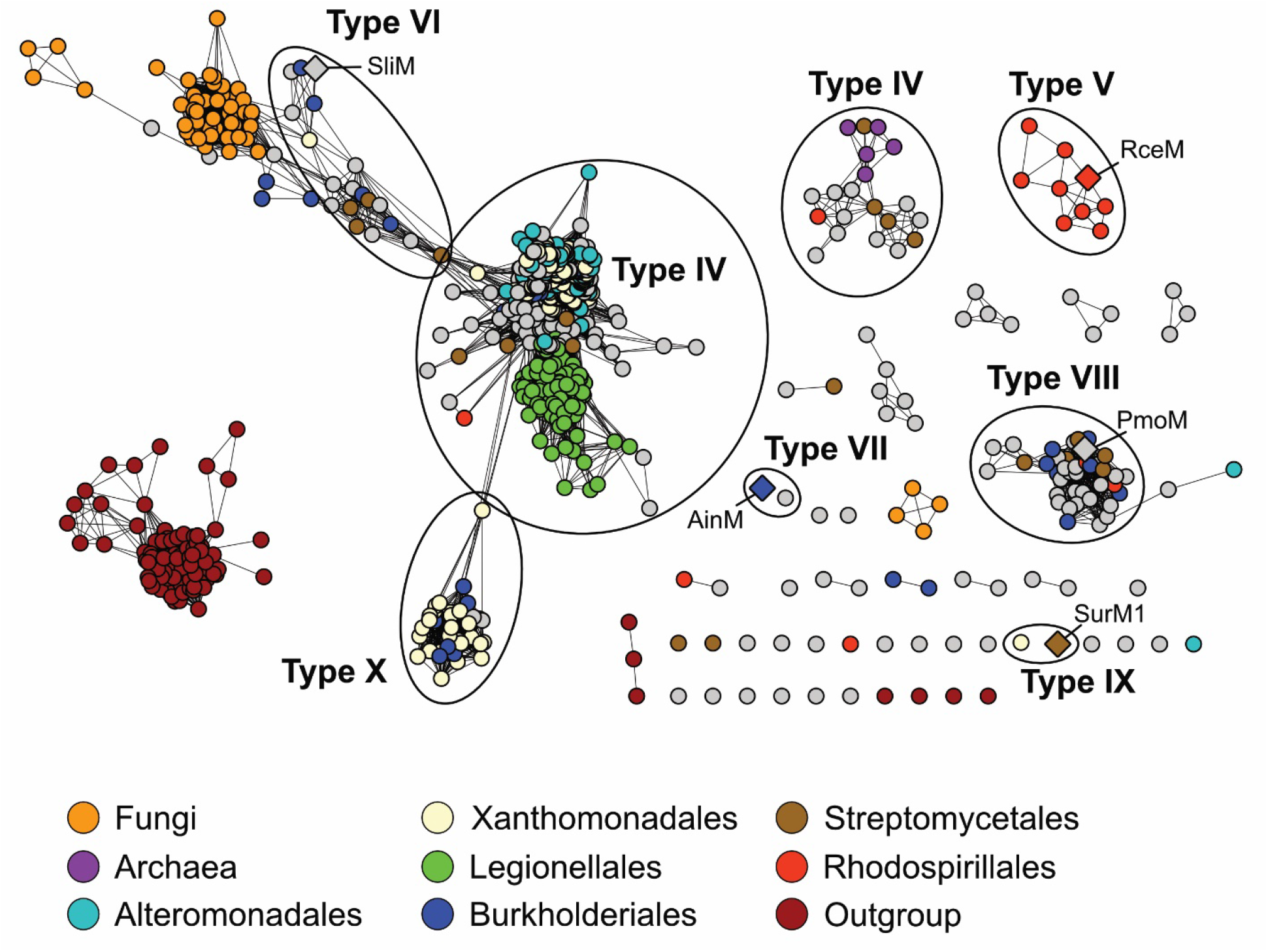
Sequence similarity network of the global architecture of putative borosin NMTs. The sequence similarity network (SSN) consists of sequences from a BLASTP search of the α-*N*-methyltransferase domain of OphMA, sequences of Type I-III methyltransferases involved in fungal borosin biosynthesis (orange), and sequences of YabN_like methyltransferases (maroon). The network was built off a global alignment-based all-vs-all analysis using the needleall module in the EMBOSS suite and visualized in Cytoscape. Nodes corresponding to non-representative sequences (as determined by MMseqs2 with a 0.90 cutoff) were removed. A cutoff value of 1.6 was applied. A select set of nodes are colored by the taxonomic groups to which their host organisms are assigned. Diamonds indicate proteins characterized in this manuscript.

Type IV borosins, which include the well-characterized SonM–SonA split borosin pathway pair, are by far the most commonly found borosins (746 members, 78.8%), with most sequences from this group coming from the orders Alteromonadales (167 sequences, 22.4%) and Legionellales (81 sequences, 10.9%). Type V borosins, represented by the multi-core RceM–RceA split borosin pair, contain only nine members, all from the order Rhodospirillales. Type VI borosin BGCs contain 19 members, including the sequentially methylating NMT SliM, and are most frequently found in Burkholderiales and Streptomycetales with four members each. Type VII borosins contain only two unique members, one from *Achromobacter insuavis* (this study) and the other from an uncharacterized *Magnetococcales* bacterium. Type VIII borosins contain 41 representative members, again with the highest represented groups being Burkholderiales and Streptomycetales (seven and five members, respectively). There were only two identified sequences of the duplicate NMT domain-containing Type IX borosins, one each from Streptomycetales and Xanthomonadales. Finally, the transmembrane-spanning, multi-domain Type X borosins contain 35 representative members with the majority (24 sequences) coming from Xanthomonadales. Curiously, some bacterial sequences remained clustered with the fungal sequences, mainly sequences from Enterobacterales.

A number of additional clusters also appeared in the network. One prominent cluster of note is a 20-member group containing both bacterial and archaeal sequences. Within this cluster, the most highly represented taxonomic groups are Streptomycetales and the archaeal Halobacteriales, each with four members. The overall domain architecture of this group does not appear to differ from that of Type IV borosins; the sequences fall between 210 and 309 residues in length and no other conserved domains were identified. This suggests strong divergence within the sequences of the methyltransferase domains between this group of putative borosins and the general Type IV borosins. Indeed, percent identity to OphMA in the original BLASTP search for this group is an average of 28.5% compared to 38.7% for Type IV borosins. Additionally, a sequence similarity network constructed from an all-vs-all BLAST analysis using only the methyltransferase domains from each sequence shows these sequences breaking off from the main group at a relatively less stringent cutoff value of 1 × 10^−60^ (Figure S5).

This domain-specific sequence similarity network also demonstrates that sequence divergence within the NMT domains is not the predominating factor in the overall separation seen among borosin architectural types in the network built from the global alignment algorithm. Under a relatively stringent cutoff value of 1 × 10^−90^, the Type VI, Type VIII, and fungal (Types I-III) borosin α-*N*-methyltransferase domains remain clustered together (Figure S6). In addition, Type X borosins cluster closely with a small set of Type IV borosins consisting predominantly of sequences from Gammaproteobacteria and Chromatiales. This indicates that the NMT domains are highly similar among these groups, though they differ in overall architecture. Interestingly, sequences from Legionellales dissociate from the main group at relatively low-stringency cutoff values (Figure S7). This is a striking contrast to the global analysis where they remained clustered with other Type IV borosin NMTs, even at high-stringency cutoffs. These observations indicate that, while the overall architecture is consistent with Type IV borosins, the NMT domains in Legionellales diverge from other borosin NMT domains, Type IV or otherwise.

### BGC analysis through BiG-SCAPE

To identify genes with high incidence within borosin BGCs, genome assemblies available in the RefSeq database corresponding to the protein accession IDs from the BLASTP search were downloaded from NCBI. Borosin BGCs were identified by running downloaded assemblies through a locally-installed copy of antiSMASH^13^ containing an added borosin detection rule, BorosinMT, based on highly conserved regions within the α-*N*-methyltransferase domains (Figure 4). Predicted borosin BGC regions were dereplicated using a 90% cutoff value in MMseqs2^24^ before construction of a BGC similarity network in BiG-SCAPE (Figures S8&S9).^25^ Unsurprisingly, BGCs from closely related organisms showed the highest degree of similarity.

**Figure 4.**
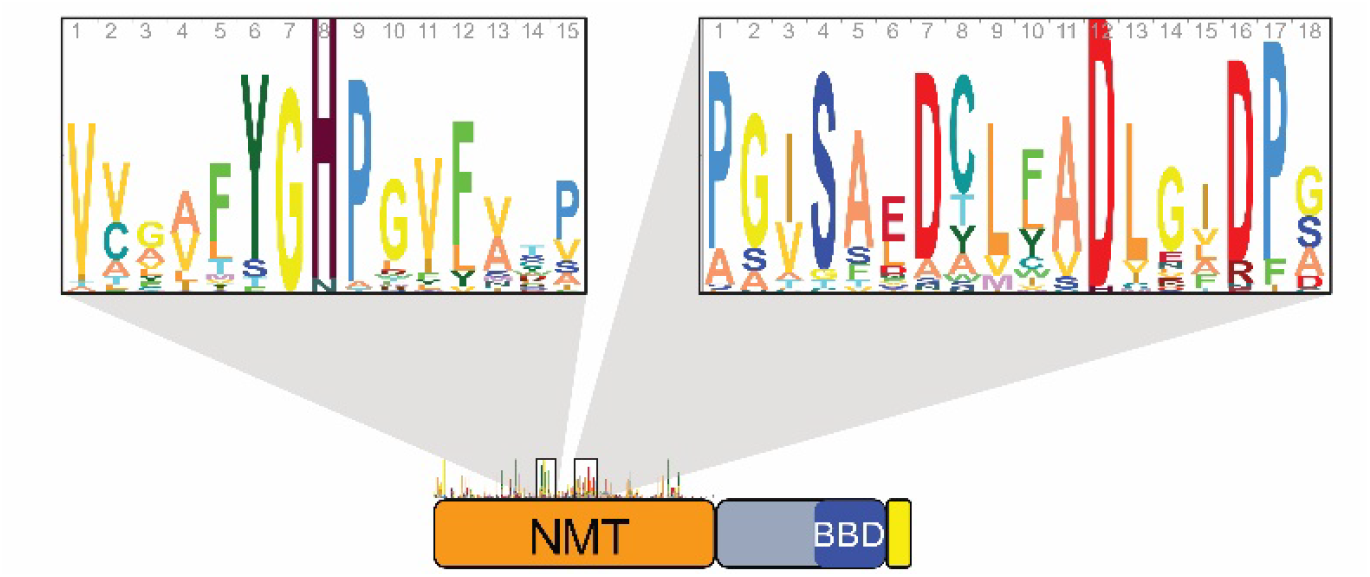
Profile hidden Markov models (HMMs) used to identify borosin BGCs. Sequence logos for the profile HMMs of conserved motifs within the NMT domain, YGHP_v3 (left) and DCLFAD_v3 (right). The profile HMMs were built from an alignment of unique borosin α-*N*-methyltransferase domains. Logos were generated using Skylign.^26^

Looking globally at Pfam domains within the split borosin BGCs, the most common is the GGDEF domain at a frequency of 56%, aside from the NMT-containing TP_methylase domain (Table 1). GGDEF- or GGEEF-domain-containing proteins synthesize the bacterial secondary messenger cyclic-di-GMP.^19^ Cyclic-di-GMP is involved in a variety of cellular processes including metabolic decisions between motility and biofilm formation, exopolysaccharide production, and virulence, among others.^27,28^ In contrast to the Type X borosins where the GGDEF domain is fused to the borosin NMT, the vast majority are encoded as separate genes, similar to what is seen in the *S.oneidensis* BGC. Furthermore, these domains appear in a diverse set of BGCs including those from Types IV, V, VI, VIII, and X borosins in multiple taxonomic groups including Proteobacteria, Actinobacteria, Firmicutes, and Bacteroidetes (Supplementary Dataset 3).

**Table 1.**
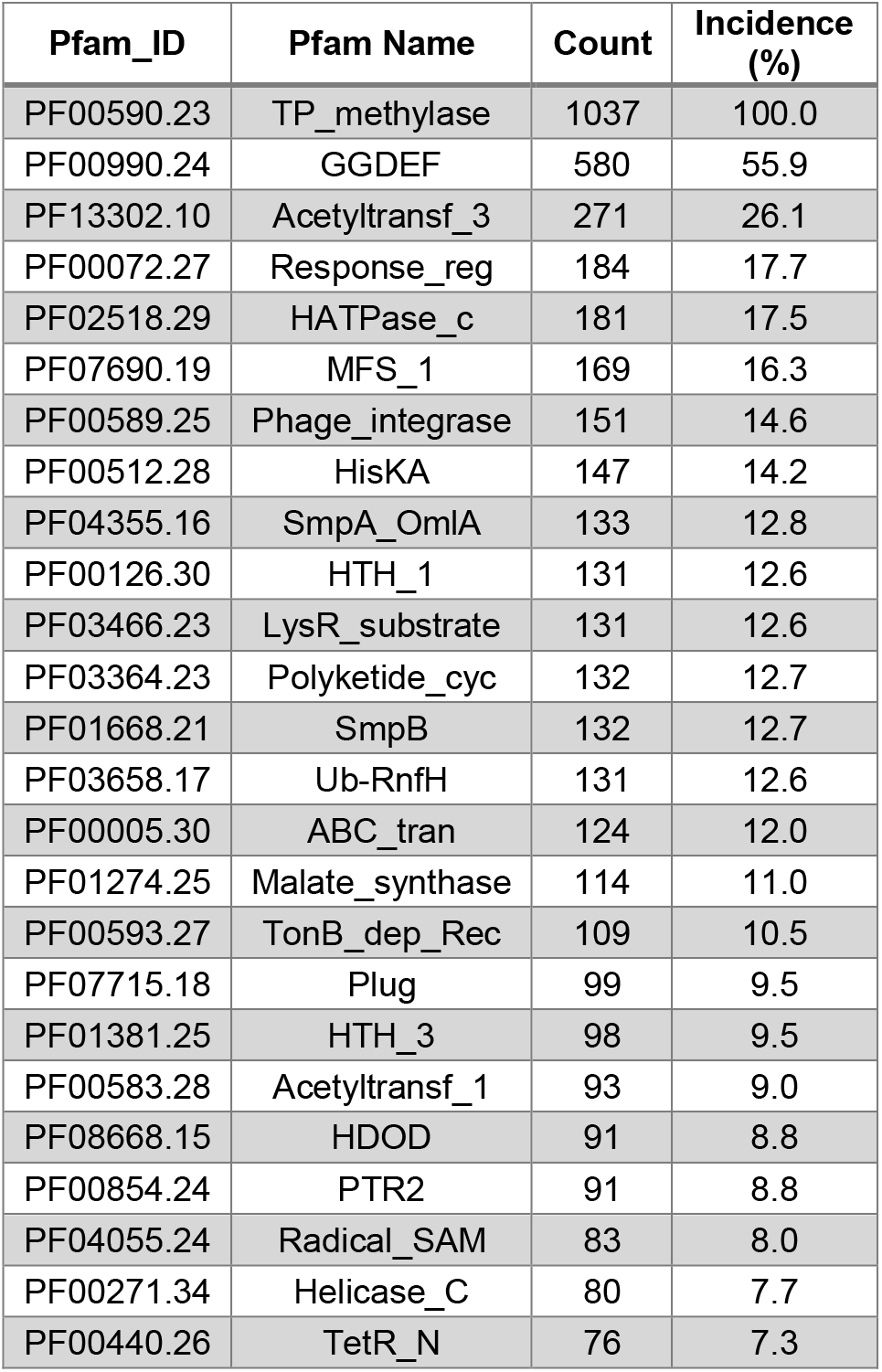
The 25 most common Pfam domains encoded in borosin BGCs, listed in descending order. The complete list of Pfams can be found in Supplemental Dataset S3.

Several frequently occurring Pfam domains distributed across split borosin BGCs are known to be involved in protein maintenance and degradation. PF04355 domains (SmpA_OmlA) include BamE homologs (SmpA, OmlA) that are a part of the BAM complex involved in outer membrane β-barrel protein assembly.^29^ Defects in BamE have been associated with minor defects in outer membrane protein assembly,^30^ activation of the σ^E^-mediated envelope stress response,^31^ and increased susceptibility to various compounds including detergents and antibiotics.^32,33^ Pfam domains PF01668 (SmpB) and PF03658 (Ub-RnfH) are associated with tmRNA-dependent trans-translation protein degradation systems.^34^ Interestingly, the tmRNA gene *ssrA* is directly adjacent to the Type IV split borosin BGC recently identified in *S. oneidensis*.^12^ Many other prevalent Pfam domains found in split borosin BGCs are related to transport, signal transduction, and regulation (Table 1).

Relatively few of the top 25 identified protein domains are associated with biosynthetic enzymes. Two frequently observed Pfams in split borosin BGCs (PF13302, ~26% incidence; PF00583, ~9% incidence) are acetyltransferases known to acetylate a wide variety of small molecule and protein substrates.^35^ Several RiPPs, including LAPs and microviridins, are α-*N*-acetylated at their N-termini.^5^ Approximately 13% of split borosin BGCs encode PF03364-containing proteins that have homology to a variety of polyketide cyclases and desaturases. Although rare, polyketide-RiPP hybrids do exist, as first confirmed with the discovery of microvionin.^36^ The last notable Pfam domain in Table 1 encoding likely biosynthetic enzymes is in the radical SAM superfamily with PF04055 domains identified in 8% of split borosin BGCs. Radical SAM enzymes are a particularly diverse set of enzymes catalyzing a wide array of chemically difficult biosynthetic transformations.^37^ Members of the radical SAM family PF04055 have been found in a wide variety of putative^38^ and known^39^ RiPP families that include the bottromycins,^40^ sactipeptides,^41^ and ranthipeptides.^42^ Despite few highly represented biosynthetic enzyme families found across all split borosin BGCs, individual borosin pathways can encode a number of putative post-translationally modifying enzymes. Of the examples functionally validated in this study, the Type VII BGC from *A. insuavis* encodes several additional biosynthetic enzymes including putative P450, glycosyltransferase, carbamoyltransferase, polyketide synthase, and hydroxylase enzymes (Figure S10). For a more detailed distribution of Pfam domains among closely related borosin BGCs identified through BiG-SCAPE, please refer to Supplemental Dataset 4.

## Conclusions

A combination of advances in DNA sequencing, gene synthesis, and bioinformatics tools have shifted the natural product discovery pipeline in many ways to favor a genomics driven approach. Through a combination of *in silico* analyses and *in vivo* heterologous expressions, we have expanded the borosin RiPP family to encompass an architecturally diverse set of α-*N*-methyltransferases and precursor pairs predominantly found in bacteria. We have developed a set of profile HMMs that, when used in a locally installed version of antiSMASH, successfully identify borosin α-*N*-methyltransferases and their surrounding BGCs. Unlike the fused NMT-precursor architectures observed in fungi, bacterially derived borosin BGCs encode discrete precursors and modifying enzymes as are more commonly seen with RiPPs. However, substantially different protein architectures are found in both NMTs and precursors in these split borosin pathways. NMTs greater than 1000 AAs and precursors in excess of 600 AAs with cores embedded in the middle of the protein are a subset of distinct structural features that have contributed to the diversification of split borosins into Types IV-X. This diversity in overall architecture as well as in core peptide sequence and methylation pattern, with examples of precursors methylated successively for >30 residues, beg the questions of what are the final metabolites, what are their biological functions, and why have they remained elusive despite their widespread distribution? We are passionately pursuing these questions and many others surrounding these disparate α-*N*-methylating borosin RiPP pathways. This work continues to highlight the advantages of genomics-driven approaches to natural product BGC discovery.

## Supporting Information

Supporting Information: Additional experimental details, materials and methods, LC-MS/MS spectra, and additional clustering analyses

Supplementary Dataset S1: Primers, genes, and protein sequences used in this study

Supplementary Dataset S2: Protein sequences used for the SSN in Figure 3

Supplementary Dataset S3: Global BGC Pfam table with organism taxonomic data from BiG-SCAPE analysis

Supplementary Dataset S4: BGC-family-specific Pfam tables from BiG-SCAPE analysis related to Figure S8

## Acknowledgment

We would like to thank L. Amoureux for strain *A. insuavis* AXX-A. We are grateful to M. Jensen and F. Miller for initial work with RceA & RceM as well as M. Quijano with AinM. We thank M. Medema for helpful discussions concerning the work in this manuscript.

