## Supporting Information for "Diverse Protein Architectures and α-*N*-Methylation Patterns Define Split Borosin RiPP Biosynthetic Gene Clusters"

### Contents


Materials and Methods S3

Fig. S1. LC-MS(/MS) data for 'split' borosin precursor *E. coli* expressions S6

Fig. S2. Location of RceA S78F mutation for LC-MS/MS analysis S43

Fig. S3. Size exclusion chromatography (SEC) of characterized NMTs S44

Fig. S4. AlphaFold2 protein models of NMTs and precursors S45

Fig. S5. Sequence similarity network of α-*N*-methyltransferase domains, cutoff $1 \times 10^{-60}$ S46

Fig. S6. Sequence similarity network of α-*N*-methyltransferase domains, cutoff $1 \times 10^{-90}$ S47

Fig. S7. Sequence similarity network of α-*N*-methyltransferase domains, cutoff $1 \times 10^{-80}$ S48

Fig. S8. BiG-SCAPE network of putative borosin BGCs, colored by taxonomic group S49

Fig. S9. BiG-SCAPE network of putative borosin BGCs, colored by borosin type S50

Fig. S10. Putative BGCs of characterized 'split' borosins S51

Fig S11. SDS-PAGE gel of purified α-*N*-methyltransferase and precursor proteins from *E. coli* S52

Supporting references S53

### Materials and Methods

#### Materials

HiFi DNA Assembly Master Mix, restriction enzymes, OneTaq and Q5 High Fidelity DNA polymerase were purchased from New England Biolabs (NEB). Gene synthesis and codon optimization was performed by Twist Biosciences (sequences found in Supplemental Dataset S1).

Commercial proteases were purchased from Promega (sequencing-grade trypsin, AspN, GluC and chymotrypsin) or Gold Biotechnology (proteinase K). Primers were ordered from Integrated DNA Technologies, and are listed in Supplemental Dataset S1. Unless otherwise stated, chemicals and reagents were purchased from Millipore Sigma.

*Achromobacter insuavis* AXX-A was obtained from Dr. Lucie Amoureux. *Rhodospirillum centenum* SW, *Pseudomonas mosselii* CIP 105259 and *Burkholderia stabilis* LMG 14294 were purchased from the American Type Culture Collection (ATCC).

#### Cloning and gene synthesis

Genes *rceA, rceM, ainA*, *ainM, pmoA*, and *pmoM* were directly cloned from genomic DNA of *R. centenum* SW (*rceA* and *rceM*), *A. insuavis* AXX-A (*ainA* and *ainM*), and *P. mosselii* (*pmoA* and *pmoM*). Genomic DNA was extracted after the bacteria were grown overnight in LB media. Genes were PCR amplified with Q5 High-Fidelity DNA Polymerase (NEB) using standard conditions (1x Q5 Reaction buffer, 1x Q5 High GC Enhancer buffer, 200 μM dNTPs, 0.5 μM final concentration of each primer, 0.04 U Q5 polymerase / 50-μL reaction) using primers listed in Supplemental Dataset S1, and the verified PCR products were excised, purified (Monarch DNA Gel Extraction Kit, NEB), and cloned into pET28a and/or pCDFDuet-1 via Gibson assembly (NEBuilder® HiFi DNA Assembly Master Mix, NEB) using manufacturer's protocols.

Gene synthesis for *sliA, sliM, surA, surM1* and *surM2* was performed by Twist Biosciences. Sequences were codon-optimized for expression in *E. coli*, and designed to include hexahistidine- or SUMO-tags.

All genes were cloned and expressed with N-terminal hexahistidine (N-His) tags or N-His SUMO tags; sequences are listed in Supplemental Dataset S1.

#### Protein expression and purification

Protein expression and purification was performed as described previously.[1] Briefly, genes were expressed in BL21(DE3) or LOBSTR (Kerafast) *E. coli* at 16 °C for 24 h, 48 h or 72 h. Cells were harvested and lysed using sonication. Recombinant proteins were purified via nickel-affinity chromatography based on manufacturer's recommendations (Ni-NTA resin, Gold Biotechnology). An SDS-PAGE gel of all proteins characterized in this work is included in Figure S11.

**Proteolytic digestion**

Proteolytic digestion was performed as previously described.[1] Briefly, Trypsin Gold, chymotrypsin, AspN, GluC (Promega) or Proteinase K (Gold Biotechnology) was used with an in-gel digestion method. Bands from soluble fractions were excised from SDS-PAGE gels stained with Coomassie Blue and cut into ~2 mm x 2 mm cubes. Gel pieces were transferred to 1.5 mL Protein LoBind tubes (Eppendorf) and then washed with a 1:1 ratio of 100 mM ammonium bicarbonate (ABC) : acetonitrile (ACN) three times until all the stain was removed. Gel pieces were then dehydrated in 100% ACN until semi-opaque (~30 sec), after which the ACN was discarded. If the putative RiPP core sequences contained cysteines, reduction (treatment with 10 mM DTT in a 65 °C water bath for 1 hr before discarding DTT solution) and alkylation (treatment with 55 mM iodoacetamide in 50 mM ABC at room temperature for 30 min) were performed. After reduction and alkylation, gel pieces were washed twice using a 1:1 ratio of ABC:ACN and then dehydrated in 100% ACN until semi-opaque as in the previous steps. Gel pieces were then rehydrated in digestion buffer (50 mM ABC, 5 mM $CaCl_2$, and appropriate units of protease) on ice for 15 min before overnight incubation at 37 °C or 25 °C (chymotrypsin only). Trypsin digests were performed in a 1:40 protease:protein ratio, whereas proteinase K digests were performed using a ~1:4-1:6 ratio. Digestion supernatants were recovered the next day, and placed in a fresh LoBind tube. Cleaved peptides were recovered by dehydrating the gel pieces in three successive steps. First, 60 µL of 50% ACN and 0.3% formic acid (FA) was added, incubated for 15 min at room temperature and recovered. Second, 60 µL of 80% ACN and 0.3% FA was added, incubated and recovered. Finally, 60 uL of 95% ACN and 0.1% FA was added, incubated, and recovered. The extracted peptides were pooled and frozen at -80 °C for 30 min to deactivate the protease. Peptide solutions were then thawed and dried using a Vacufuge Concentrator (Eppendorf). Peptides were resuspended in 0.1% FA and further purified and desalted using C18 ZipTips (Millipore Sigma) according to the manufacturer's specifications. After drying the samples again, peptides were resuspended in 15-30 µl of 20% ACN, 0.1% FA, and transferred to glass vials for MS analysis.

**Peptide mass spectrometric analysis (LC-MS/MS HCD)**

LC-MS/MS analysis of digested peptides was performed as described previously.[1] Briefly, data were recorded on a Thermo Scientific Fusion or Fusion Lumos mass spectrometer equipped with a Dionex Ultimate 3000 UHPLC system using a nLC column (200 mm × 75 µm) packed using Vydac 5-µm particles with a 300 Å pore size (Hichrom Limited). Elution was performed with a linear gradient using water with 0.1% FA (solvent A) and ACN with 0.1% FA (solvent B) at a flow rate of 0.3 µl/min. The column was equilibrated with 20% solvent B for 5 min, followed by a linear increase of solvent B to 85% over 32 min and a final elution step with 85% solvent B for 2 min. Mass spectra were acquired in positive-ion mode. Full MS was done at a resolution of 60,000 [automatic gain control (AGC) target, $4 \times 10^5$; maximum injection time (IT), 50-100 ms; range, 300 to 1800 m/z], and data-dependent as well as targeted MS/MS was performed at a resolution of 15,000 (AGC target, $5 \times 10^5$; maximum IT, 100 - 500 ms; isolation window, 2.2 m/z) using higher-energy collisional dissociation (HCD) energies from 14-25% with steps of ±4% were used. Data were processed using Thermo Fisher Xcalibur software and MaxQuant v1.6.10.

#### AlphaFold2 protein models

Version 2.1.1 of AlphaFold2[2] was accessed through the Minnesota Supercomputing Institute (MSI). FASTA files containing the amino acid sequences of the α-*N*-methyltransferases and precursor proteins were used as input query with default prediction parameters. Alphafold2 through MSI uses a non-docker implementation and a bash script to run in a Linux environment. Output files were loaded into PyMOL (Schrödinger, Inc.) Models were colored based on a per-residue confidence score (pLDDT) stored within the b-factor putty preset. The sequence alignment quality graphs were captured from the output using ColabFold: AlphaFold2 using MMseqs2.

#### Creating the sequence similarity networks

A BLASTP[3] search of the non-redundant protein database using residues 1-250 of the OphMA sequence was conducted using default settings (April 2021). The search returned 1804 protein sequences. Sequences that did not contain a full methyltransferase region (corresponding to residues -250 of OphMA) were removed. An all-vs-all BLASTP analysis under default settings was performed on the remaining 1,704 sequences in addition to the amino acid sequences of 55 fungal borosins and 76 representative sequences from the YabN-like protein superfamily (Supplementary Dataset 2). The all-vs-all similarity scores were visualized in Cytoscape v3.6.1[4] with various E-value cutoff values. For global alignment, all-vs-all analysis was performed using the needleall module in the Emboss suite[5] with a gap penalty of -10, an extension penalty of -0.5, and an endgap penalty of -10. Scores were normalized by alignment length and then visualized in Cytoscape[4] using various cutoff values. Representative sequences were identified using MMseqs2 v13.45111[6] with a minimum identity of 90% using cluster mode (0). Non-representative sequences were then removed from the networks.

#### Identification of bacterial and archaeal borosin BGCs

A profile hidden Markov model (HMM) was built from an alignment of 137 high-confidence fungal and bacterial borosin methyltransferase domains using HMMER version 3.3.1 (http://hmmer.org). This profile HMM was added to a locally installed version of antiSMASH v5.2.0[7] and used to parse genome assemblies from the *Burkholderia* genus (taxid 32008, accessed October, 2020), the cyanobacteria clade (taxid 1798711, accessed November, 2020), the Streptomycetales order (taxid 85011, accessed November, 2020), and the Alteromonadales order (taxid 135622, accessed January, 2021) available on NCBI (10,868 assemblies). Selection of these groups was based on a preliminary BLASTP search showing these groups as the predominant sources of homologs to the *N*-methyltransferase domain of OphMA. A subset of biosynthetic gene cluster (BGC) regions containing the borosin profile HMM was pulled from the antiSMASH output. From this subset, protein sequences satisfying antiSMASH's "TP_methylase" rule were extracted, aligned using Clustal W 2.1,[8] then manually inspected. Low-quality sequences, sequences with high variance from previously identified conserved regions, as well as redundant sequences were discarded, leaving 460 sequences.[1] These sequences were appended to the 137 sequences used in the first profile HMM. 574 sequences remained after redundant sequences were removed. A profile HMM, borosinMT.hmm, was built from the methyltransferase domain (corresponding to residues 1-250 of OphMA) of this newly curated protein alignment. In addition, profile HMMs were built from dereplicated alignments of regions containing the highly conserved and distinctive YGHP and DCLFAD amino acid motifs (YGHP.hmm and DCLFAD.hmm, respectively) (Figure 3). A detection rule, BorosinMT, for a coding DNA sequence (CDS) region containing MTdomain.hmm and either YGHP.hmm or DCLFAD.hmm with a cutoff distance of 20 kb and a neighborhood distance of 10 kb was added to antiSMASH.[7] This rule was used to parse bacterial and archaeal genomes for putative borosin biosynthetic gene clusters in antiSMASH under default settings and using Prodigal for gene prediction.

**Domain analysis of putative borosin BGCs**

Genome assemblies in the refseq database linked to the protein accession IDs of the borosin α-*N*-methyltransferases identified through BLASTP were downloaded (1902 assemblies) and analyzed through antiSMASH[7] (containing the previously-described BorosinMT rule). Representative sequences were identified from the resulting 2789 clusters using the cluster function in MMseqs2[6] with a minimum identity of 90% with cluster mode (2) and coverage mode (1) (80% coverage). This resulted in 1055 BGC region sequences which were then run through BiG-SCAPE v1.1.0[9] for cluster analysis. Clusters with less than 5 predicted pfam domains or that did not contain a predicted TP_methylase pfam domain were removed from the dataset, leaving 1030 clusters (Supplemental Dataset 3). Network information was then imported into and visualized using Cytoscape. BGCs that clustered together with a raw distance cutoff value of 0.5 were assigned to the same BGC family (Supplemental Dataset 4). Heatmaps showing the most commonly occurring Pfam domains within cluster families were created using the heatmaply[10] in Rstudio v 1.2.5033.

**Fig. S1. LC-MS(/MS) data for borosin precursor *E. coli* expressions**. LC-MS and LC-MS/MS spectra for borosin precursor expressions reveal methylated residues in proteolytically released core peptide fragments. The borosin precursor, time of in-vivo expression, parent ion details, and LC retention times (RT) are denoted in the upper right-hand corner of the LC-MS/MS spectra. Observed MS/MS fragmented masses are listed above (b-ions) and below (y-ions) the listed sequence with grey lines marking sites of fragmentation. The mass difference from the theoretical expected masses are labelled in parentheses. A mass cutoff of 10.0-ppm was used for the annotated LC-MS/MS peaks. Ion masses are denoted with varying numbers of methylations in brackets, where 'Me' marks a mass shift corresponding to methylation. Please see the Materials and Methods section for more details concerning the acquisition of the LC-MS/MS data. **(a)** Organization of LC-MS(/MS) data for *E. coli* expressions of borosins. The table indicates figure panel numbers, summary of expression data presented in the panels, corresponding borosin precursor protein, and LC-MS(/MS) peptide sequences. Residues in the LC-MS fragments ('Sequence' column) are shaded orange based on verified or inferred α-*N*-methylation position. **(b-h)** LC-MS/MS data for RceA (or RceA S78F)/M expressions for 24 or 48 hrs. **(i-l)** LC-MS/MS data for SliA/M expressions for 24 hrs. **(m-r)** LC-MS/MS data for AinA/M expressions for 72 hrs. **(s-x)** LC-MS/MS data for PmoA/M expressions for 24 or 72 hrs. **(y-ac)** LC-MS/MS data for SurA/M1 expressions for 24 or 48 hrs. **(ad-af)** LC-MS/MS data for SurA/M2 expressions for 24 or 48 hrs. **(ag-aj)** LC-MS/MS data for SurA/M1/M2 expressions for 24 or 48 hrs.

| Fig. | Borosin Expression Data | Protein | Sequence |
|---|---|---|---|
| S1b | LC-MS/MS fragmentation data: 1 *N*-Me | RceA/M | DVAELSGGE**L**DVAELFGGEL |
| S1c | LC-MS/MS fragmentation data: 1 *N*-Me | RceA/M | DVIELSGGE**L**DVAELFGGEL |
| S1d | LC-MS/MS fragmentation data: 1 *N*-Me | RceA/M | SGGE**L**DVAEIGIINTFDL |
| S1e | LC-MS/MS fragmentation data: 2 *N*-Me | RceA/M | ELFGGE**L**DVAELSGGE**L** |
| S1f | LC-MS/MS fragmentation data: 2 *N*-Me | RceA/M | DVIELSGGE**L**DVAELSGGE**L** |
| S1g | LC-MS/MS fragmentation data: 4 *N*-Me | RceA S78F/M | GGE**L**DVAELSGGE**L**DVAELSGGE**L** DVAELSGGE**L**DVAELF |
| S1h | LC-MS/MS fragmentation data: 4 *N*-Me | RceA S78F/M | GGE**L**DVAELSGGE**L**DVAELSGGE**L** DVAELSGGE**L**DVAEIGIINTF |
| S1i | LC-MS/MS fragmentation data: 7 *N*-Me | SliA/M | **TEVT**EV**V**EVV**EV**VE |
| S1j | LC-MS/MS fragmentation data: 11 *N*-Me | SliA/M | ATPEPPT**ITAVVAIVINS**STTEVA |
| S1k | LC-MS/MS fragmentation data: 12 *N*-Me | SliA/M | **EVAEVTEVTEVT**EVT |
| S1l | LC-MS/MS fragmentation data: 16 *N*-Me | SliA/M | **TAVVAIVINSSTTEVA**EVT |
| S1m | LC-MS/MS fragmentation data: 1 *N*-Me | AinA/M | EPGI**D**FVTG |
| S1n | LC-MS/MS fragmentation data: 3 *N*-Me | AinA/M | VDVDV**D**V**D**V**T**LE |
| S1o | LC-MS/MS fragmentation data: 4 *N*-Me | AinA/M | VDVDV**D**V**D**V**TL**EM |
| S1p | LC-MS/MS fragmentation data: 4 *N*-Me | AinA/M | VEIPDDPNV**D**V**D**V**D**A**D**VEAD |

| | | | |
|---|---|---|---|
| S1q | LC-MS/MS fragmentation data: 5 *N*-Me | AinA/M | VEIPDDPNVDVDVDADVEA |
| S1r | LC-MS/MS fragmentation data: 6 *N*-Me | AinA/M | TEVDVDVDVDVTLEM |
| S1s | LC-MS/MS fragmentation data: 5 N-Me | PmoA/M | IAVVDVDVDVDIDVD |
| S1t | LC-MS/MS fragmentation data: 5 *N*-Me | PmoA/M | DIDVDVDIDTDVN |
| S1u | LC-MS/MS fragmentation data: 6 N-Me | PmoA/M | ADPVNVDVDIDVDIDVDVD |
| S1v | LC-MS/MS fragmentation data: 6 *N*-Me | PmoA/M | IAVVDVDVDVDIDVDID |
| S1w | LC-MS/MS fragmentation data: 7 *N*-Me | PmoA/M | IAVVDVDVDVDIDVDID |
| S1x | LC-MS/MS fragmentation data: 8 *N*-Me | PmoA/M | TDTETDIDIDVDVDIDTDVN |
| S1y | LC-MS/MS fragmentation data: 1 *N*-Me | SurA/M1 | LMAEILFKAWSDDDFRE |
| S1z | LC-MS/MS fragmentation data: 1 *N*-Me | SurA/M1 | AGGPRTLLRPGNDEIPVVEAQGSE |
| S1aa | LC-MS/MS fragmentation data: 1 *N*-Me | SurA/M1 | GSHMATSALTNLLTEISEDPAR |
| S1ab | LC-MS/MS fragmentation data: 1 *N*-Me | SurA/M1 | LTEWLEDPDSYMNR |
| S1ac | LC-MS/MS fragmentation data: 1 *N*-Me | SurA/M1 | LREIAEAGGPR |
| S1ad | LC-MS/MS fragmentation data: 1 *N*-Me | SurA/M2 | ILFKAWSDDDFRE |
| S1ae | LC-MS/MS fragmentation data: 1 *N*-Me | SurA/M2 | LTEWLEDPDSYMNR |
| S1af | LC-MS/MS fragmentation data: 1 *N*-Me | SurA/M2 | HIVIPWLPPDVEEETR |
| S1ag | LC-MS/MS fragmentation data: 1 *N*-Me | SurA/M1/M2 | SELMAEILFK |
| S1ah | LC-MS/MS fragmentation data: 1 *N*-Me | SurA/M1/M2 | AGLTEEETR |
| S1ai | LC-MS/MS fragmentation data: 2 *N*-Me | SurA/M1/M2 | AGLTEEETR |
| S1aj | LC-MS/MS fragmentation data: 2 *N*-Me | SurA/M1/M2 | HIVIPWLPPDVEEETR |

a

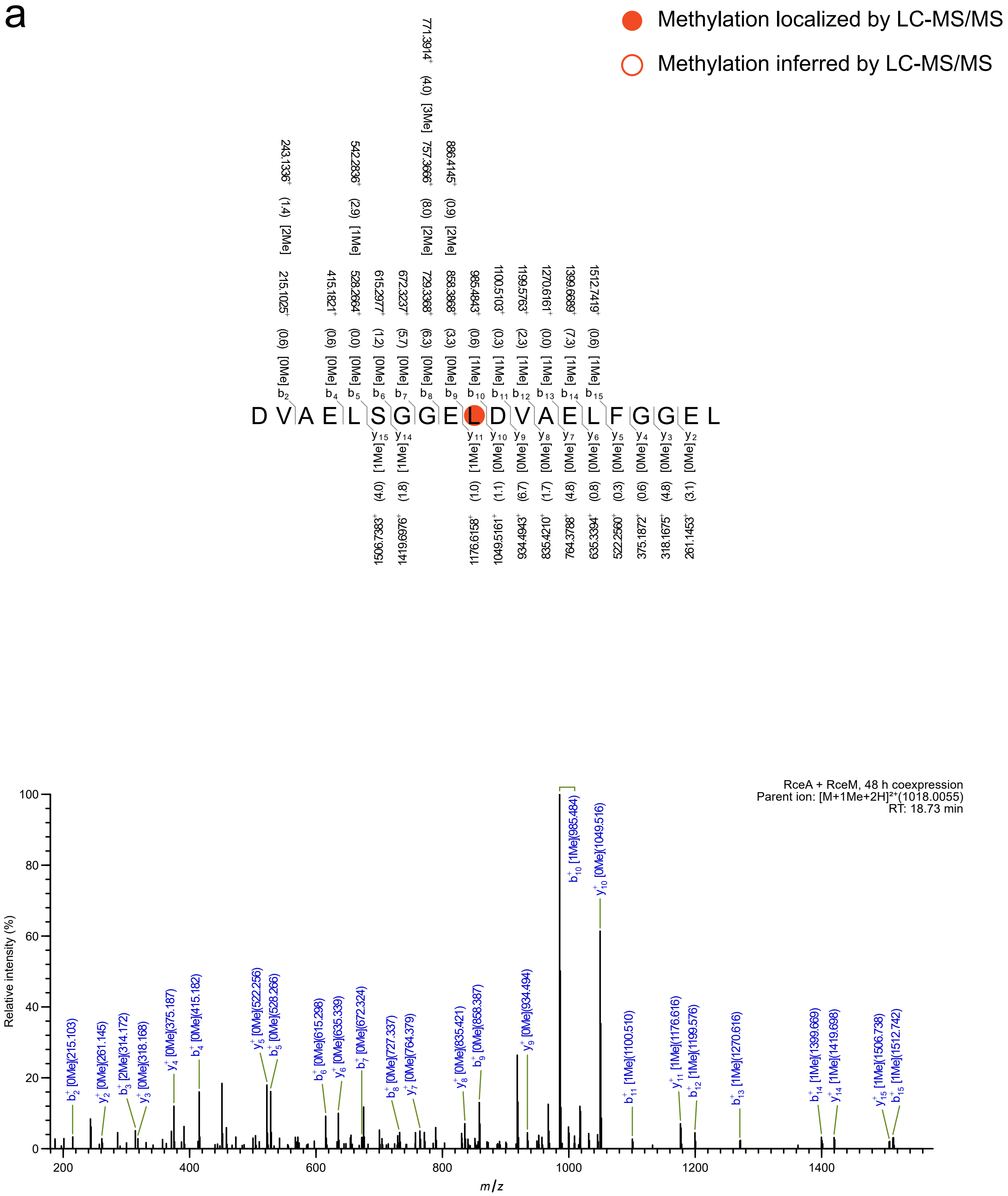
Methylation localized by LC-MS/MS
Methylation inferred by LC-MS/MS
D V A E L S G G E L D V A E L F G G E L
RceA + RceM, 48 h coexpression
Parent ion: [M+1Me+2H]²⁺(1018.0055)
RT: 18.73 min
Relative intensity (%)
m/z

b

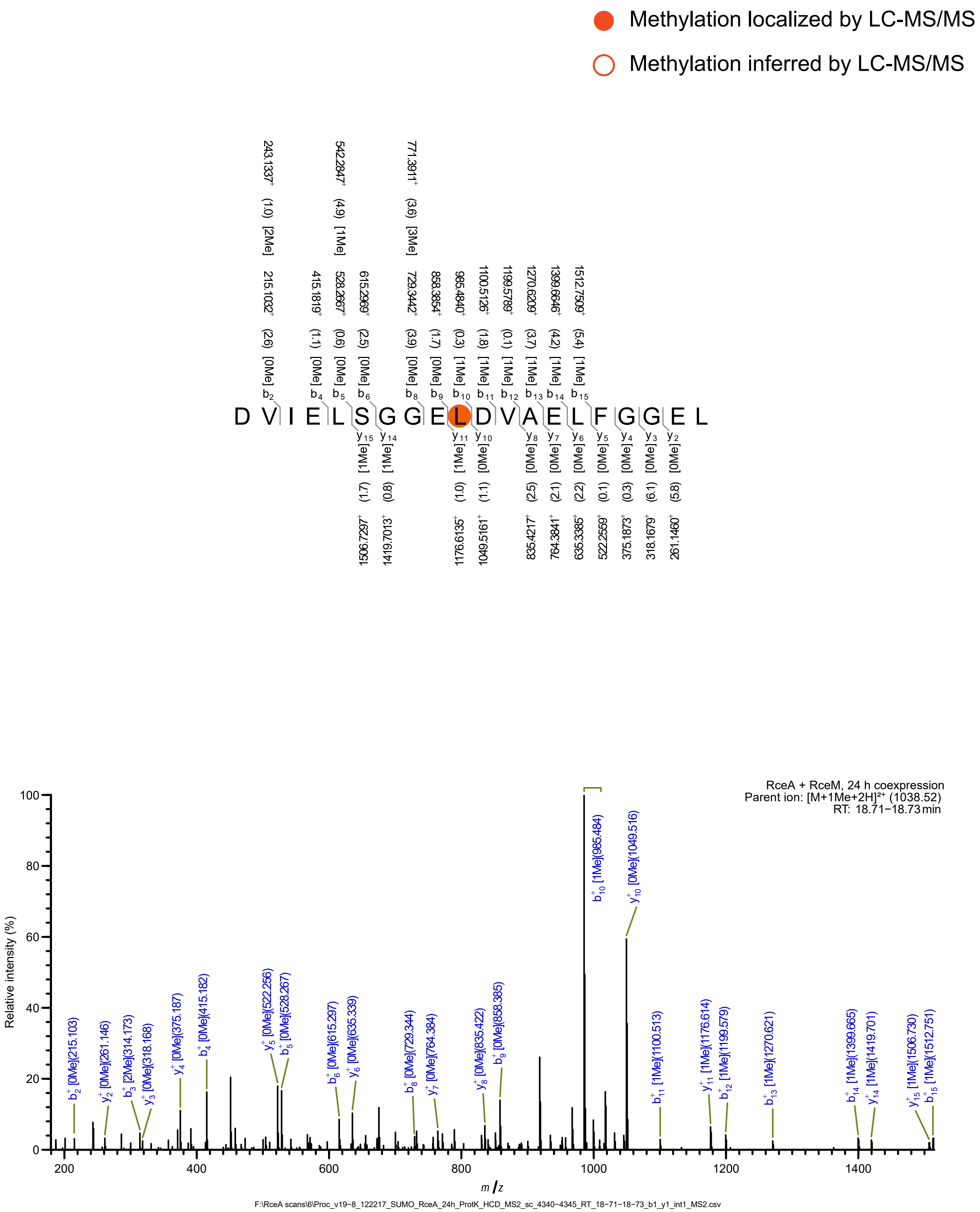

Methylation localized by LC-MS/MS
Methylation inferred by LC-MS/MS
243.1337+ (1.0) [2Me]
542.2847+ (4.9) [1Me]
771.3911+ (3.6) [3Me]
215.1032+ (2.6) [0Me] b2
415.1819+ (1.1) [0Me] b4
528.2667+ (0.6) [0Me] b5
615.2969+ (2.5) [0Me] b6
729.3442+ (3.9) [0Me] b8
858.3854+ (1.7) [0Me] b9
985.4840+ (0.3) [1Me] b10
1100.5126+ (1.8) [1Me] b11
1199.5789+ (0.1) [1Me] b12
1270.6209+ (3.7) [1Me] b13
1399.6646+ (4.2) [1Me] b14
1512.7509+ (5.4) [1Me] b15
D V I E L S G G E L D V A E L F G G E L
y15 [1Me] (1.7) 1506.7297+
y14 [1Me] (0.8) 1419.7013+
y11 [1Me] (1.0) 1176.6135+
y10 [0Me] (1.1) 1049.5161+
y8 [0Me] (2.5) 835.4217+
y7 [0Me] (2.1) 764.3841+
y6 [0Me] (2.2) 635.3385+
y5 [0Me] (0.1) 522.2559+
y4 [0Me] (0.3) 375.1873+
y3 [0Me] (6.1) 318.1679+
y2 [0Me] (5.8) 261.1460+
RceA + RceM, 24 h coexpression
Parent ion: [M+1Me+2H]²⁺ (1038.52)
RT: 18.71–18.73 min
Relative intensity (%)
m/z
b2+ [0Me](215.103)
y2+ [0Me](261.146)
b3+ [2Me](314.173)
y3+ [0Me](318.168)
y4+ [0Me](375.187)
b4+ [0Me](415.182)
y5+ [0Me](522.256)
b5+ [0Me](528.267)
b6+ [0Me](615.297)
y6+ [0Me](635.339)
b8+ [0Me](729.344)
y7+ [0Me](764.384)
y8+ [0Me](835.422)
b9+ [0Me](858.385)
b10+ [1Me](985.484)
y10+ [0Me](1049.516)
b11+ [1Me](1100.513)
y11+ [1Me](1176.614)
b12+ [1Me](1199.579)
b13+ [1Me](1270.621)
b14+ [1Me](1399.665)
y14+ [1Me](1419.701)
y15+ [1Me](1506.730)
b15+ [1Me](1512.751)
F:\RceA scans\6\Proc_v19-8_122217_SUMO_RceA_24h_ProtK_HCD_MS2_sc_4340-4345_RT_18-71-18-73_b1_y1_int1_MS2.csv

C

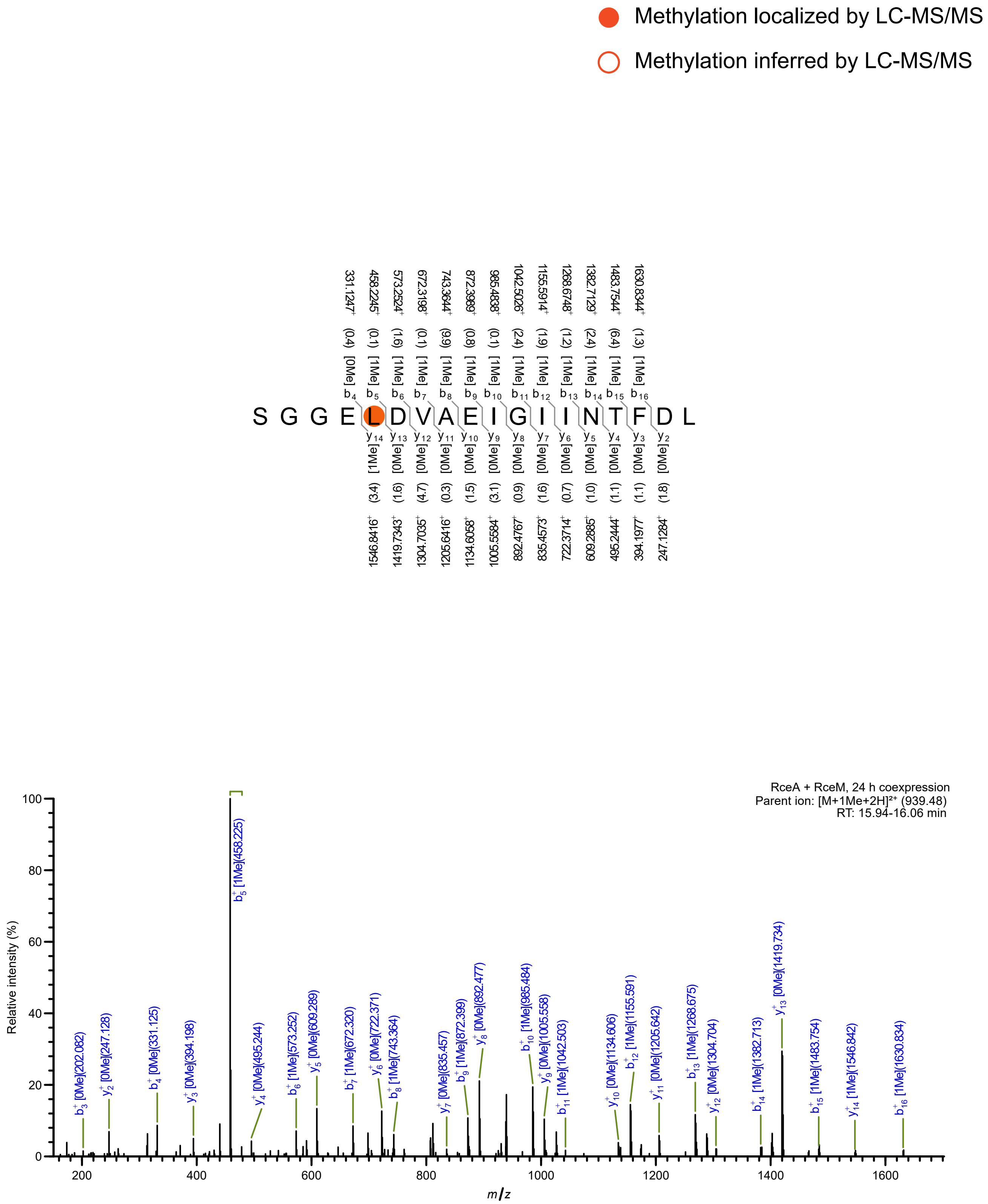

Methylation localized by LC-MS/MS
Methylation inferred by LC-MS/MS
S G G E L D V A E I G I I N T F D L
RceA + RceM, 24 h coexpression
Parent ion: [M+1Me+2H]²⁺ (939.48)
RT: 15.94-16.06 min
Relative intensity (%)
m/z

d

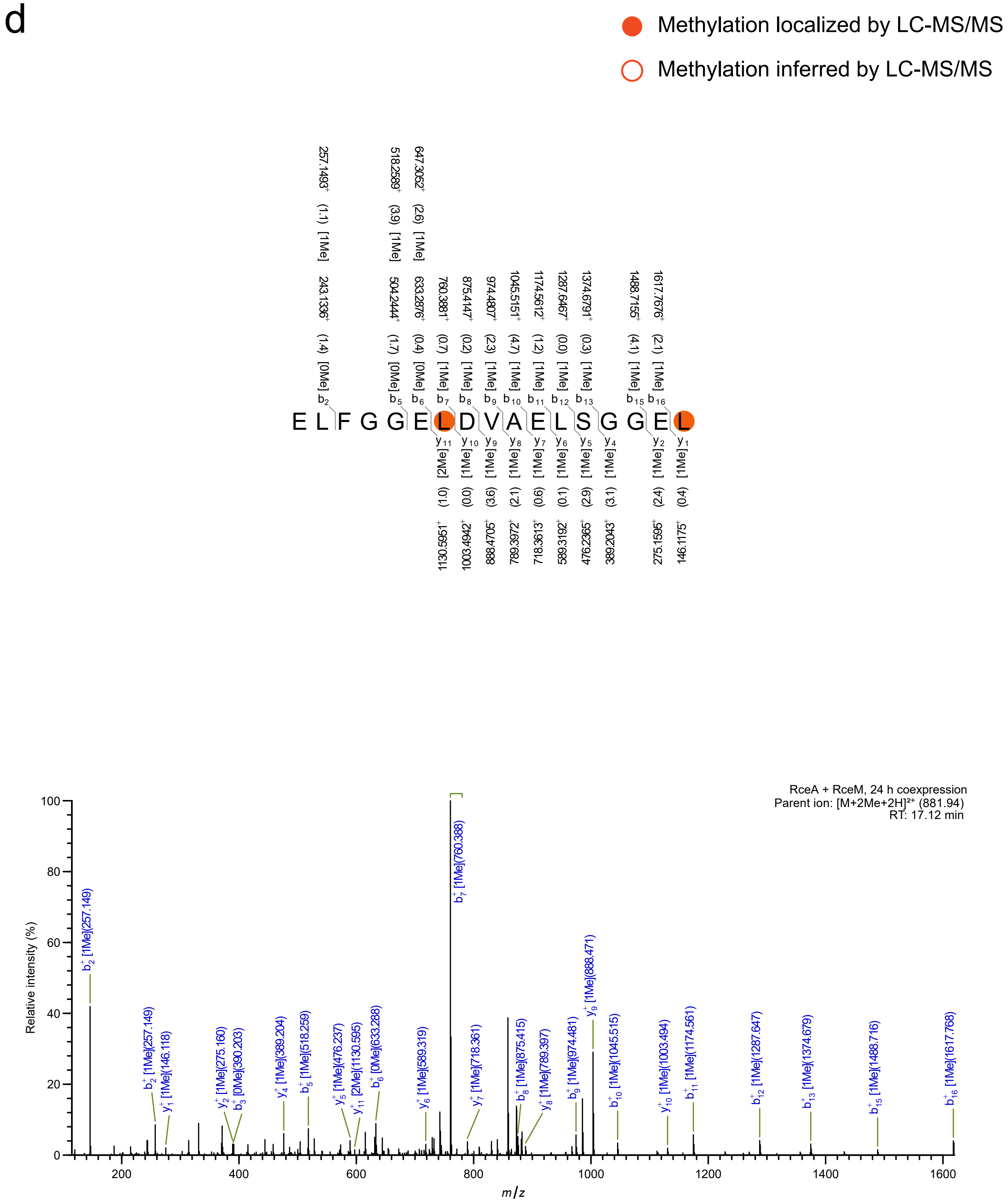

Methylation localized by LC-MS/MS
Methylation inferred by LC-MS/MS
E L F G G E L D V A E L S G G E L
RceA + RceM, 24 h coexpression
Parent ion: [M+2Me+2H]²⁺ (881.94)
RT: 17.12 min
Relative intensity (%)
m/z

e

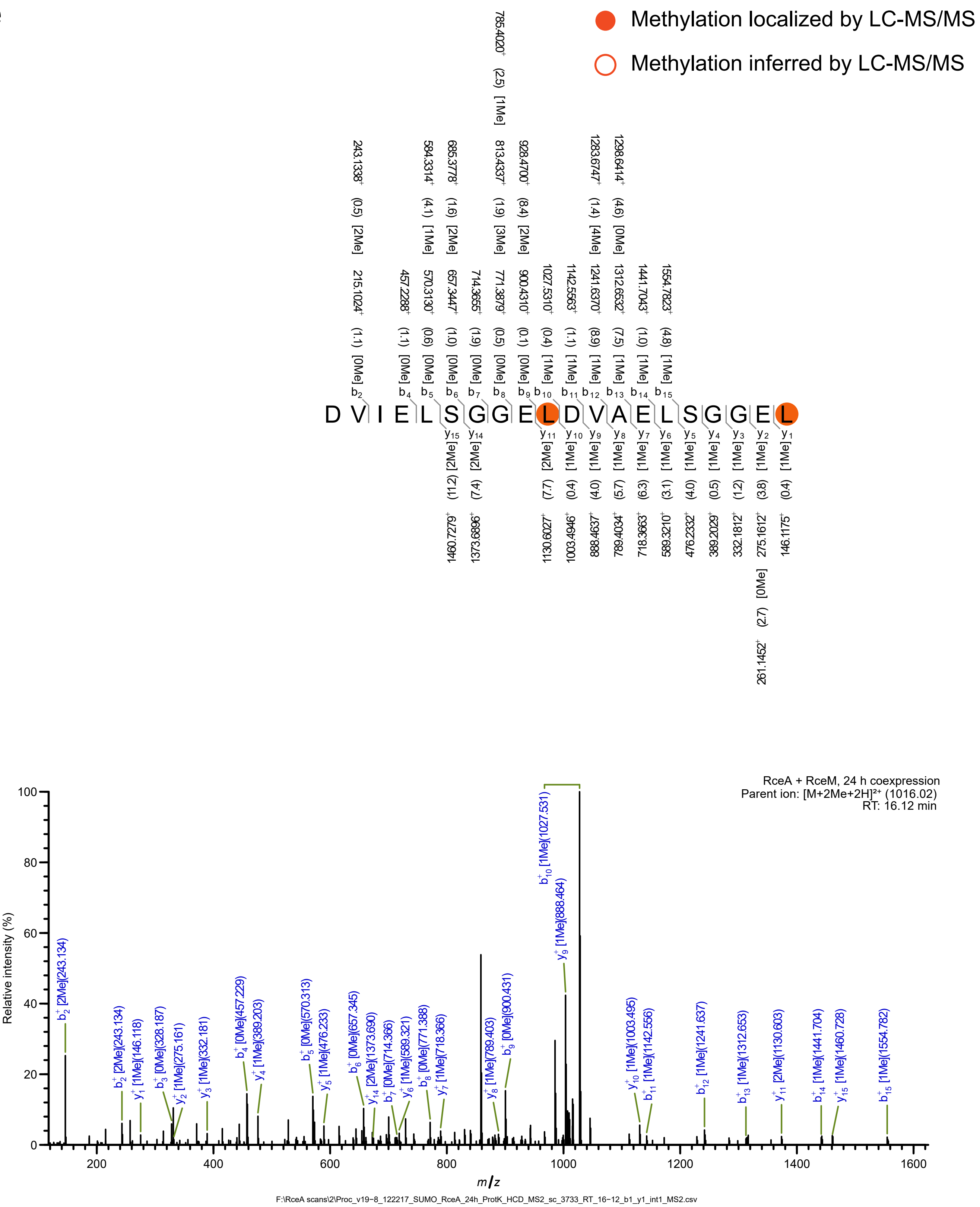

Methylation localized by LC-MS/MS
Methylation inferred by LC-MS/MS
D V I E L S G G E L D V A E L S G G E L
RceA + RceM, 24 h coexpression
Parent ion: [M+2Me+2H]²⁺ (1016.02)
RT: 16.12 min
Relative intensity (%)
m/z
F:\RceA scans\2\Proc_v19-8_122217_SUMO_RceA_24h_ProtK_HCD_MS2_sc_3733_RT_16-12_b1_y1_int1_MS2.csv

f

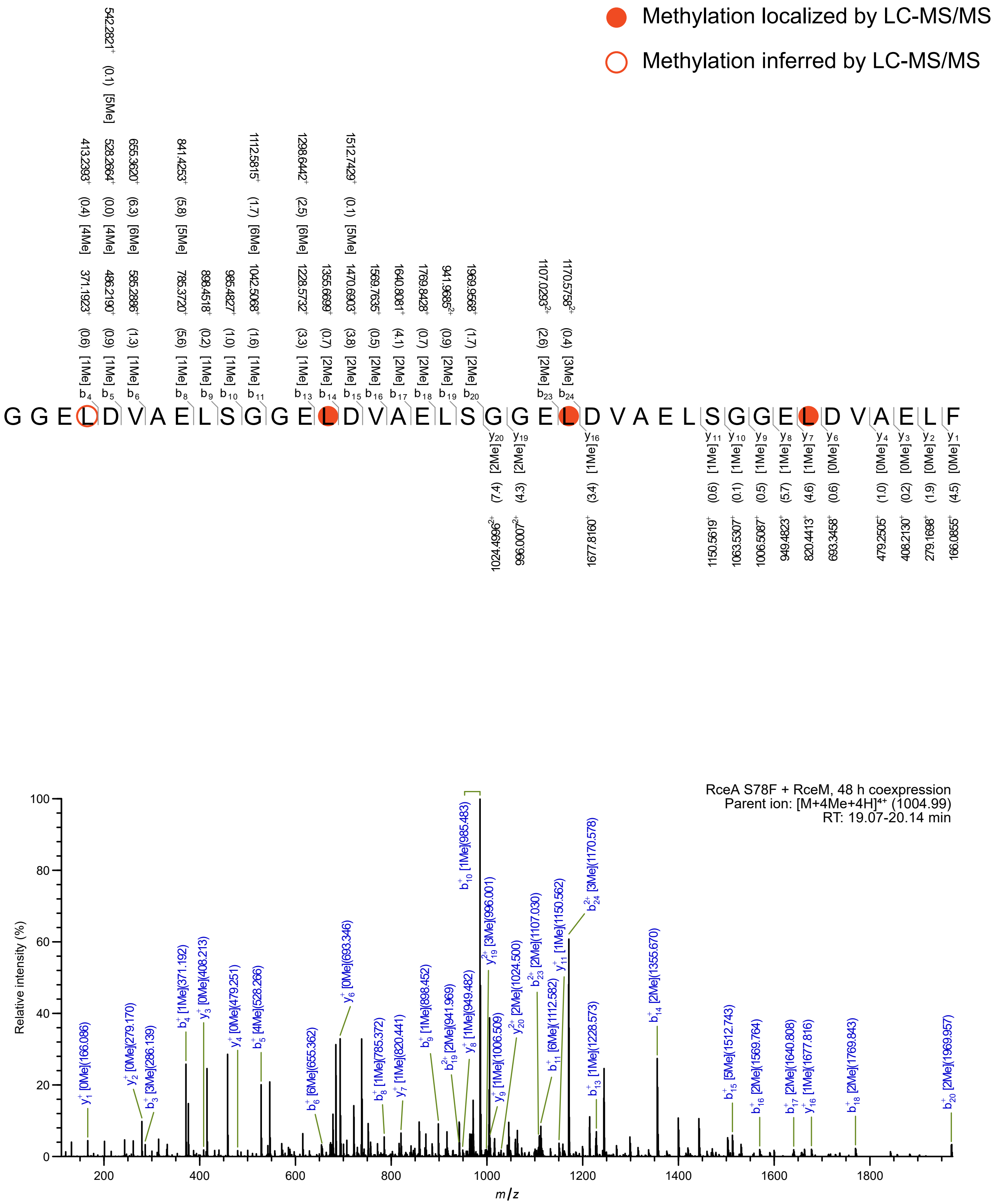

Methylation localized by LC-MS/MS
Methylation inferred by LC-MS/MS
GGELDVAELSGGELDVAELSGGELDVAELSGGELDVAELF
RceA S78F + RceM, 48 h coexpression
Parent ion: [M+4Me+4H]4+ (1004.99)
RT: 19.07-20.14 min
Relative intensity (%)
m/z

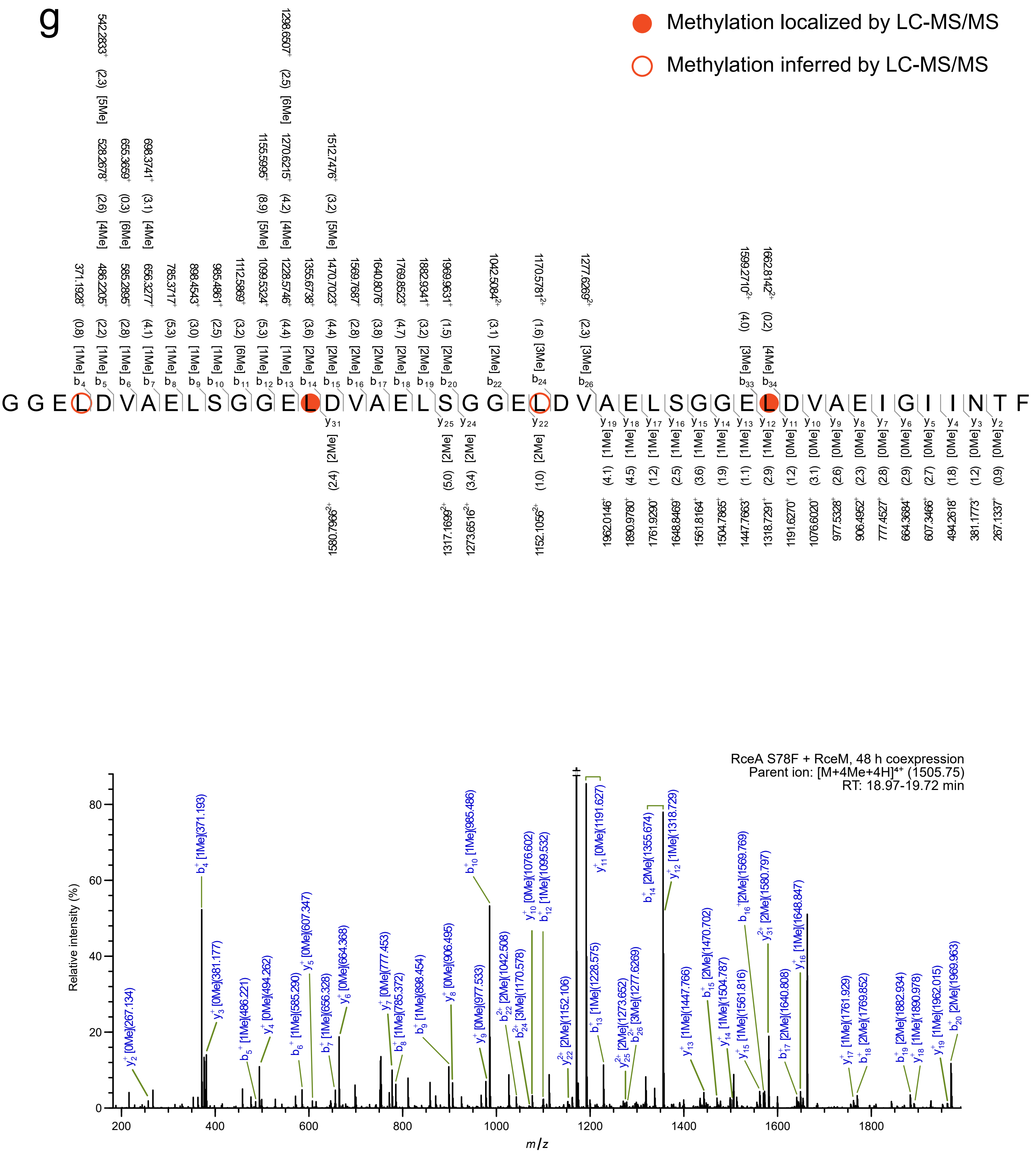
g
Methylation localized by LC-MS/MS
Methylation inferred by LC-MS/MS
GGELDVAELSGGELDVAELSGGELDVAELSGGELDVAEIGIINTF
RceA S78F + RceM, 48 h coexpression
Parent ion: [M+4Me+4H]4+ (1505.75)
RT: 18.97-19.72 min
Relative intensity (%)
m/z

h

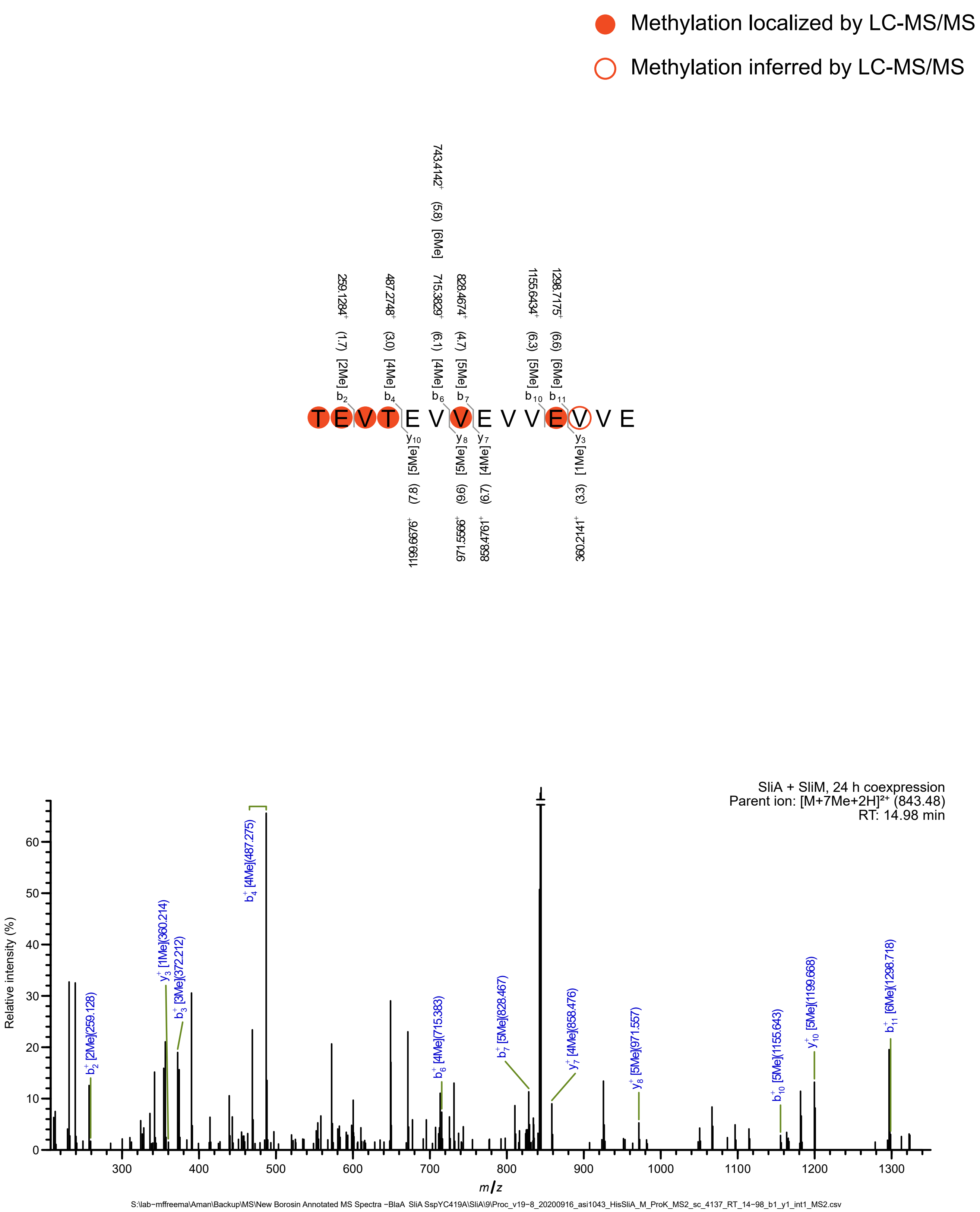

Methylation localized by LC-MS/MS
Methylation inferred by LC-MS/MS
T E V T E V V E V V E V V E
b2 259.1284+ (1.7) [2Me]
b4 487.2748+ (3.0) [4Me]
b6 715.3829+ (6.1) [4Me]
743.4142+ (5.8) [6Me]
b7 828.4674+ (4.7) [5Me]
b10 1155.6434+ (6.3) [5Me]
b11 1298.7175+ (6.6) [6Me]
y10 [5Me] (7.8) 1199.6676+
y8 [5Me] (9.6) 971.5566+
y7 [4Me] (6.7) 858.4761+
y3 [1Me] (3.3) 360.2141+
SliA + SliM, 24 h coexpression
Parent ion: [M+7Me+2H]²⁺ (843.48)
RT: 14.98 min
Relative intensity (%)
m/z
b2+ [2Me](259.128)
y3+ [1Me](360.214)
b3+ [3Me](372.212)
b4+ [4Me](487.275)
b6+ [4Me](715.383)
b7+ [5Me](828.467)
y7+ [4Me](858.476)
y8+ [5Me](971.557)
b10+ [5Me](1155.643)
y10+ [5Me](1199.668)
b11+ [6Me](1298.718)
S:\lab-mffreema\Aman\Backup\MS\New Borosin Annotated MS Spectra -BlaA SliA SspYC419A\SliA\9\Proc_v19-8_20200916_asi1043_HisSliA_M_ProK_MS2_sc_4137_RT_14-98_b1_y1_int1_MS2.csv

i

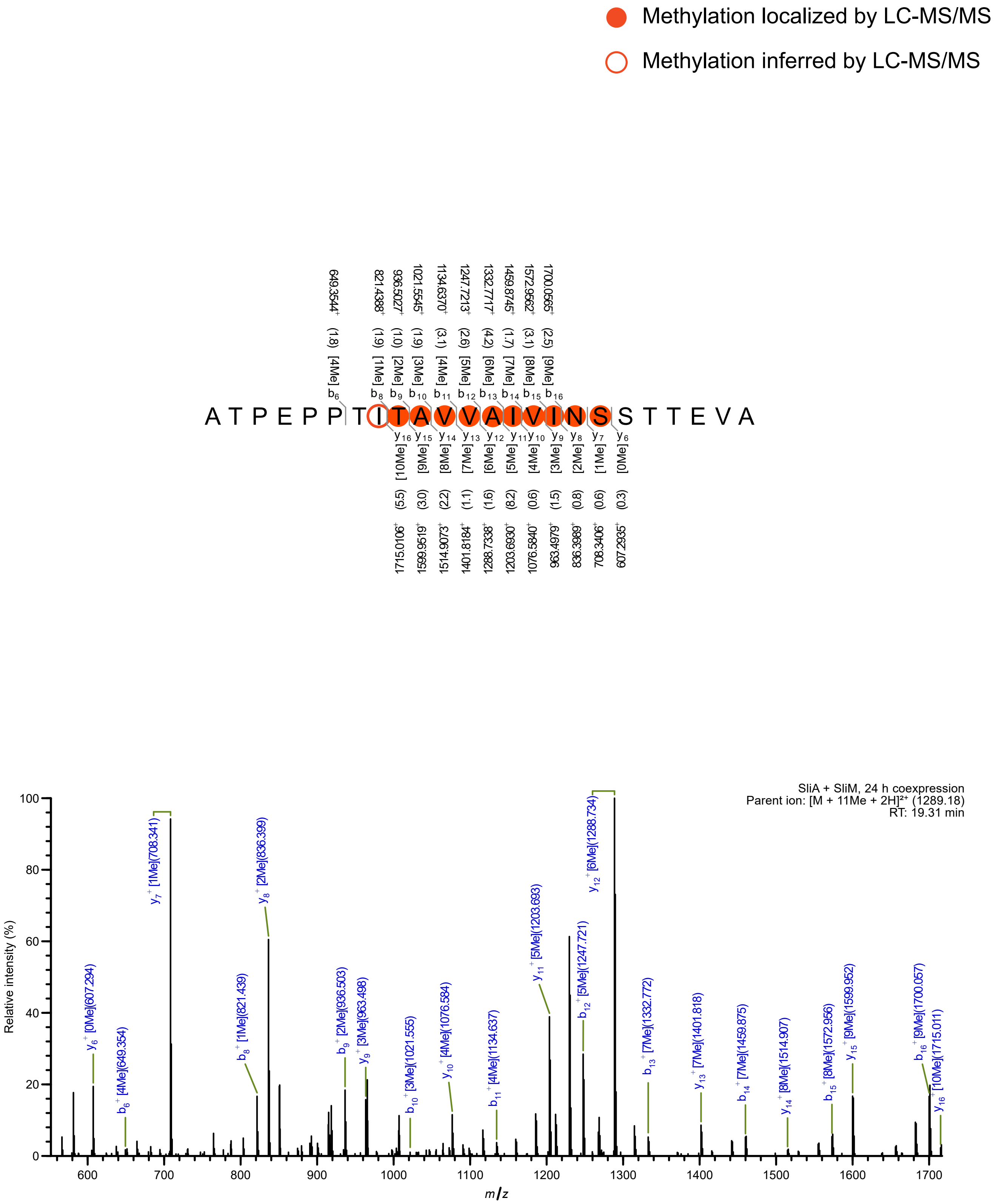

Methylation localized by LC-MS/MS
Methylation inferred by LC-MS/MS
A T P E P P T I T A V V A I V I N S S T T E V A
SliA + SliM, 24 h coexpression
Parent ion: [M + 11Me + 2H]²⁺ (1289.18)
RT: 19.31 min
Relative intensity (%)
m/z

j

Methylation localized by LC-MS/MS

Methylation inferred by LC-MS/MS

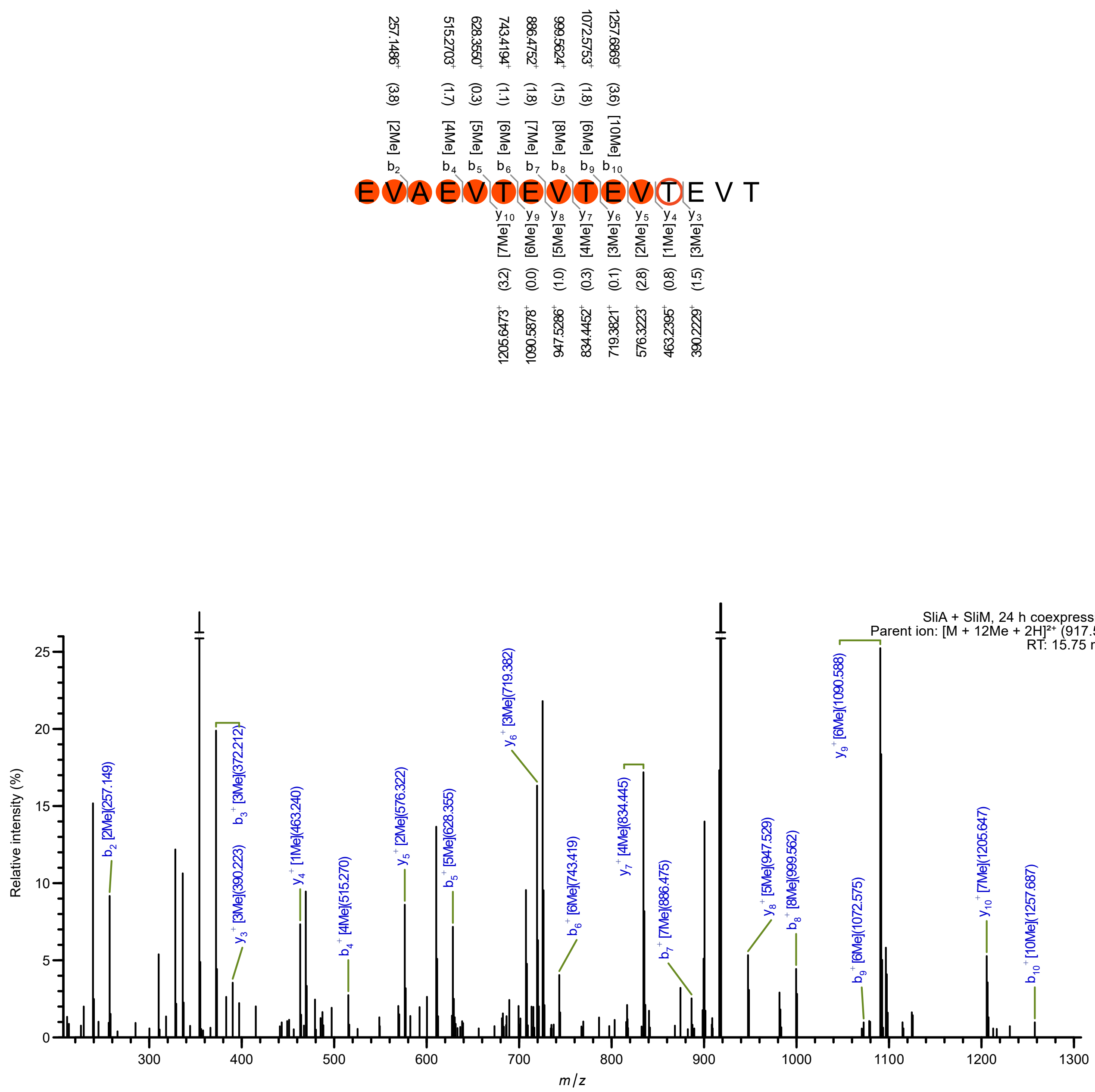

k

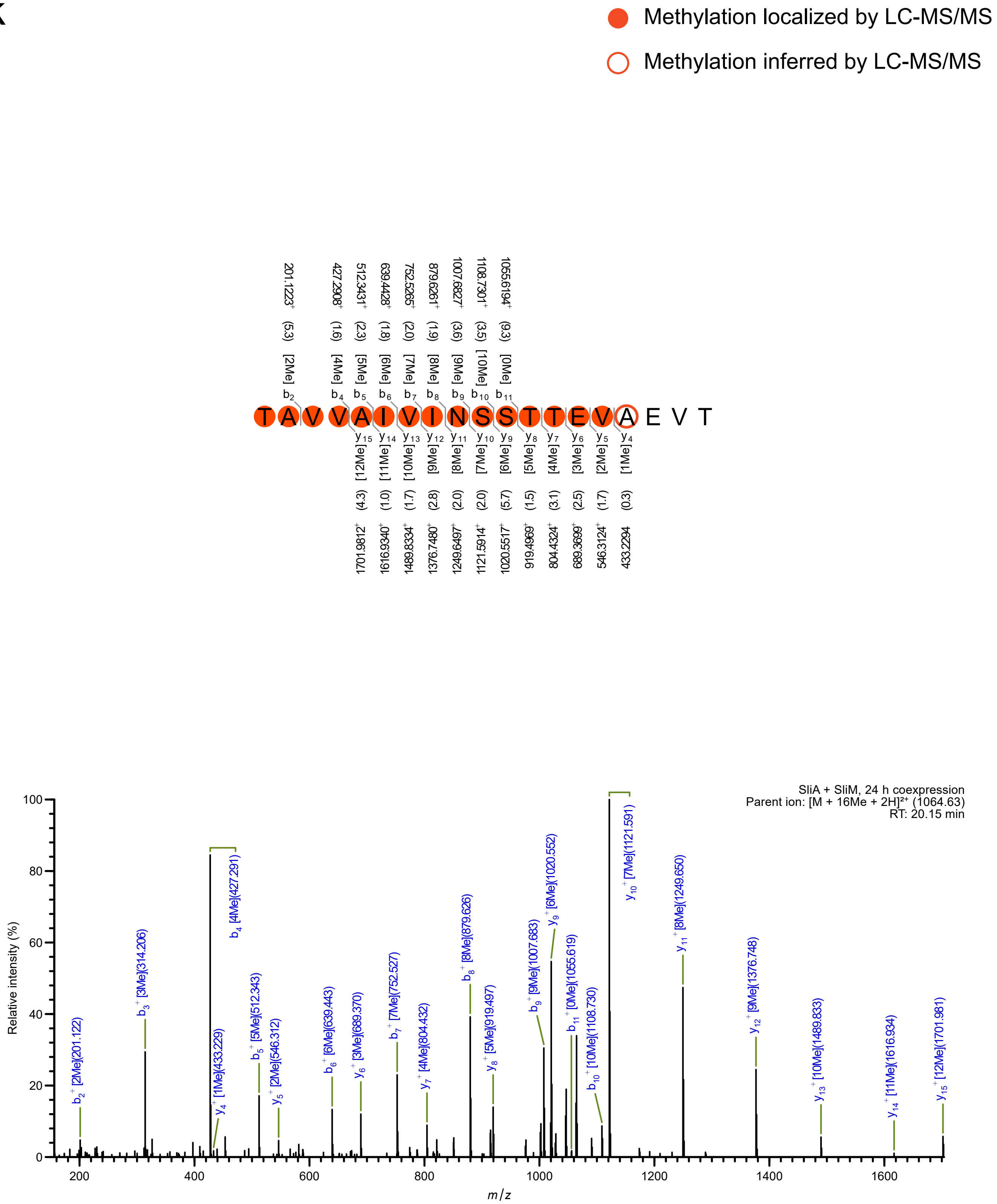

Methylation localized by LC-MS/MS
Methylation inferred by LC-MS/MS
T A V V A I V I N S S T T E V A E V T
b2 [2Me] (5.3) 201.1223+
b4 [4Me] (1.6) 427.2908+
b5 [5Me] (2.3) 512.3431+
b6 [6Me] (1.8) 639.4428+
b7 [7Me] (2.0) 752.5265+
b8 [8Me] (1.9) 879.6261+
b9 [9Me] (3.6) 1007.6827+
b10 [10Me] (3.5) 1108.7301+
b11 [0Me] (9.3) 1055.6194+
y15 [12Me] (4.3) 1701.9812+
y14 [11Me] (1.0) 1616.9340+
y13 [10Me] (1.7) 1489.8334+
y12 [9Me] (2.8) 1376.7480+
y11 [8Me] (2.0) 1249.6497+
y10 [7Me] (2.0) 1121.5914+
y9 [6Me] (5.7) 1020.5517+
y8 [5Me] (1.5) 919.4969+
y7 [4Me] (3.1) 804.4324+
y6 [3Me] (2.5) 689.3699+
y5 [2Me] (1.7) 546.3124+
y4 [1Me] (0.3) 433.2294
SliA + SliM, 24 h coexpression
Parent ion: [M + 16Me + 2H]2+ (1064.63)
RT: 20.15 min
Relative intensity (%)
m/z
b2+ [2Me](201.122)
b3+ [3Me](314.206)
y4+ [1Me](433.229)
b4+ [4Me](427.291)
b5+ [5Me](512.343)
y5+ [2Me](546.312)
b6+ [6Me](639.443)
y6+ [3Me](689.370)
b7+ [7Me](752.527)
y7+ [4Me](804.432)
b8+ [8Me](879.626)
y8+ [5Me](919.497)
b9+ [9Me](1007.683)
y9+ [6Me](1020.552)
b11+ [0Me](1055.619)
b10+ [10Me](1108.730)
y10+ [7Me](1121.591)
y11+ [8Me](1249.650)
y12+ [9Me](1376.748)
y13+ [10Me](1489.833)
y14+ [11Me](1616.934)
y15+ [12Me](1701.981)

I

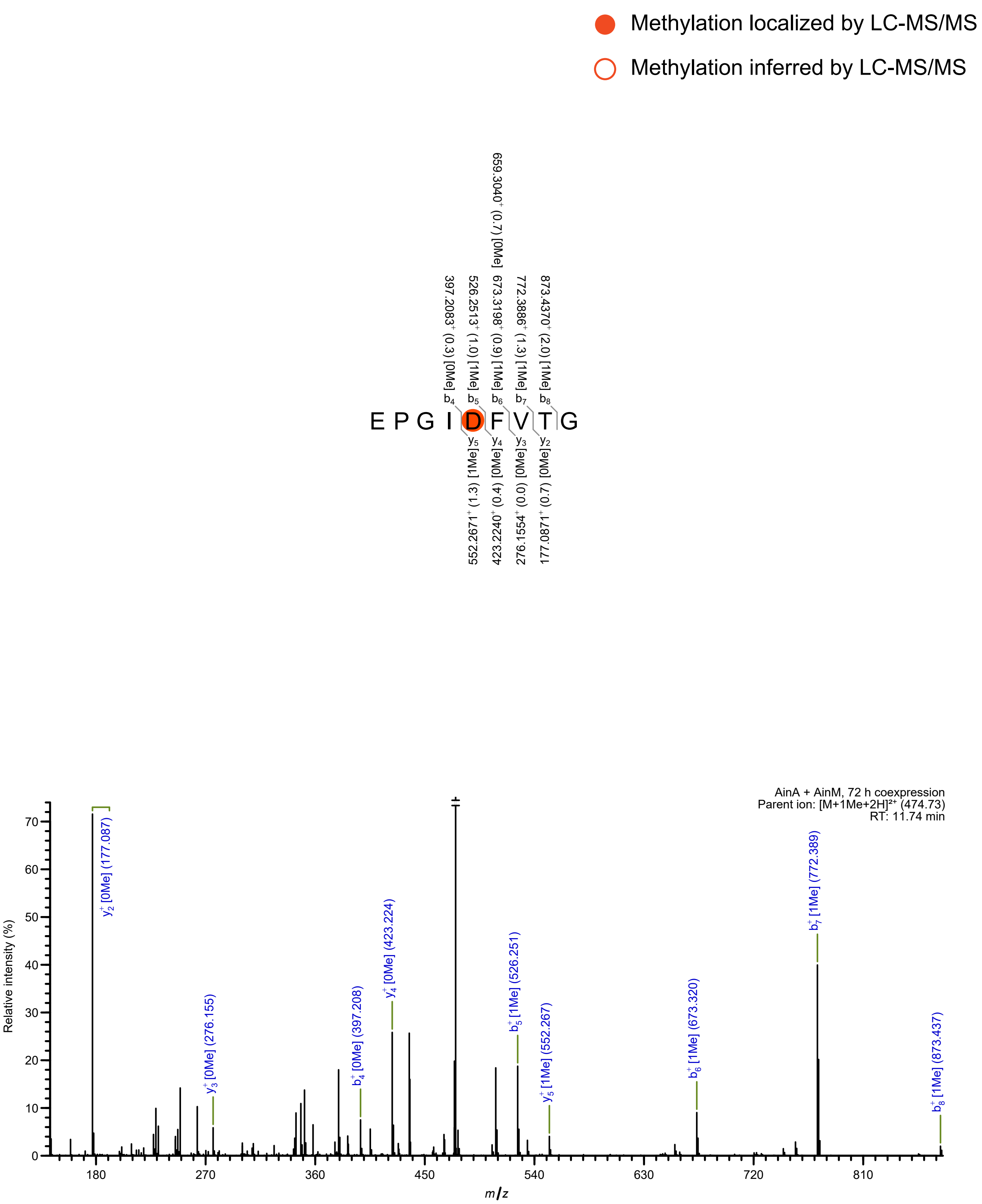

Methylation localized by LC-MS/MS
Methylation inferred by LC-MS/MS
E P G I D F V T G
b4 397.2083+ (0.3) [0Me]
b5 526.2513+ (1.0) [1Me]
b6 673.3198+ (0.9) [1Me] 659.3040+ (0.7) [0Me]
b7 772.3886+ (1.3) [1Me]
b8 873.4370+ (2.0) [1Me]
y5 [1Me] (1.3) 552.2671+
y4 [0Me] (0.4) 423.2240+
y3 [0Me] (0.0) 276.1554+
y2 [0Me] (0.7) 177.0871+
AinA + AinM, 72 h coexpression
Parent ion: [M+1Me+2H]2+ (474.73)
RT: 11.74 min
Relative intensity (%)
m/z
y2+ [0Me] (177.087)
y3+ [0Me] (276.155)
b4+ [0Me] (397.208)
y4+ [0Me] (423.224)
b5+ [1Me] (526.251)
y5+ [1Me] (552.267)
b6+ [1Me] (673.320)
b7+ [1Me] (772.389)
b8+ [1Me] (873.437)

m

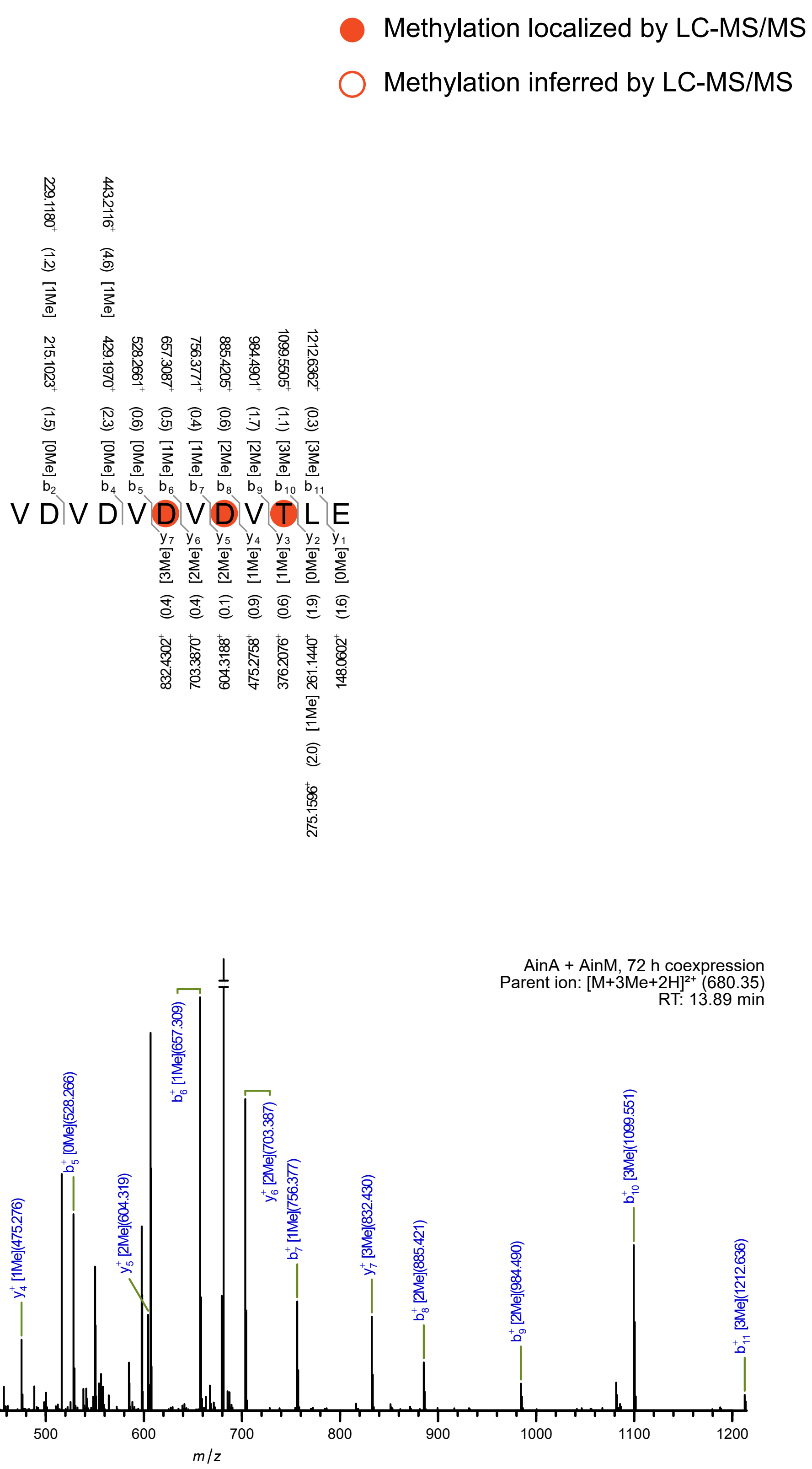
Methylation localized by LC-MS/MS
Methylation inferred by LC-MS/MS
V D V D V D V D V T L E
AinA + AinM, 72 h coexpression
Parent ion: [M+3Me+2H]²⁺ (680.35)
RT: 13.89 min
Relative intensity (%)
m/z

n

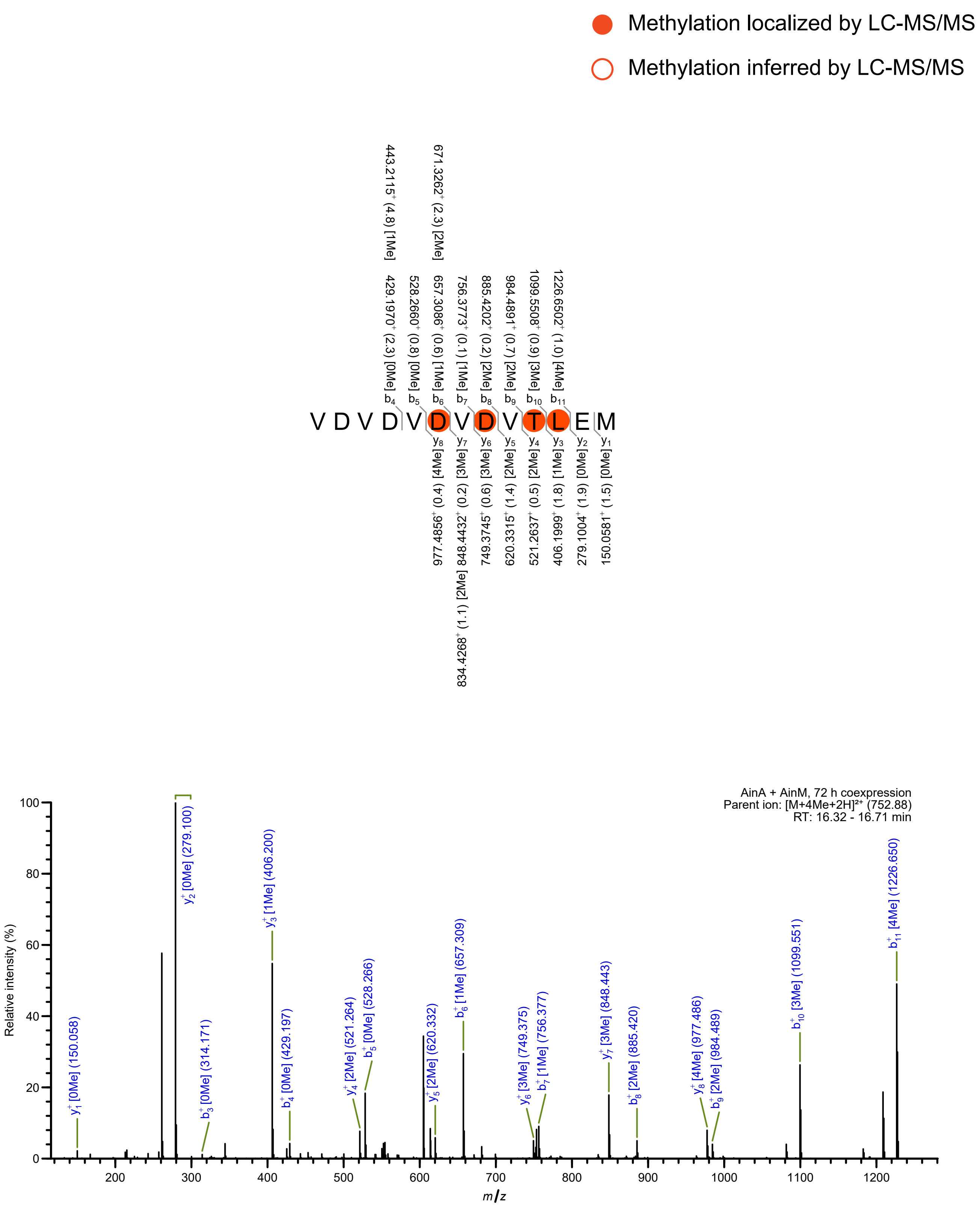

Methylation localized by LC-MS/MS
Methylation inferred by LC-MS/MS
V D V D V D V D V T L E M
AinA + AinM, 72 h coexpression
Parent ion: [M+4Me+2H]²⁺ (752.88)
RT: 16.32 - 16.71 min
Relative intensity (%)
m/z

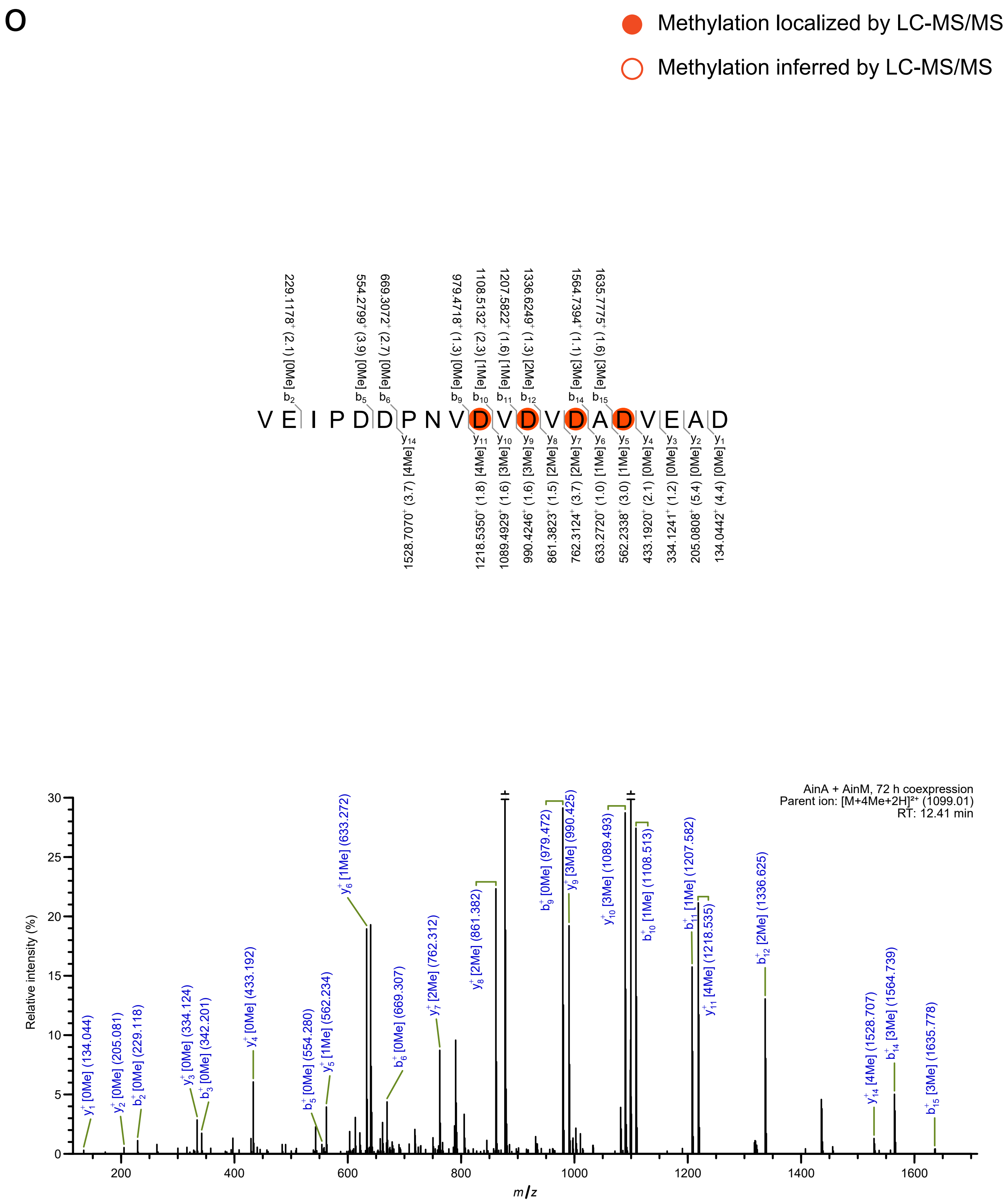

O
Methylation localized by LC-MS/MS
Methylation inferred by LC-MS/MS
V E I P D D P N V D V D V D A D V E A D
AinA + AinM, 72 h coexpression
Parent ion: [M+4Me+2H]²⁺ (1099.01)
RT: 12.41 min
Relative intensity (%)
m/z

p

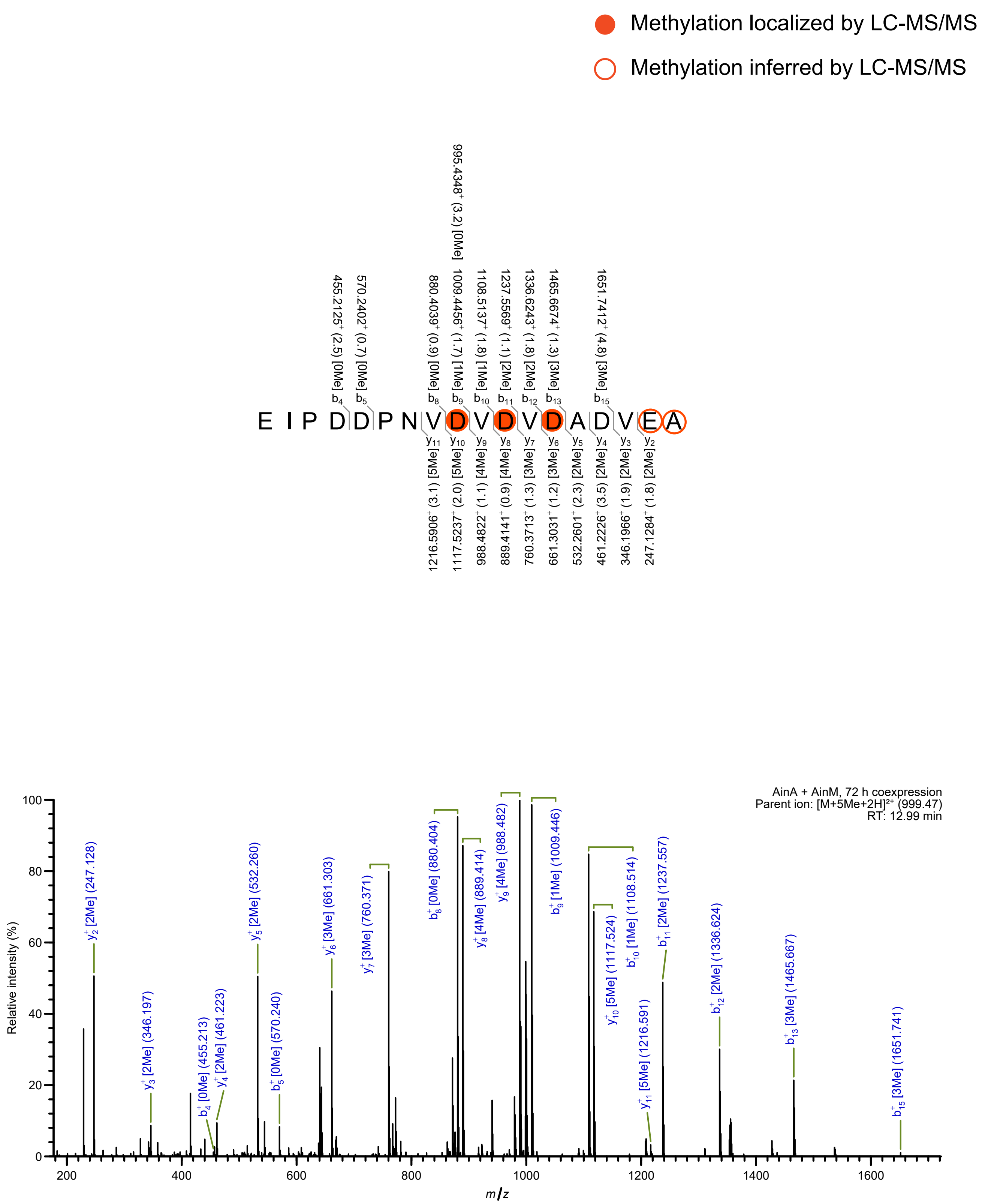

Methylation localized by LC-MS/MS
Methylation inferred by LC-MS/MS
E I P D D P N V D V D V D A D V E A
AinA + AinM, 72 h coexpression
Parent ion: [M+5Me+2H]2+ (999.47)
RT: 12.99 min
Relative intensity (%)
m/z

q

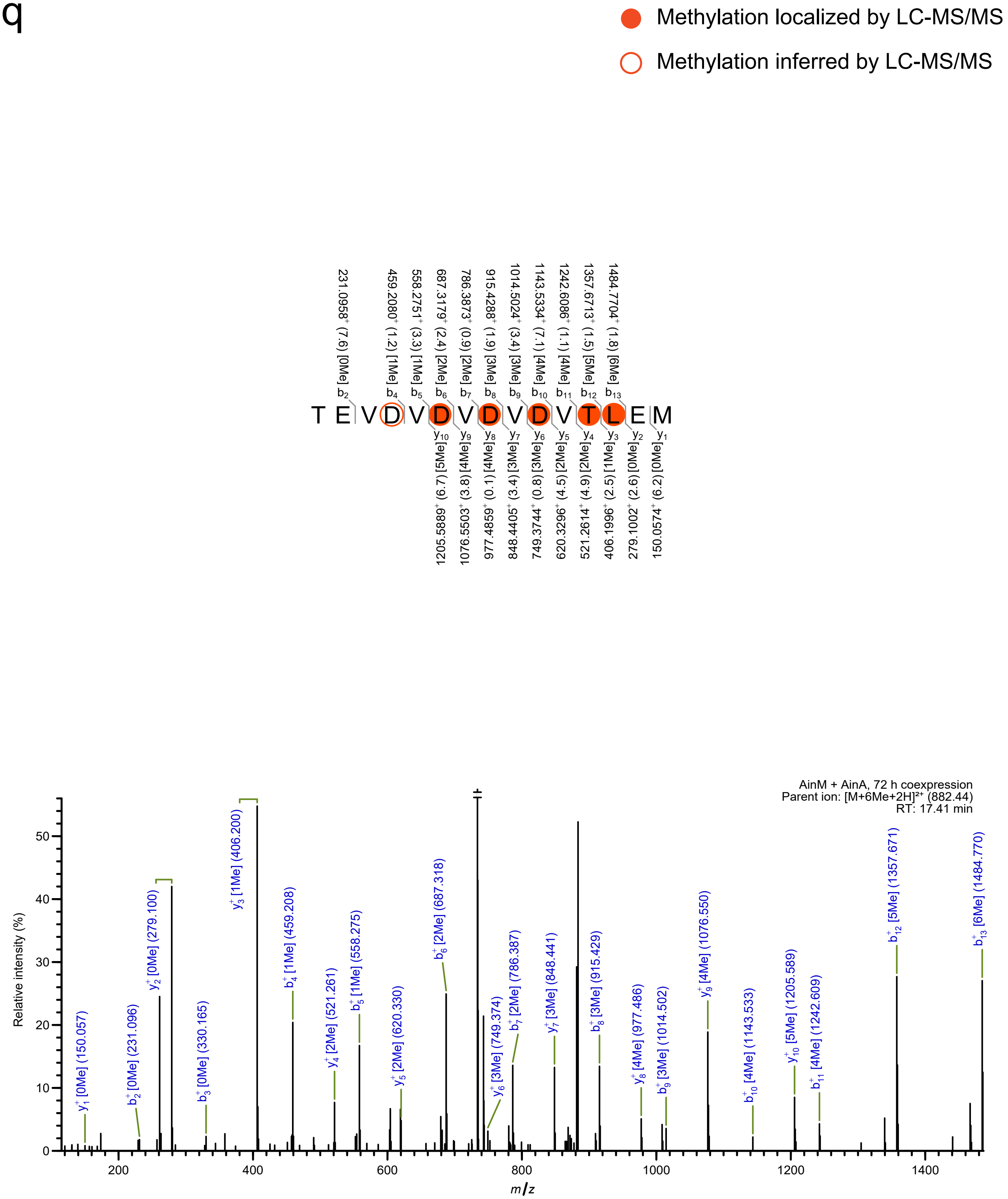
Methylation localized by LC-MS/MS
Methylation inferred by LC-MS/MS
T E V D V D V D V D V T L E M
231.0958⁺ (7.6) [0Me] b2
459.2080⁺ (1.2) [1Me] b4
558.2751⁺ (3.3) [1Me] b5
687.3179⁺ (2.4) [2Me] b6
786.3873⁺ (0.9) [2Me] b7
915.4288⁺ (1.9) [3Me] b8
1014.5024⁺ (3.4) [3Me] b9
1143.5334⁺ (7.1) [4Me] b10
1242.6086⁺ (1.1) [4Me] b11
1357.6713⁺ (1.5) [5Me] b12
1484.7704⁺ (1.8) [6Me] b13
y10 1205.5889⁺ (6.7) [5Me]
y9 1076.5503⁺ (3.8) [4Me]
y8 977.4859⁺ (0.1) [4Me]
y7 848.4405⁺ (3.4) [3Me]
y6 749.3744⁺ (0.8) [3Me]
y5 620.3296⁺ (4.5) [2Me]
y4 521.2614⁺ (4.9) [2Me]
y3 406.1996⁺ (2.5) [1Me]
y2 279.1002⁺ (2.6) [0Me]
y1 150.0574⁺ (6.2) [0Me]
AinM + AinA, 72 h coexpression
Parent ion: [M+6Me+2H]²⁺ (882.44)
RT: 17.41 min
Relative intensity (%)
m/z
y1⁺ [0Me] (150.057)
b2⁺ [0Me] (231.096)
y2⁺ [0Me] (279.100)
b3⁺ [0Me] (330.165)
y3⁺ [1Me] (406.200)
b4⁺ [1Me] (459.208)
y4⁺ [2Me] (521.261)
b5⁺ [1Me] (558.275)
y5⁺ [2Me] (620.330)
b6⁺ [2Me] (687.318)
y6⁺ [3Me] (749.374)
b7⁺ [2Me] (786.387)
y7⁺ [3Me] (848.441)
b8⁺ [3Me] (915.429)
y8⁺ [4Me] (977.486)
b9⁺ [3Me] (1014.502)
y9⁺ [4Me] (1076.550)
b10⁺ [4Me] (1143.533)
y10⁺ [5Me] (1205.589)
b11⁺ [4Me] (1242.609)
b12⁺ [5Me] (1357.671)
b13⁺ [6Me] (1484.770)

r

Methylation localized by LC-MS/MS

Methylation inferred by LC-MS/MS

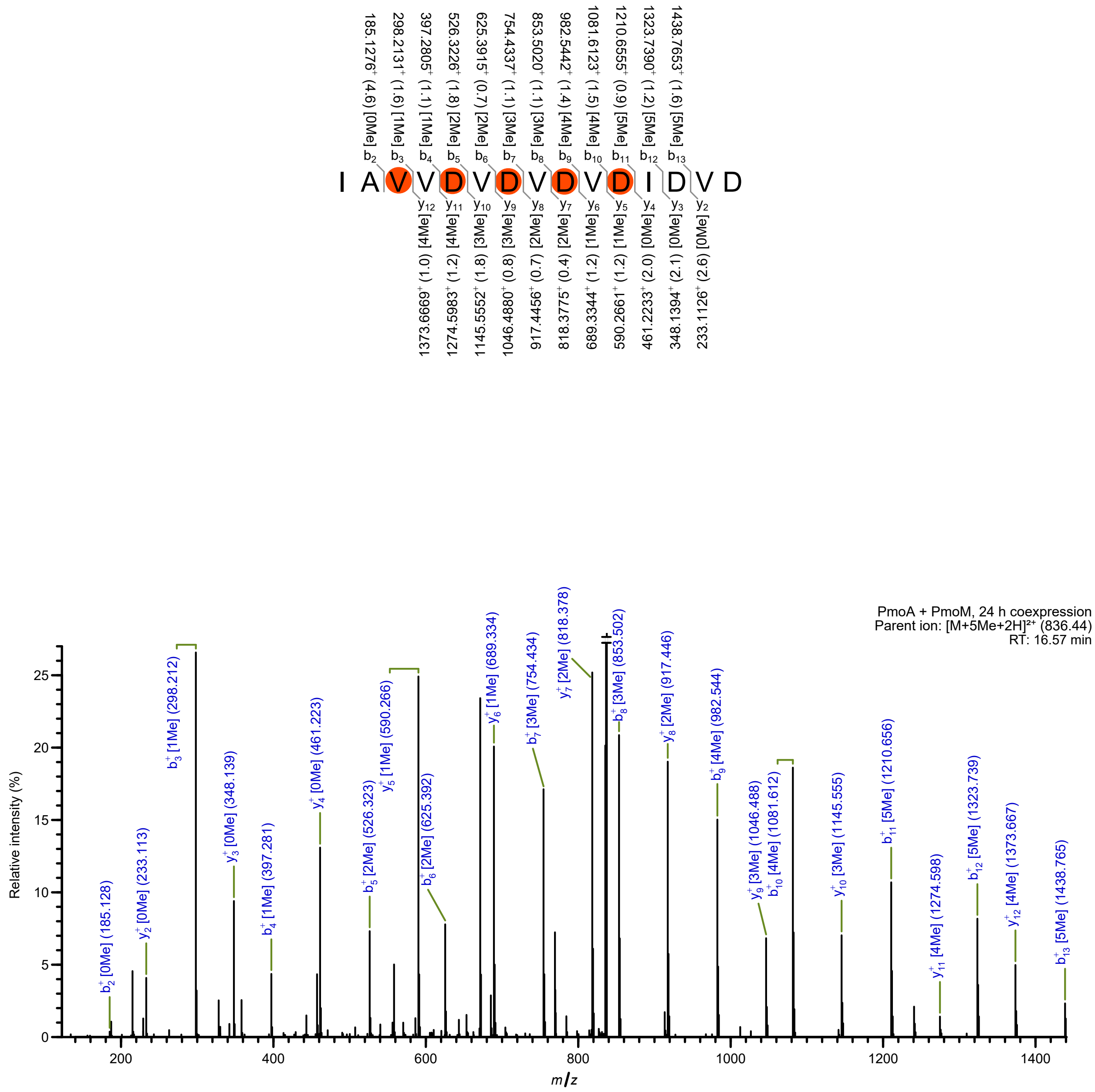

S

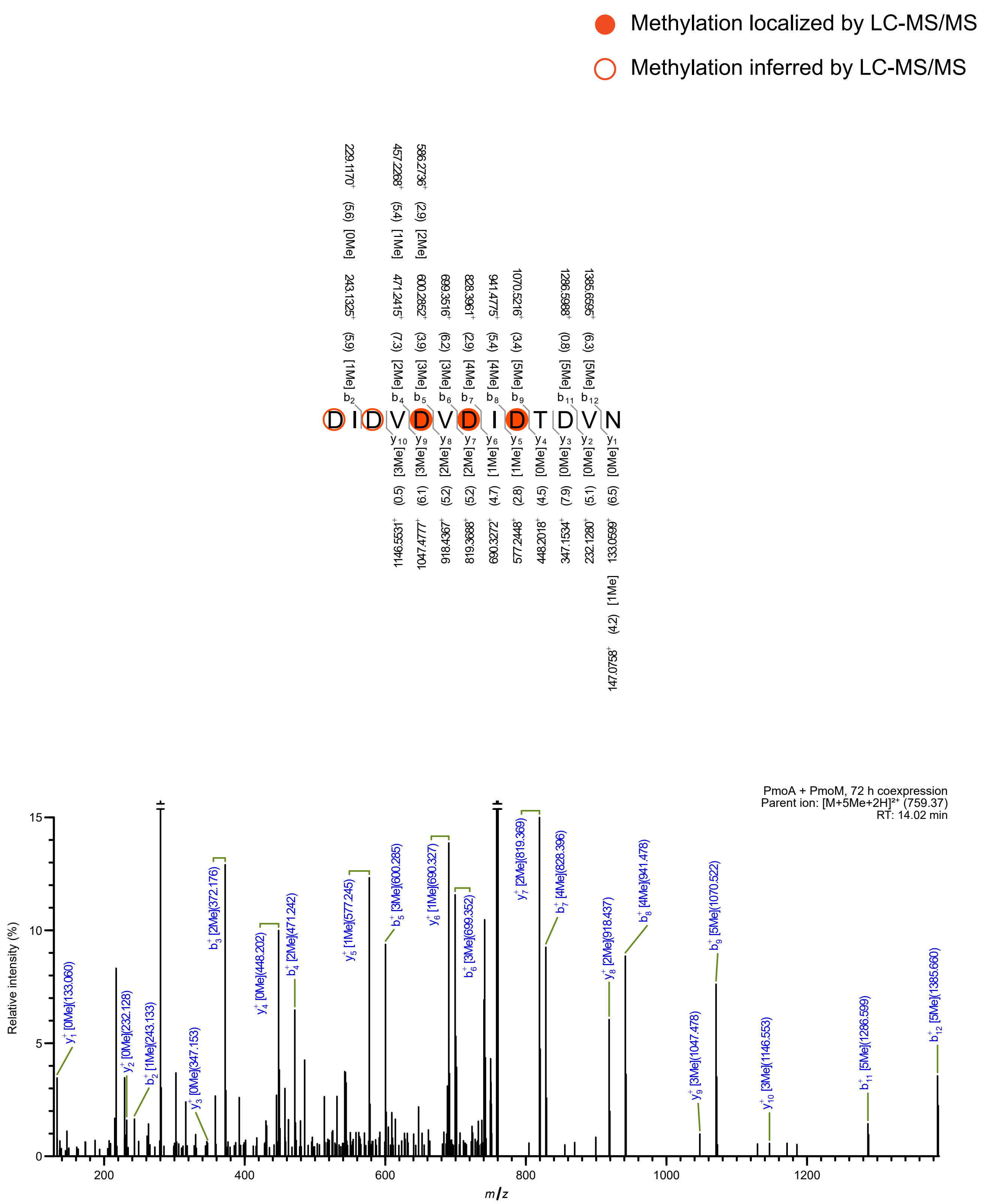

Methylation localized by LC-MS/MS
Methylation inferred by LC-MS/MS
D I D V D V D I D T D V N
PmoA + PmoM, 72 h coexpression
Parent ion: [M+5Me+2H]²⁺ (759.37)
RT: 14.02 min
Relative intensity (%)
m/z

t

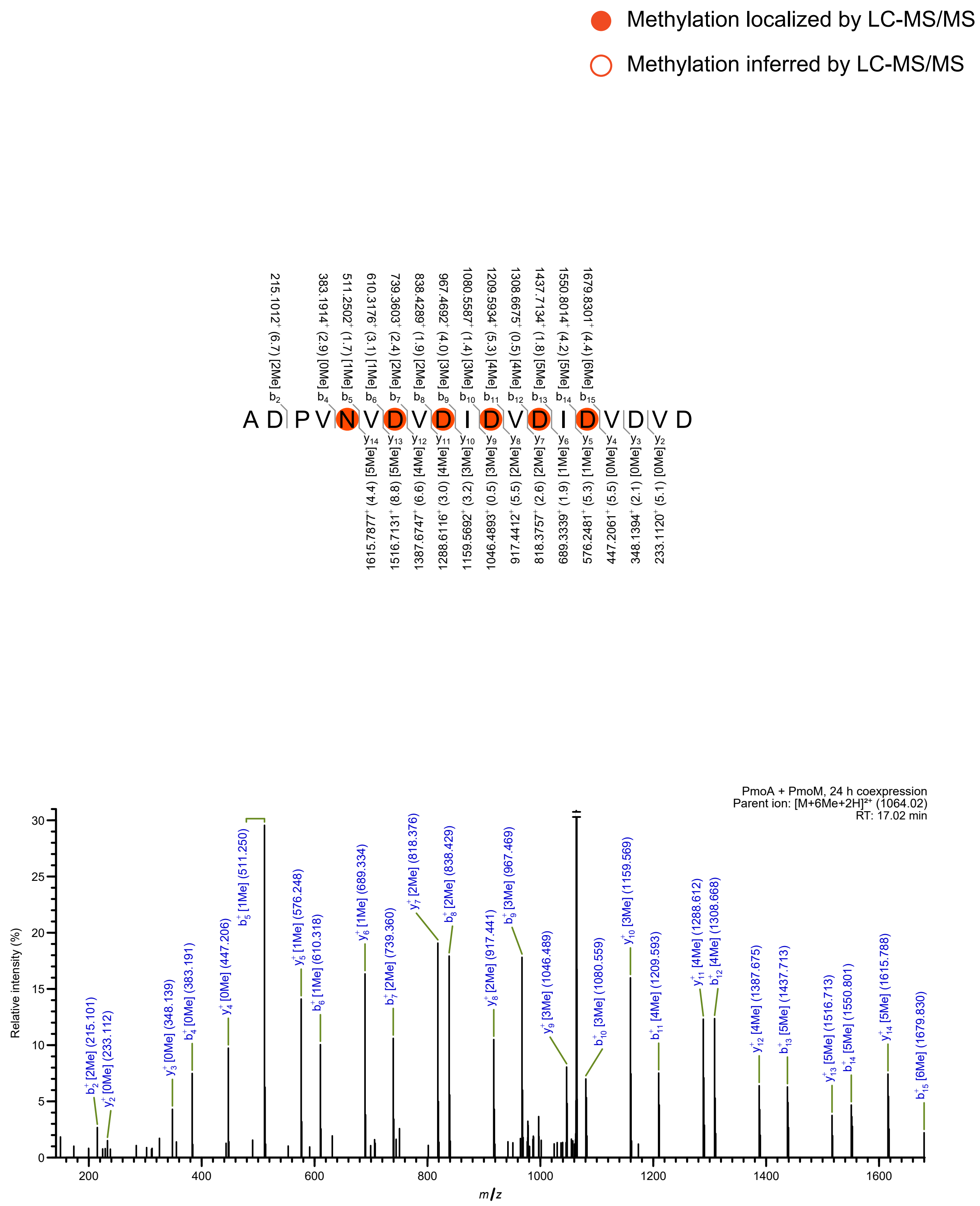

Methylation localized by LC-MS/MS
Methylation inferred by LC-MS/MS
A D P V N V D V D I D V D I D V D V D
215.1012⁺ (6.7) [2Me] b2
383.1914⁺ (2.9) [0Me] b4
511.2502⁺ (1.7) [1Me] b5
610.3176⁺ (3.1) [1Me] b6
739.3603⁺ (2.4) [2Me] b7
838.4289⁺ (1.9) [2Me] b8
967.4692⁺ (4.0) [3Me] b9
1080.5587⁺ (1.4) [3Me] b10
1209.5934⁺ (5.3) [4Me] b11
1308.6675⁺ (0.5) [4Me] b12
1437.7134⁺ (1.8) [5Me] b13
1550.8014⁺ (4.2) [5Me] b14
1679.8301⁺ (4.4) [6Me] b15
y14 [5Me] (4.4) 1615.7877⁺
y13 [5Me] (8.8) 1516.7131⁺
y12 [4Me] (6.6) 1387.6747⁺
y11 [4Me] (3.0) 1288.6116⁺
y10 [3Me] (3.2) 1159.5692⁺
y9 [3Me] (0.5) 1046.4893⁺
y8 [2Me] (5.5) 917.4412⁺
y7 [2Me] (2.6) 818.3757⁺
y6 [1Me] (1.9) 689.3339⁺
y5 [1Me] (5.3) 576.2481⁺
y4 [0Me] (5.5) 447.2061⁺
y3 [0Me] (2.1) 348.1394⁺
y2 [0Me] (5.1) 233.1120⁺
PmoA + PmoM, 24 h coexpression
Parent ion: [M+6Me+2H]²⁺ (1064.02)
RT: 17.02 min
Relative intensity (%)
m/z
b2⁺ [2Me] (215.101)
y2⁺ [0Me] (233.112)
y3⁺ [0Me] (348.139)
b4⁺ [0Me] (383.191)
y4⁺ [0Me] (447.206)
b5⁺ [1Me] (511.250)
y5⁺ [1Me] (576.248)
b6⁺ [1Me] (610.318)
y6⁺ [1Me] (689.334)
b7⁺ [2Me] (739.360)
y7⁺ [2Me] (818.376)
b8⁺ [2Me] (838.429)
y8⁺ [2Me] (917.441)
b9⁺ [3Me] (967.469)
y9⁺ [3Me] (1046.489)
b10⁺ [3Me] (1080.559)
y10⁺ [3Me] (1159.569)
b11⁺ [4Me] (1209.593)
y11⁺ [4Me] (1288.612)
b12⁺ [4Me] (1308.668)
y12⁺ [4Me] (1387.675)
b13⁺ [5Me] (1437.713)
y13⁺ [5Me] (1516.713)
b14⁺ [5Me] (1550.801)
y14⁺ [5Me] (1615.788)
b15⁺ [6Me] (1679.830)

u

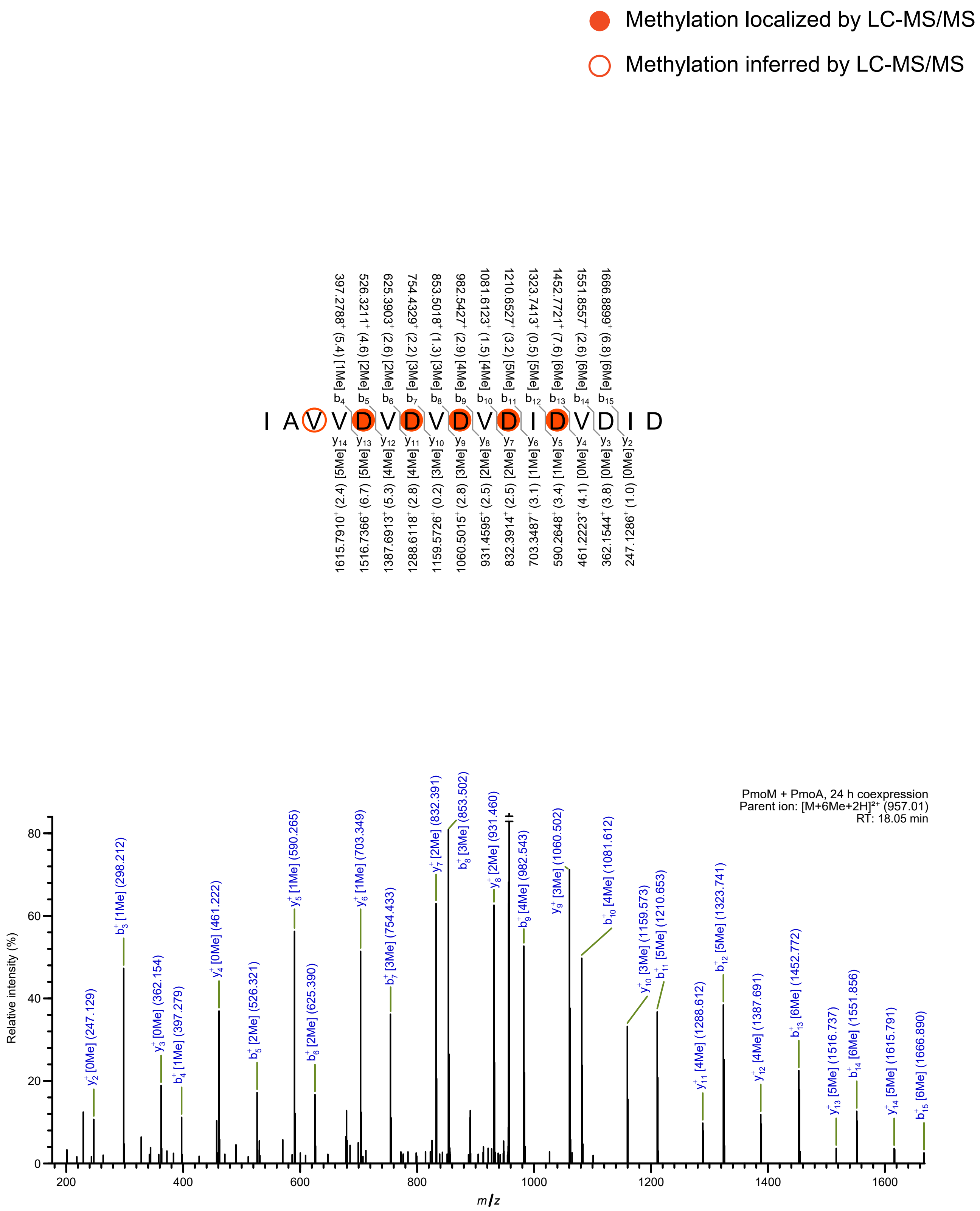
Methylation localized by LC-MS/MS
Methylation inferred by LC-MS/MS
I A V V D V D V D V D I D V D I D
PmoM + PmoA, 24 h coexpression
Parent ion: [M+6Me+2H]²⁺ (957.01)
RT: 18.05 min
Relative intensity (%)
m/z

**V**

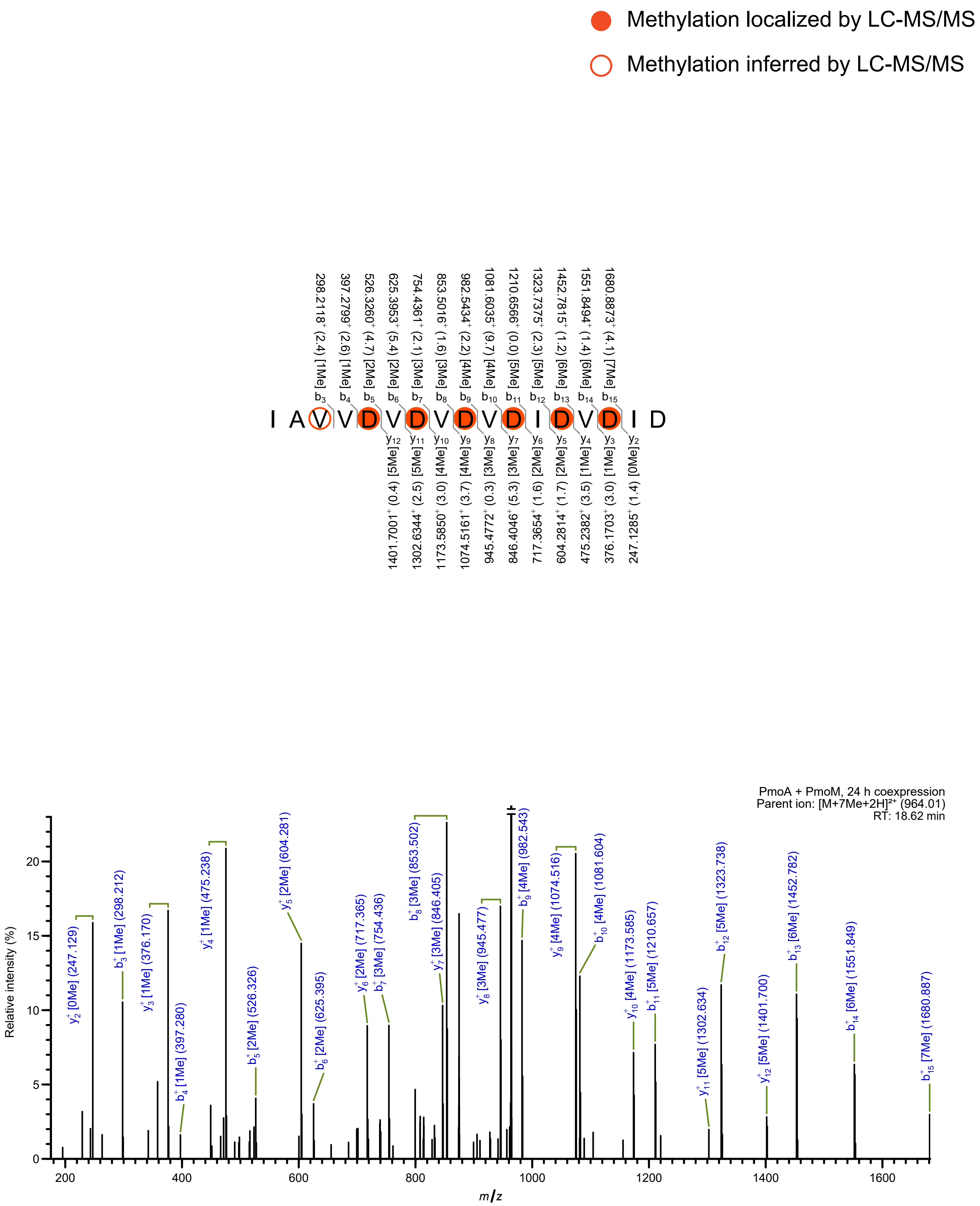
Methylation localized by LC-MS/MS
Methylation inferred by LC-MS/MS
I A V V D V D V D V D I D V D I D
b3 298.2118+ (2.4) [1Me]
b4 397.2799+ (2.6) [1Me]
b5 526.3260+ (4.7) [2Me]
b6 625.3953+ (5.4) [2Me]
b7 754.4361+ (2.1) [3Me]
b8 853.5016+ (1.6) [3Me]
b9 982.5434+ (2.2) [4Me]
b10 1081.6035+ (9.7) [4Me]
b11 1210.6566+ (0.0) [5Me]
b12 1323.7375+ (2.3) [5Me]
b13 1452.7815+ (1.2) [6Me]
b14 1551.8494+ (1.4) [6Me]
b15 1680.8873+ (4.1) [7Me]
y12 1401.7001+ (0.4) [5Me]
y11 1302.6344+ (2.5) [5Me]
y10 1173.5850+ (3.0) [4Me]
y9 1074.5161+ (3.7) [4Me]
y8 945.4772+ (0.3) [3Me]
y7 846.4046+ (5.3) [3Me]
y6 717.3654+ (1.6) [2Me]
y5 604.2814+ (1.7) [2Me]
y4 475.2382+ (3.5) [1Me]
y3 376.1703+ (3.0) [1Me]
y2 247.1285+ (1.4) [0Me]
PmoA + PmoM, 24 h coexpression
Parent ion: [M+7Me+2H]2+ (964.01)
RT: 18.62 min
Relative intensity (%)
m/z
y2+ [0Me] (247.129)
b3+ [1Me] (298.212)
y3+ [1Me] (376.170)
b4+ [1Me] (397.280)
y4+ [1Me] (475.238)
b5+ [2Me] (526.326)
y5+ [2Me] (604.281)
b6+ [2Me] (625.395)
y6+ [2Me] (717.365)
b7+ [3Me] (754.436)
b8+ [3Me] (853.502)
y7+ [3Me] (846.405)
y8+ [3Me] (945.477)
b9+ [4Me] (982.543)
y9+ [4Me] (1074.516)
b10+ [4Me] (1081.604)
y10+ [4Me] (1173.585)
b11+ [5Me] (1210.657)
y11+ [5Me] (1302.634)
b12+ [5Me] (1323.738)
y12+ [5Me] (1401.700)
b13+ [6Me] (1452.782)
b14+ [6Me] (1551.849)
b15+ [7Me] (1680.887)

W

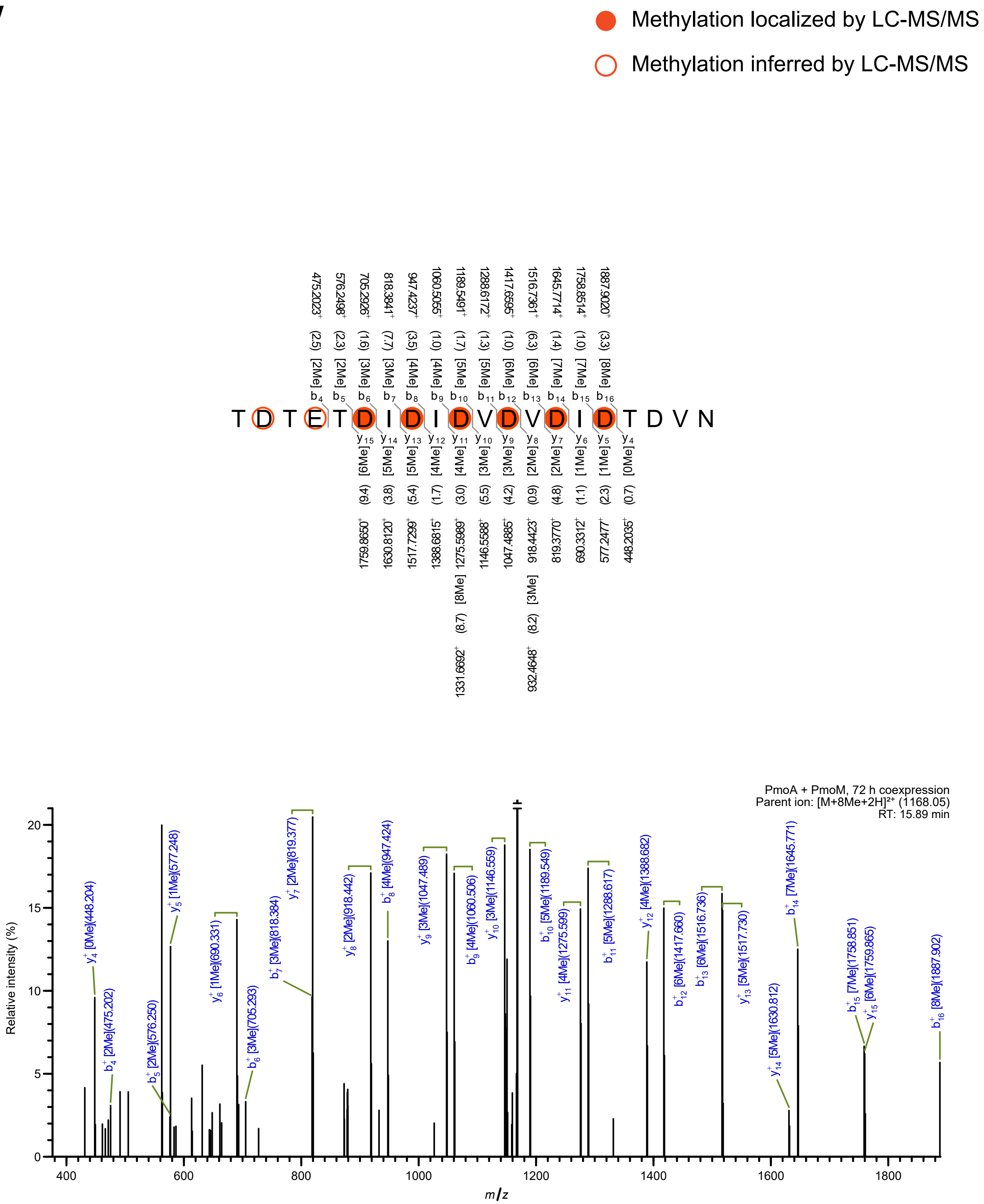

Methylation localized by LC-MS/MS
Methylation inferred by LC-MS/MS
T D T E T D I D I D V D V D I D T D V N
PmoA + PmoM, 72 h coexpression
Parent ion: [M+8Me+2H]²⁺ (1168.05)
RT: 15.89 min
Relative intensity (%)
m/z

X

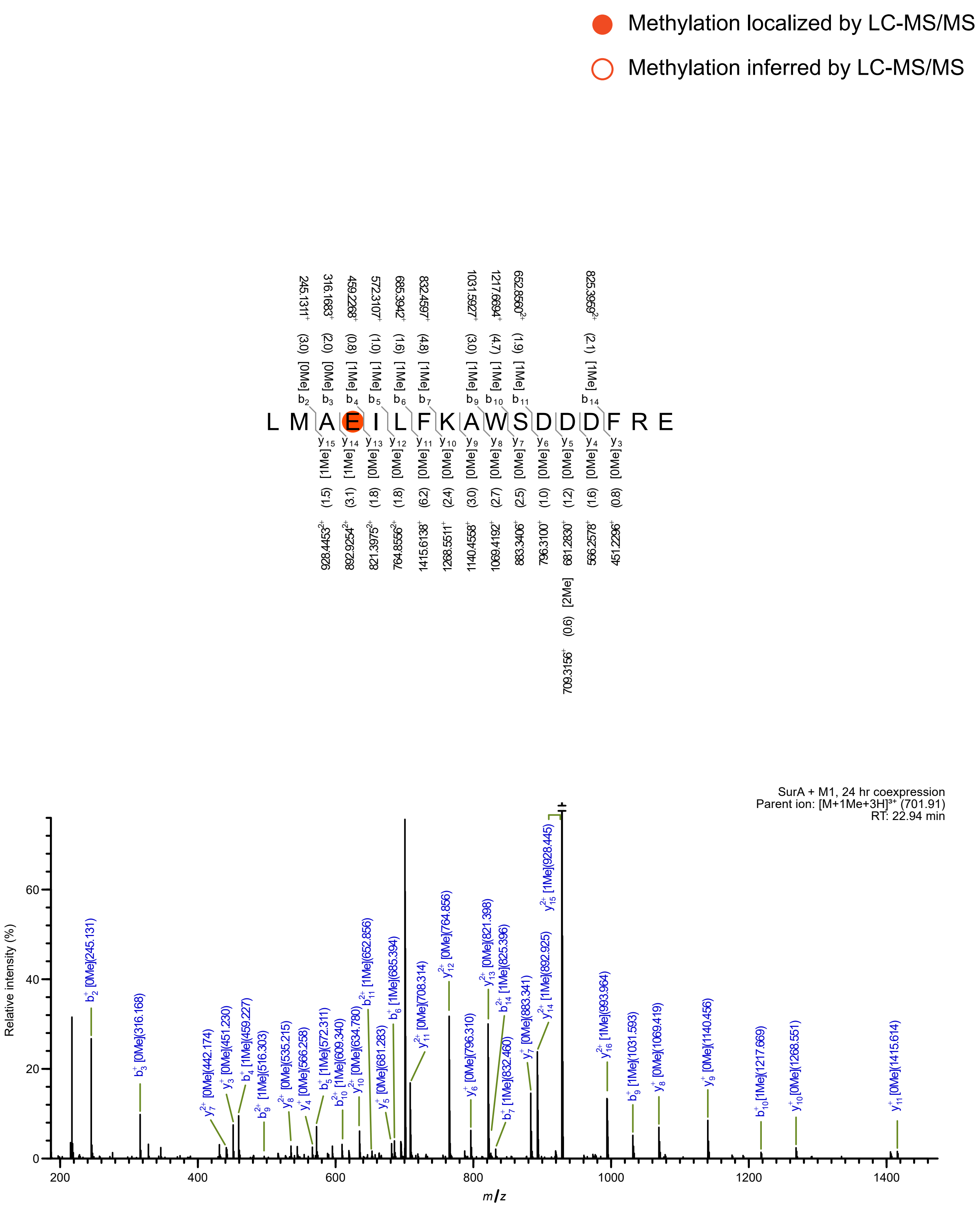

Methylation localized by LC-MS/MS
Methylation inferred by LC-MS/MS
L M A E I L F K A W S D D D F R E
SurA + M1, 24 hr coexpression
Parent ion: [M+1Me+3H]³⁺ (701.91)
RT: 22.94 min
Relative intensity (%)
m/z

y

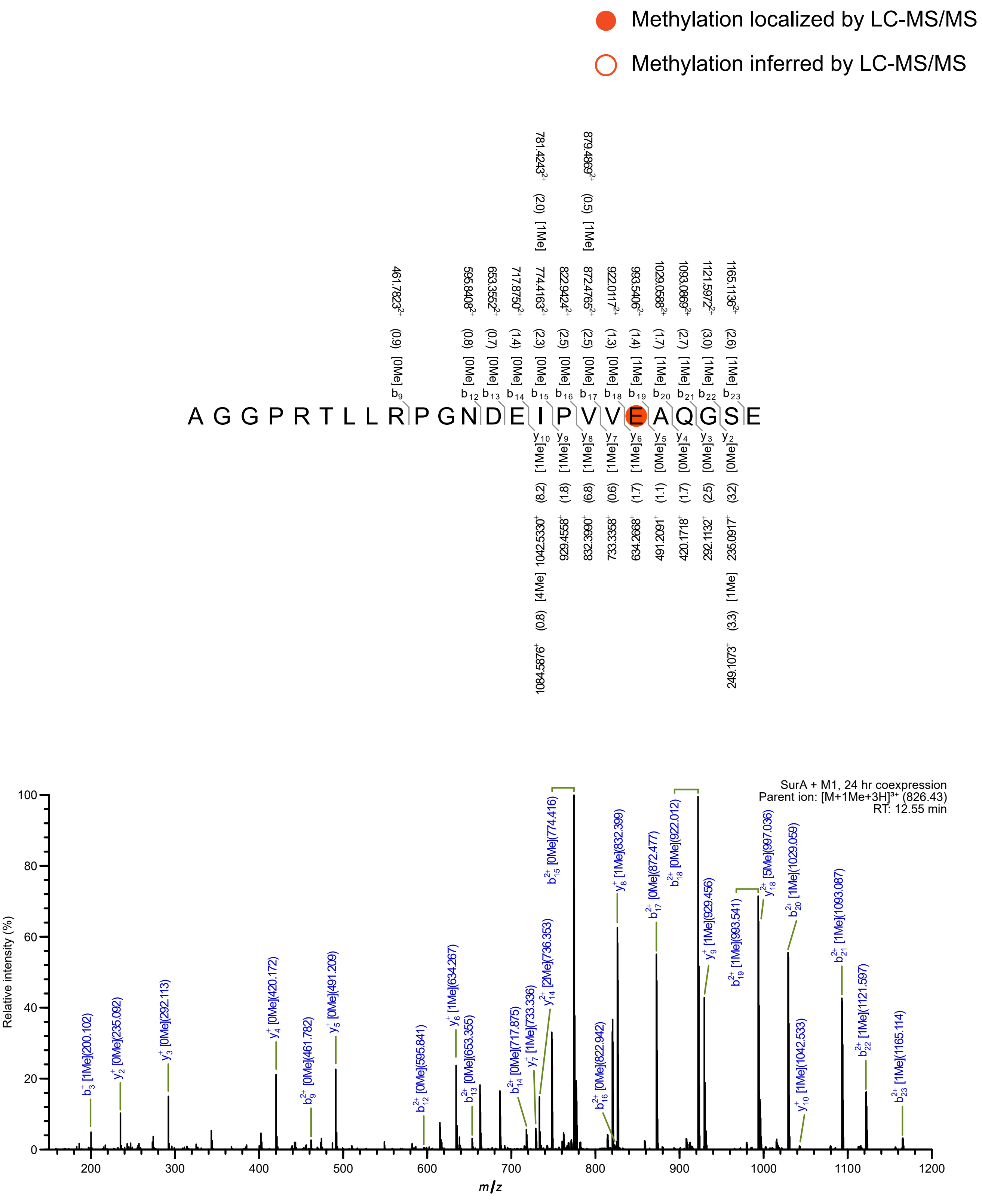

Methylation localized by LC-MS/MS
Methylation inferred by LC-MS/MS
A G G P R T L L R P G N D E I P V V E A Q G S E
SurA + M1, 24 hr coexpression
Parent ion: [M+1Me+3H]3+ (826.43)
RT: 12.55 min
Relative intensity (%)
m/z

Z

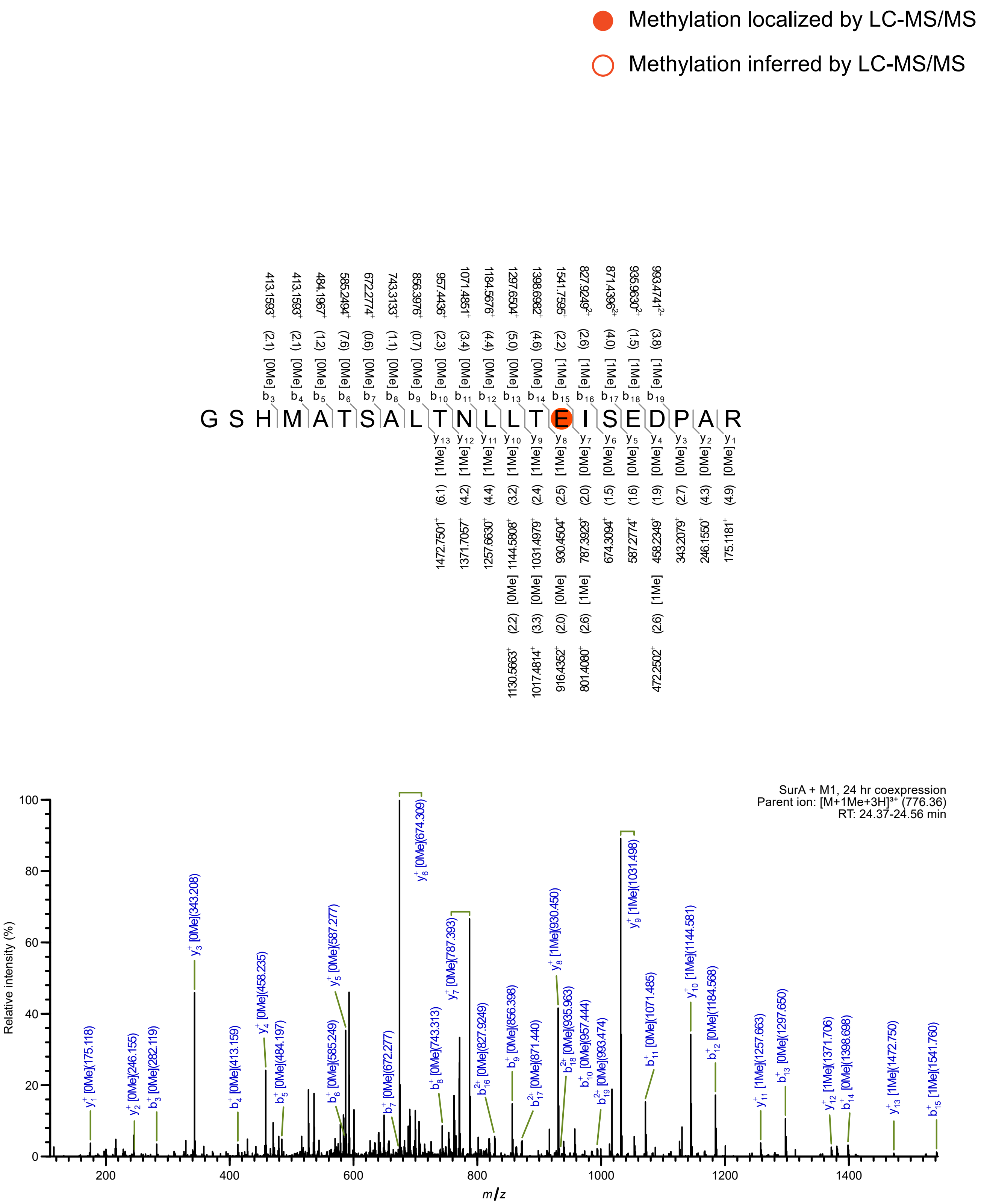

Methylation localized by LC-MS/MS
Methylation inferred by LC-MS/MS
G S H M A T S A L T N L L T E I S E D P A R
SurA + M1, 24 hr coexpression
Parent ion: [M+1Me+3H]³⁺ (776.36)
RT: 24.37-24.56 min
Relative intensity (%)
m/z

aa

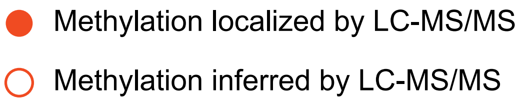
Methylation localized by LC-MS/MS
Methylation inferred by LC-MS/MS

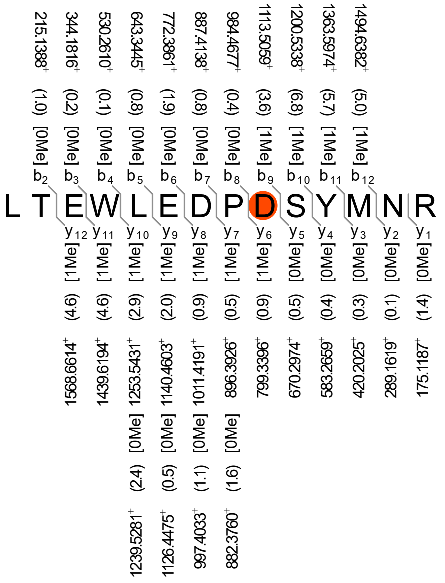
L T E W L E D P D S Y M N R

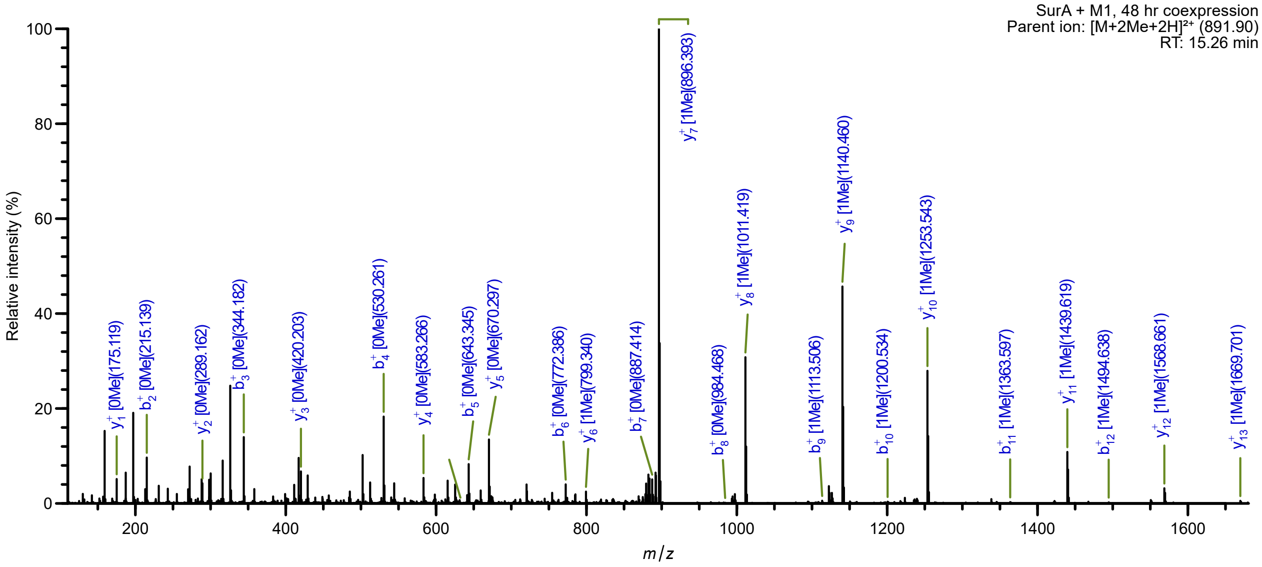
SurA + M1, 48 hr coexpression
Parent ion: [M+2Me+2H]²⁺ (891.90)
RT: 15.26 min
Relative intensity (%)
m/z

ab

Methylation localized by LC-MS/MS

Methylation inferred by LC-MS/MS

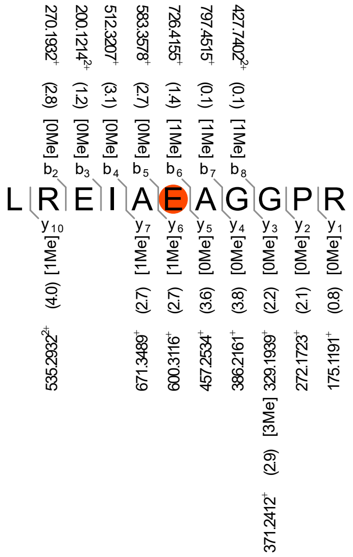

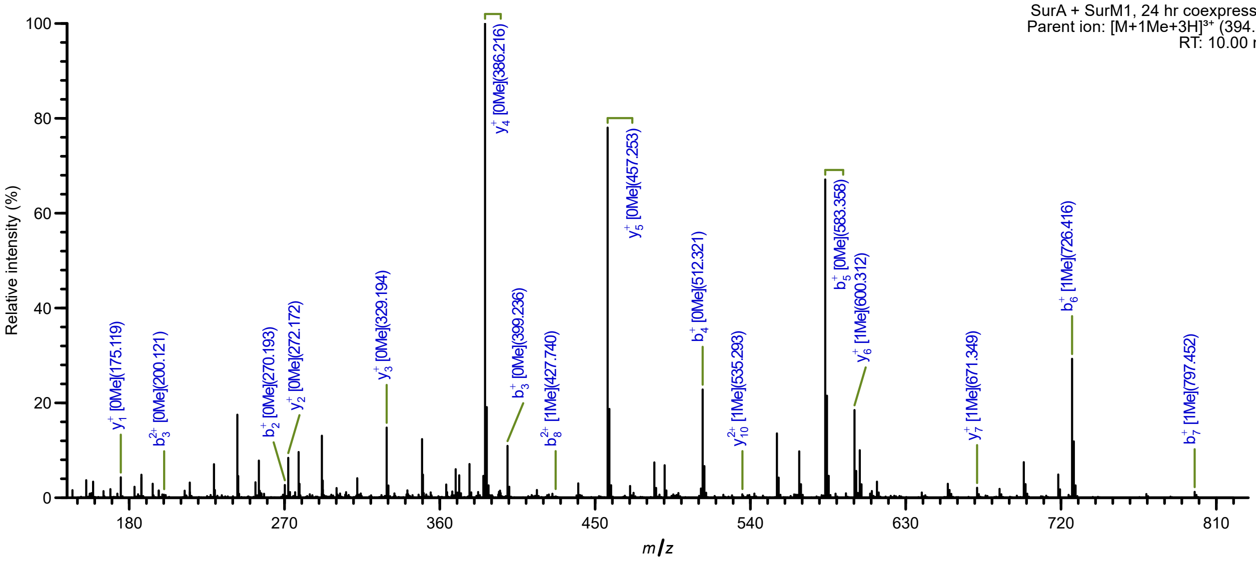

ac

Methylation localized by LC-MS/MS

Methylation inferred by LC-MS/MS

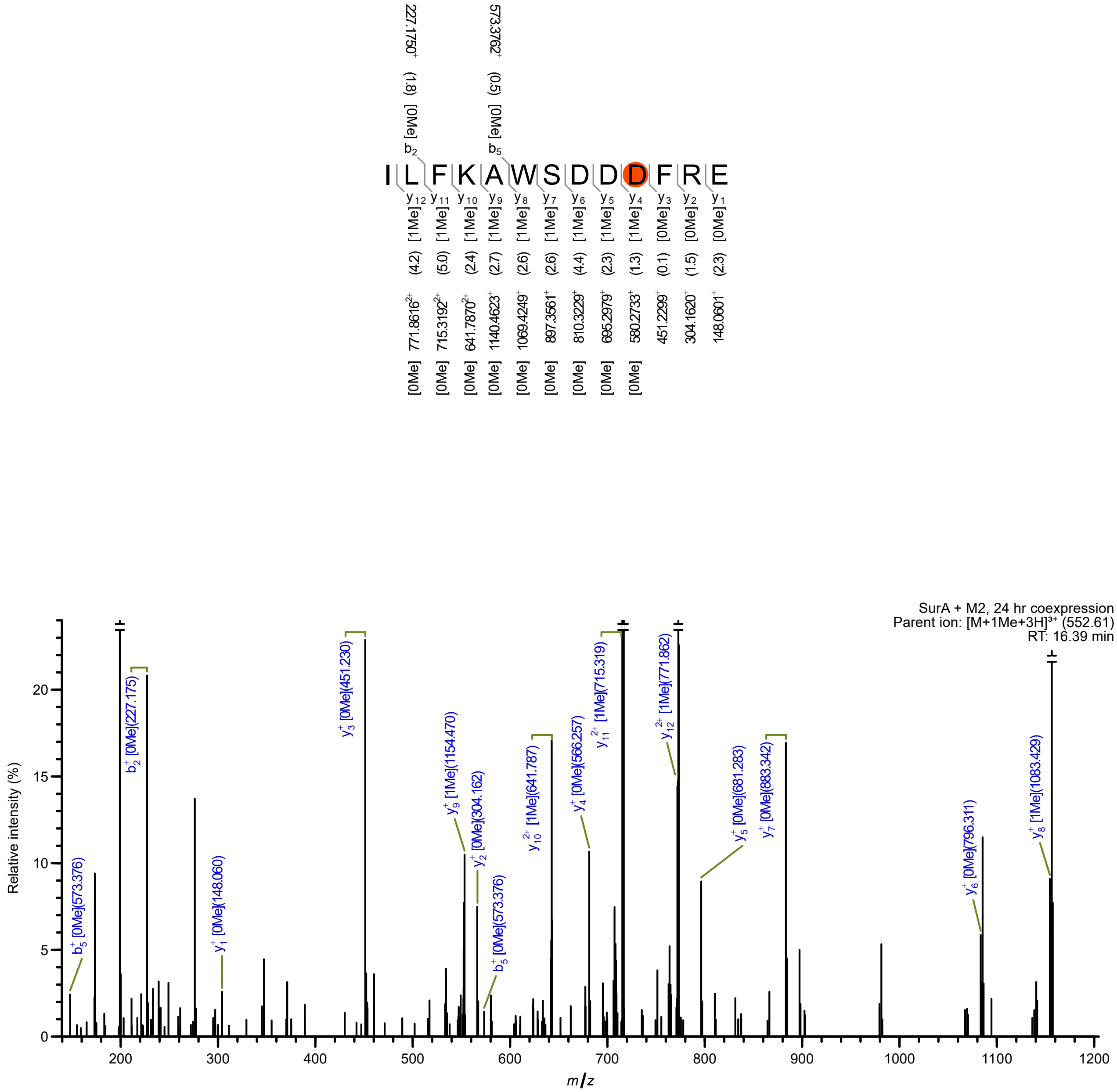

ad

Methylation localized by LC-MS/MS

Methylation inferred by LC-MS/MS

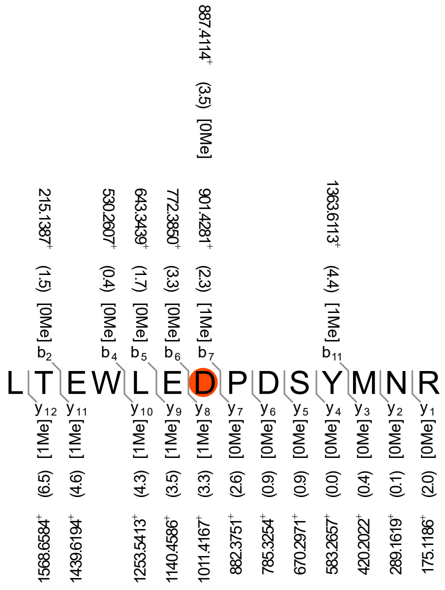

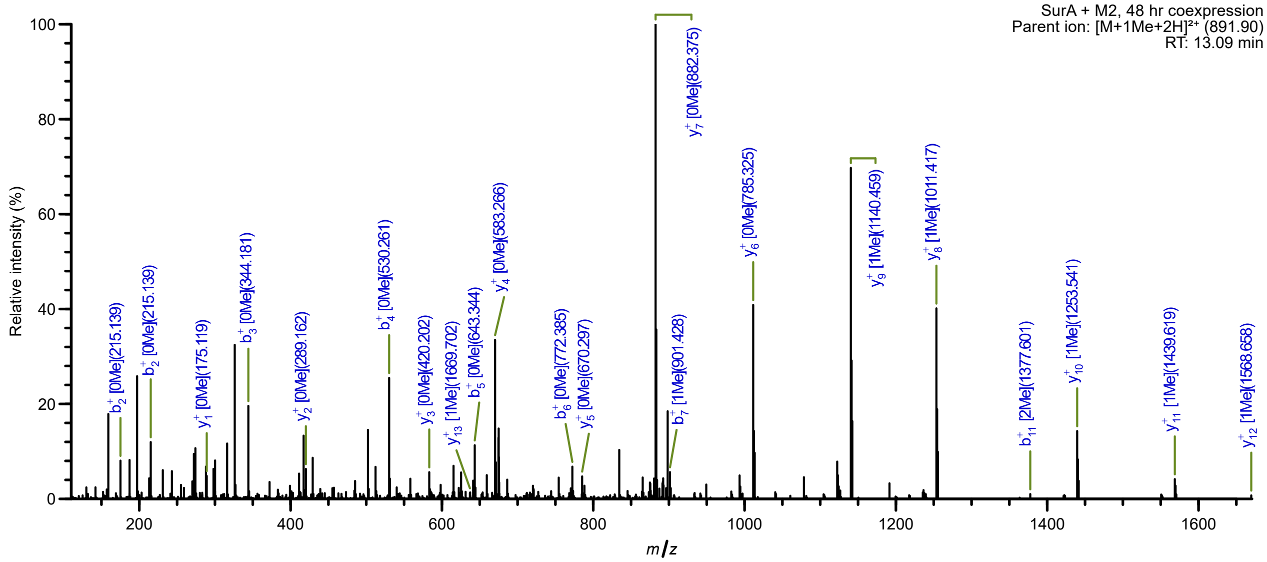

ae

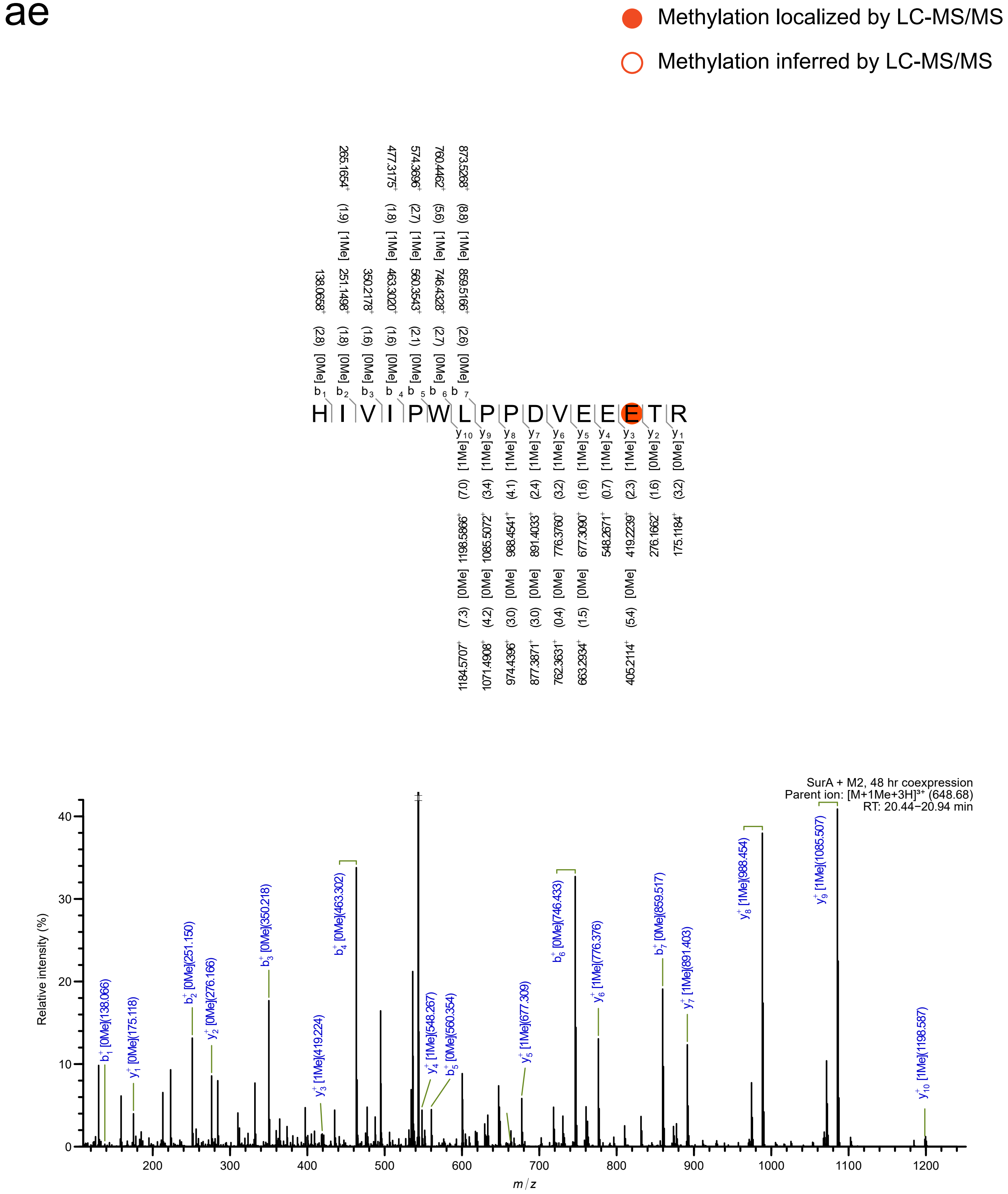
Methylation localized by LC-MS/MS
Methylation inferred by LC-MS/MS
HIVIPWLPPDVEEETR
SurA + M2, 48 hr coexpression
Parent ion: [M+1Me+3H]³⁺ (648.68)
RT: 20.44–20.94 min
Relative intensity (%)
m/z

af

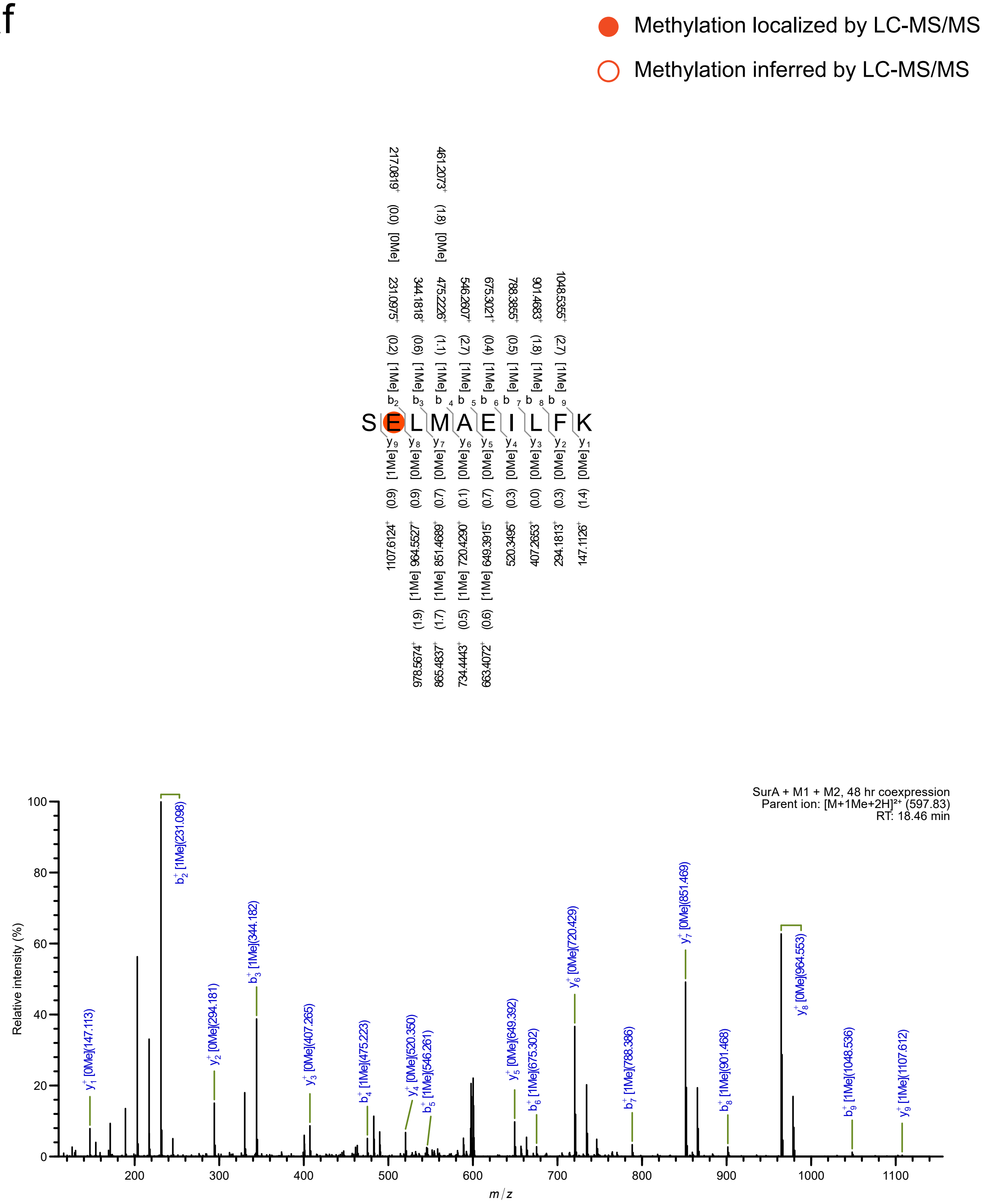

Methylation localized by LC-MS/MS
Methylation inferred by LC-MS/MS
SELMAEILFK
SurA + M1 + M2, 48 hr coexpression
Parent ion: [M+1Me+2H]²⁺ (597.83)
RT: 18.46 min
Relative intensity (%)
m/z

ag

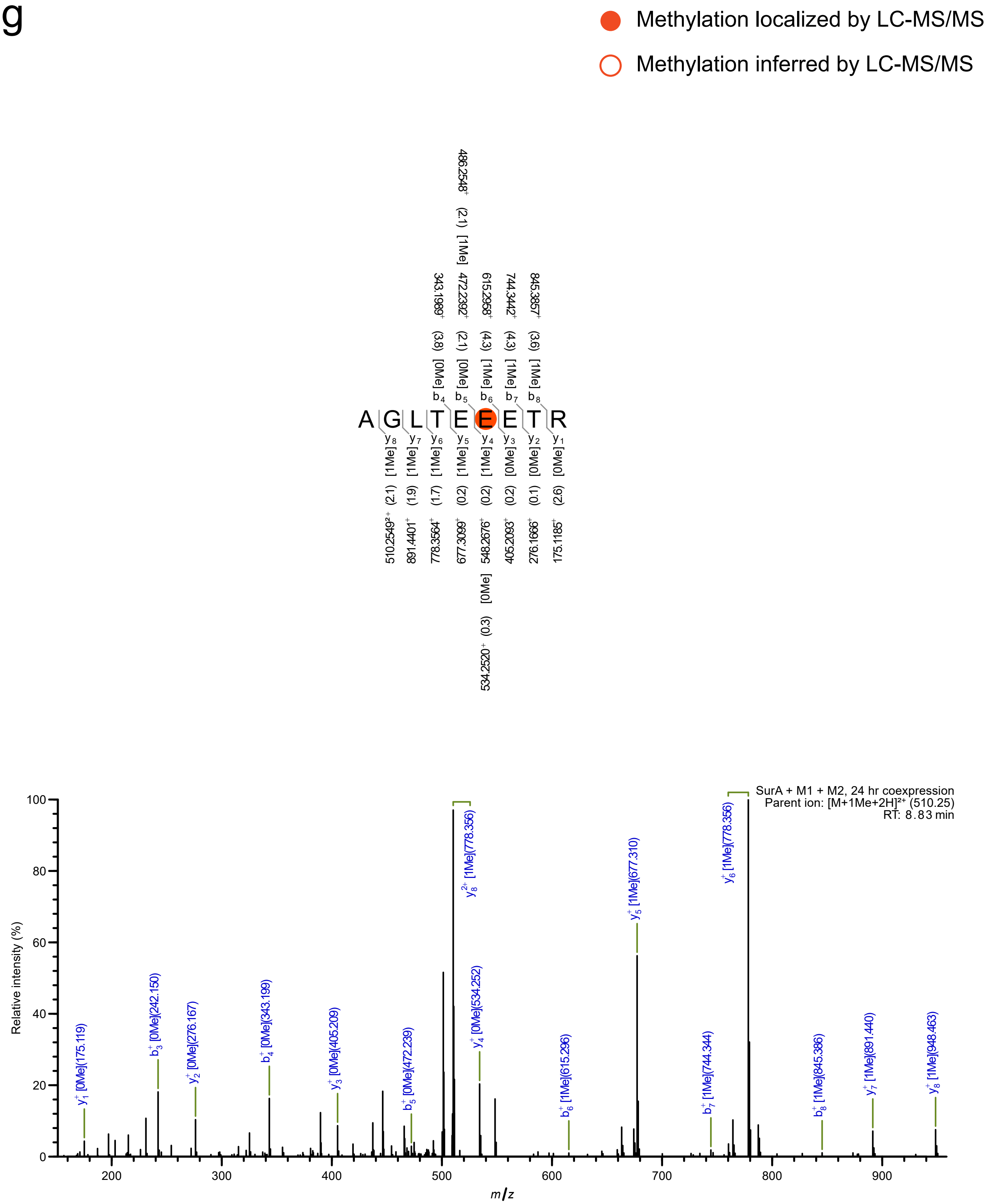

Methylation localized by LC-MS/MS
Methylation inferred by LC-MS/MS
A G L T E E E T R
486.2548+ (2.1) [1Me]
343.1989+ (3.8) [0Me] b4
472.2392+ (2.1) [0Me] b5
615.2958+ (4.3) [1Me] b6
744.3442+ (4.3) [1Me] b7
845.3857+ (3.6) [1Me] b8
y8 [1Me] (2.1) 510.2549²+
y7 [1Me] (1.9) 891.4401+
y6 [1Me] (1.7) 778.3564+
y5 [1Me] (0.2) 677.3099+
y4 [1Me] (0.2) 548.2676+
[0Me] (0.3) 534.2520+
y3 [0Me] (0.2) 405.2093+
y2 [0Me] (0.1) 276.1666+
y1 [0Me] (2.6) 175.1185+
SurA + M1 + M2, 24 hr coexpression
Parent ion: [M+1Me+2H]²⁺ (510.25)
RT: 8.83 min
Relative intensity (%)
m/z
y1+ [0Me](175.119)
b3+ [0Me](242.150)
y2+ [0Me](276.167)
b4+ [0Me](343.199)
y3+ [0Me](405.209)
b5+ [0Me](472.239)
y8 2+ [1Me](778.356)
y4+ [0Me](534.252)
b6+ [1Me](615.296)
y5+ [1Me](677.310)
b7+ [1Me](744.344)
y6+ [1Me](778.356)
b8+ [1Me](845.386)
y7+ [1Me](891.440)
y8+ [1Me](948.463)

ah

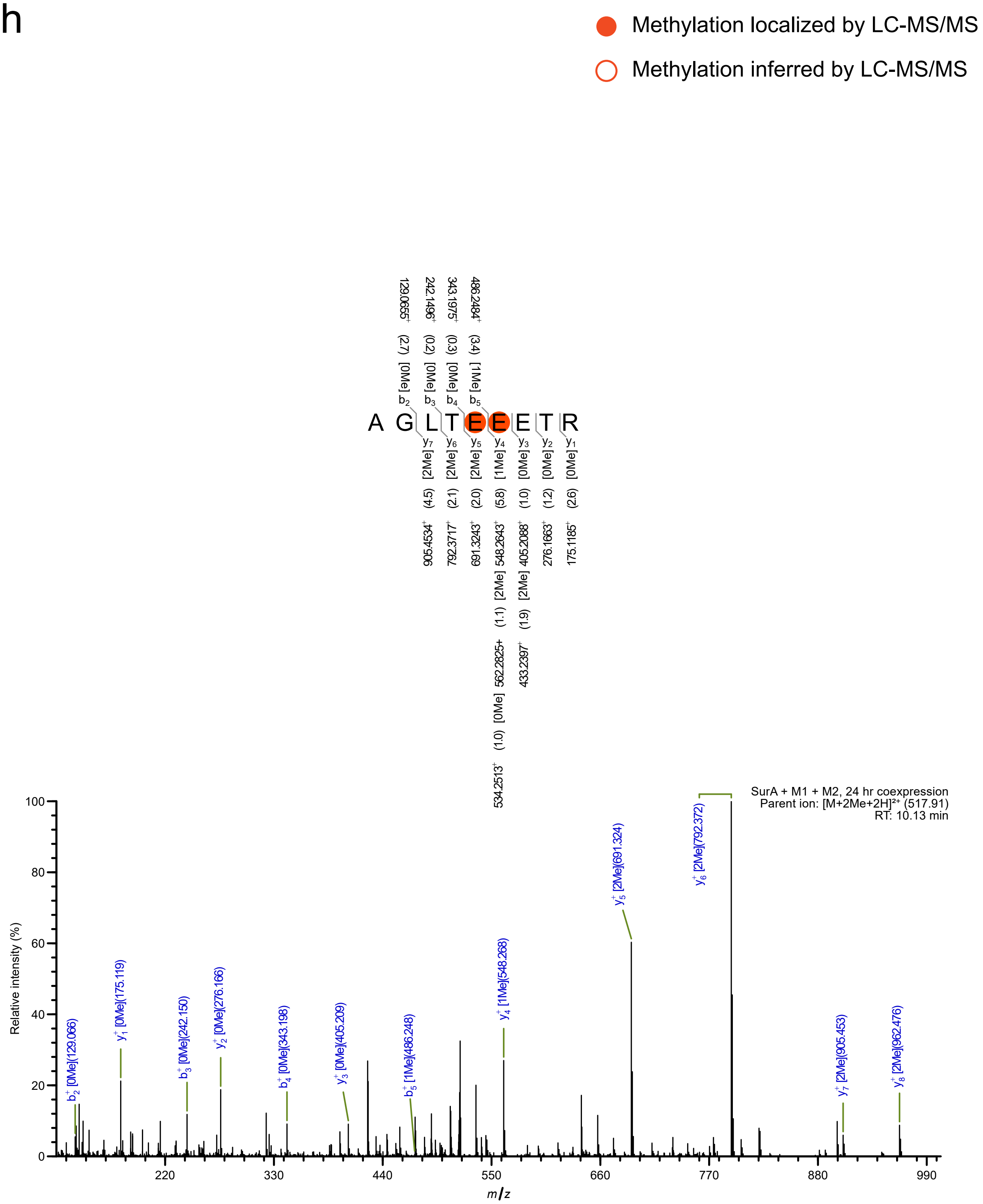

Methylation localized by LC-MS/MS
Methylation inferred by LC-MS/MS
A G L T E E E T R
SurA + M1 + M2, 24 hr coexpression
Parent ion: [M+2Me+2H]²⁺ (517.91)
RT: 10.13 min
Relative intensity (%)
m/z

ai

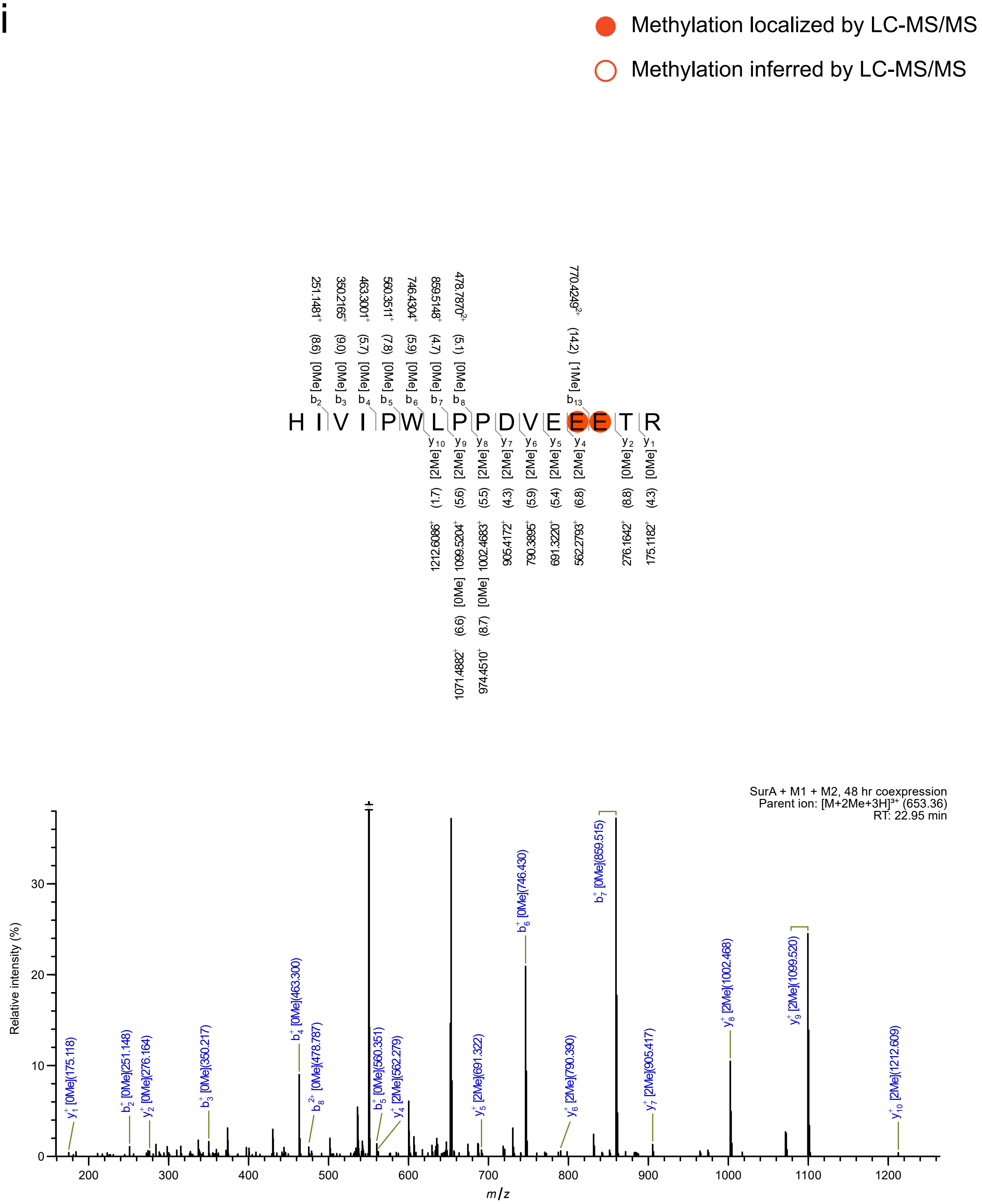
Methylation localized by LC-MS/MS
Methylation inferred by LC-MS/MS
H I V I P W L P P D V E E E T R
SurA + M1 + M2, 48 hr coexpression
Parent ion: [M+2Me+3H]³⁺ (653.36)
RT: 22.95 min
Relative intensity (%)
m/z

**Fig. S2: Location of RceA S78F mutation for LC-MS/MS analysis.** Alternate decapeptide repeats are shown in grey italics for better visibility.

>RceA

MTTIVPTELDQPDVIELSGGEL*DVAELSGGEL*DVAELFGGEL*DVAELSGGEL*DVAE
LSGGEL*DVAELSGGEL*DVAELSGGEL*DVAELSGGEL*DVAELSGGEL*DVAELSGGE*
*L*DVAEIGIINTFDL

Ser to Phe substitution

**Fig S3. Size exclusion chromatography (SEC) of characterized NMTs**. α-*N*-Methyltransferase enzymes purified from *E. coli* BL21(DE3) or LOBSTR(DE3) elute as high molecular weight aggregates (1st peak) followed by active fractions (typically dimers). Inset: SDS-PAGE profile of SEC peak. Protein of interest is indicated with a "◀". **(a)** His-RceM (theoretical [dimer]: 89 kD; observed [soluble aggregate or tetramer]: 220 kD); **(b)** His-SliM (theoretical [dimer]: 122 kD; observed [dimer]: 151 kD); **(c)** His-AinM (theoretical [dimer]: 226 kD); observed [dimer]: 270 kD); **(d)** His-SUMO PmoM (theoretical [dimer]: 179 kD; observed [dimer]: 153 kD); **(e)** His-SurM1 (theoretical [monomer]: 61 kD; observed [monomer]: 64 kD); **(f)** His-SurM2 (theoretical [dimer]: 62 kD; observed [dimer]: 69 kD); **(g)** The calibration curve used to estimate the observed molecular weight. For calibration of the column the following size markers were used: vitamin B12 (1.35 kD), myoglobin (17 kD), ovalbumin (44 kD), γ-globulin (158 kD), and thyroglobulin (670 kD). The Y-axis represents the partition coefficient (Kav) and X-axis represents the log of the molecular weight (MW).

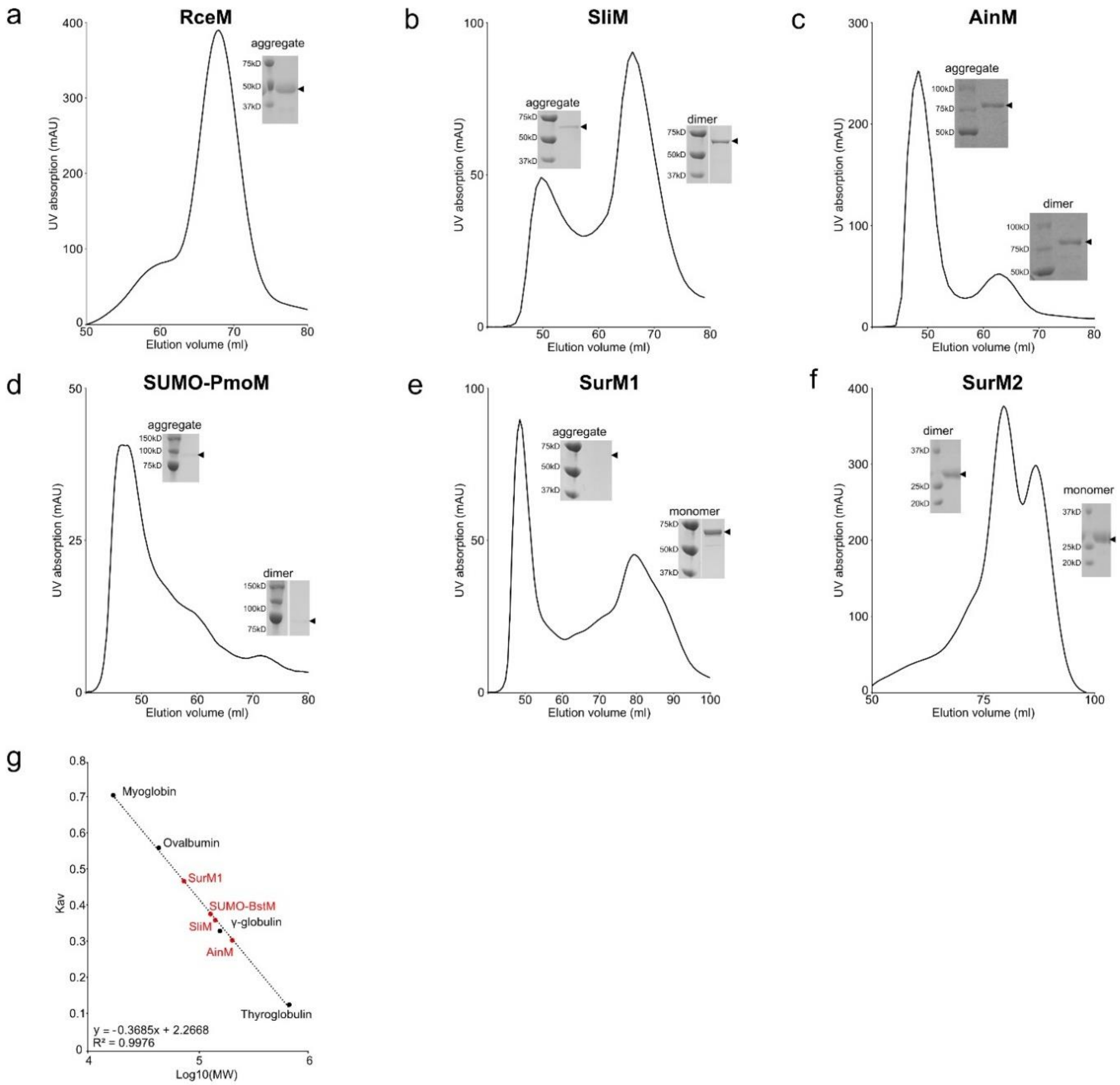

**Fig. S4. AlphaFold2 protein models of NMTs and precursors**. Structures are colored according to their per-residue confidence score (pLDDT). Sequence alignment quality graphs for each protein are displayed alongside their predicted structures. Despite high confidence predictions (red) in parts of the precursor proteins, the sequence alignment quality graph shows that much fewer sequences were used in model construction as compared to NMTs.

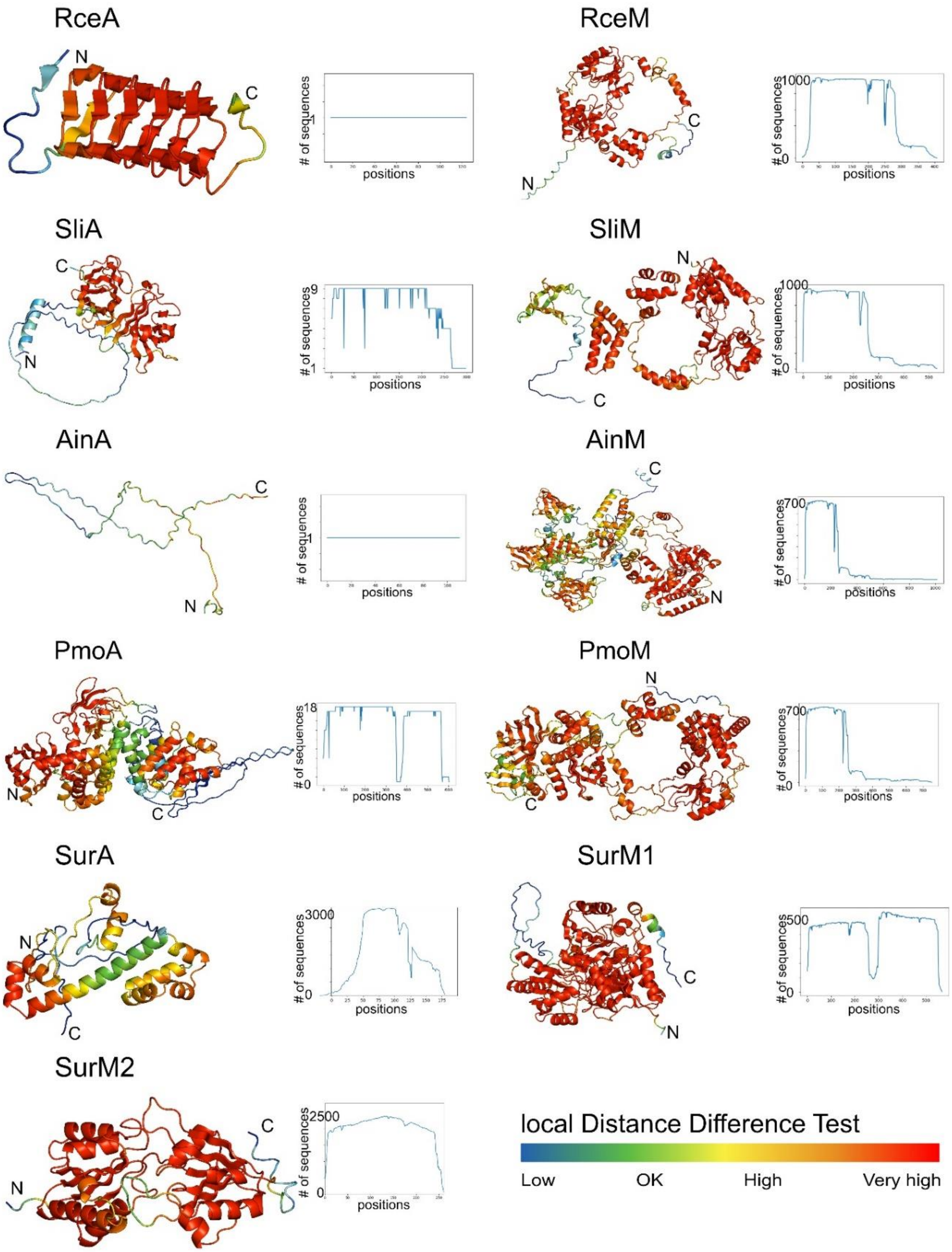

**Figure S5. Sequence similarity network of α-*N*-methyltransferase domains, cutoff 1 x $10^{-60}$.** The sequence similarity network (SSN) consists of sequences from a BLASTP search of the α-*N*-methyltransferase domain of OphMA, sequences of methyltransferases involved in fungal borosin biosynthesis (orange), and sequences of YabN_like methyltransferases (maroon). The network was built off of a local alignment-based all-vs-all analysis using BLASTP and visualized in Cytoscape. A cutoff value of 1 x $10^{-60}$ was applied. A select set of nodes are colored by the taxonomic groups to which their host organisms are assigned.

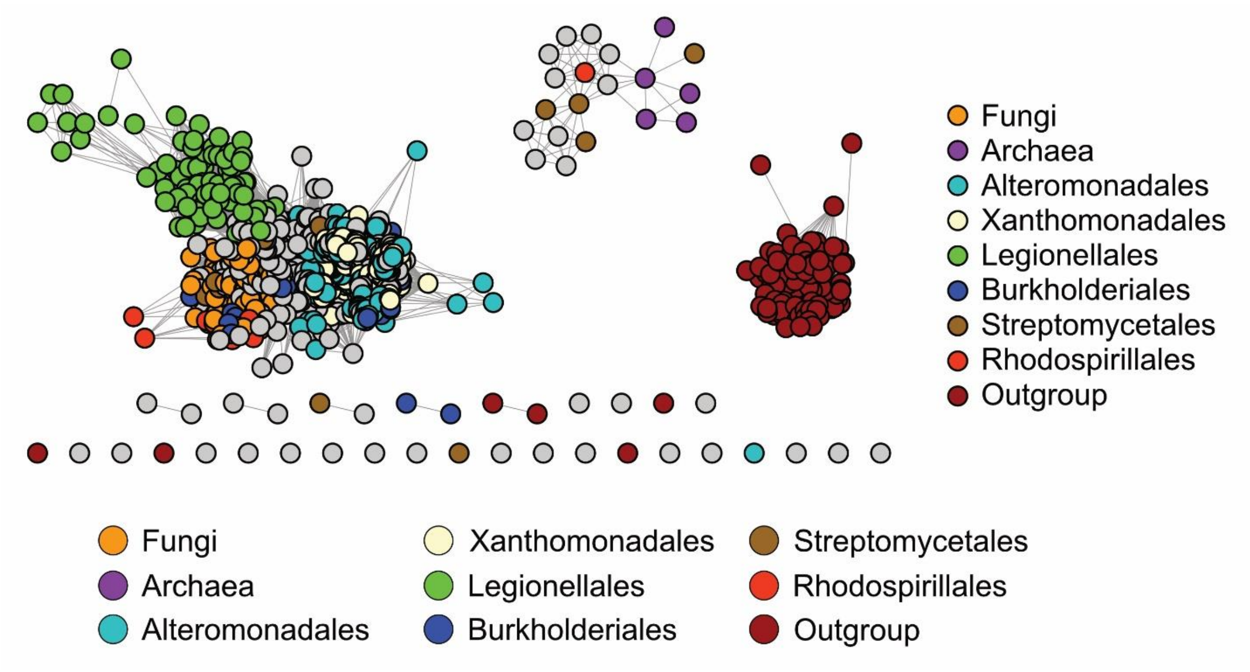

**Figure S6. Sequence similarity network of α-*N*-methyltransferase domains, cutoff 1 x 10$^{-90}$**. The sequence similarity network (SSN) consists of sequences from a BLASTP search of the *N*-methyltransferase domain of OphMA, sequences of methyltransferases involved in fungal borosin biosynthesis (orange), and sequences of YabN_like methyltransferases (maroon). The network was built off of a local alignment-based all-vs-all analysis using BLASTP and visualized in Cytoscape. A cutoff value of 1 x 10$^{-90}$ was applied. A) A select set of nodes are colored by the taxonomic groups to which their host organisms are assigned. B) Nodes are colored by the identified borosin type.

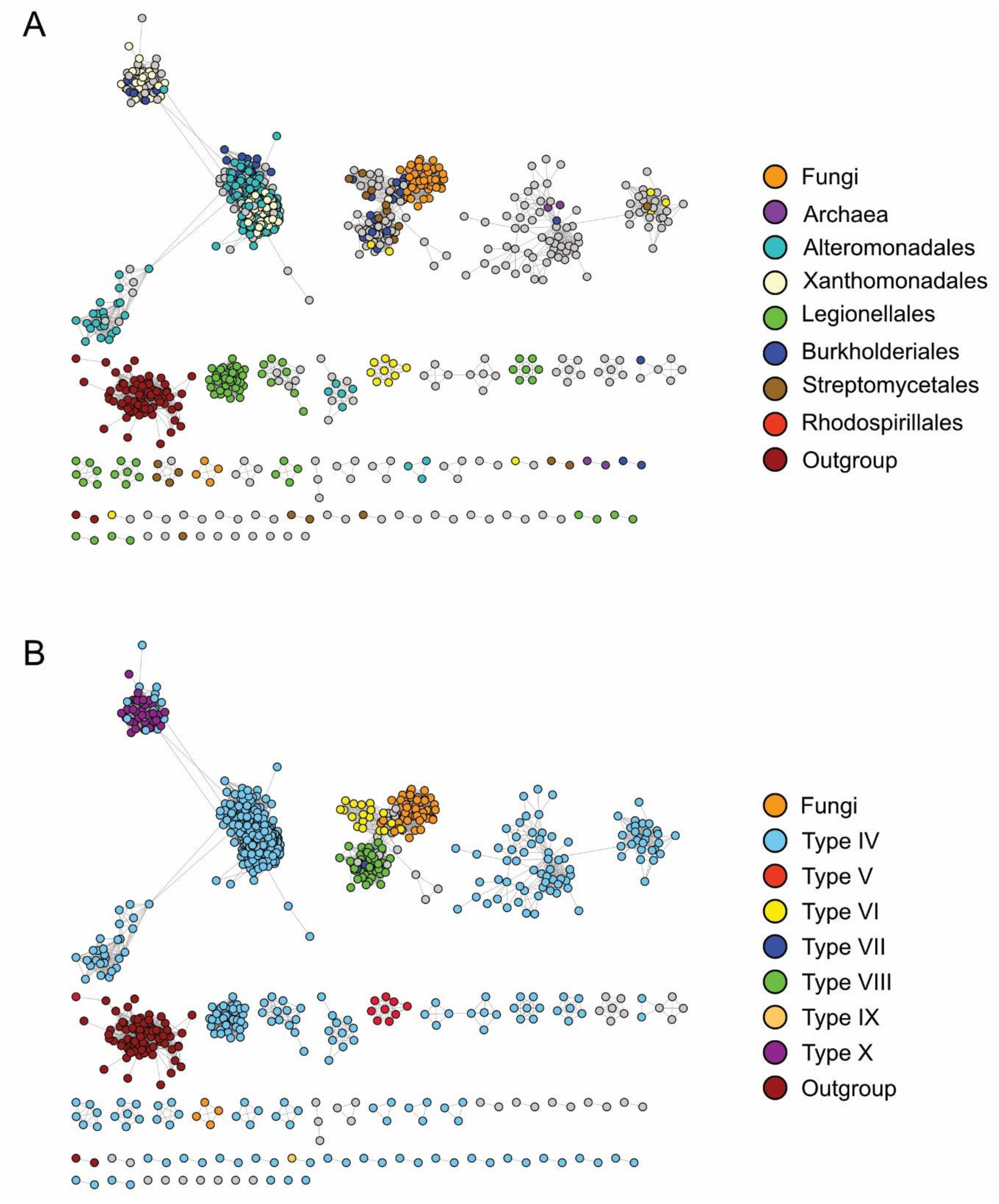

**Figure S7. Sequence similarity network of α-*N*-methyltransferase domains, cutoff 1 x $10^{-80}$**. The sequence similarity network (SSN) consists of sequences from a BLASTP search of the *N*-methyltransferase domain of OphMA, sequences of methyltransferases involved in fungal borosin biosynthesis (orange), and sequences of YabN_like methyltransferases (maroon). The network was built off of a local alignment-based all-vs-all analysis using BLASTP and visualized in Cytoscape. A cutoff value of 1 x $10^{-80}$ was applied. **(a)** A select set of nodes are colored by the taxonomic groups to which their host organisms are assigned. **(b)** Nodes are colored by the identified borosin type.

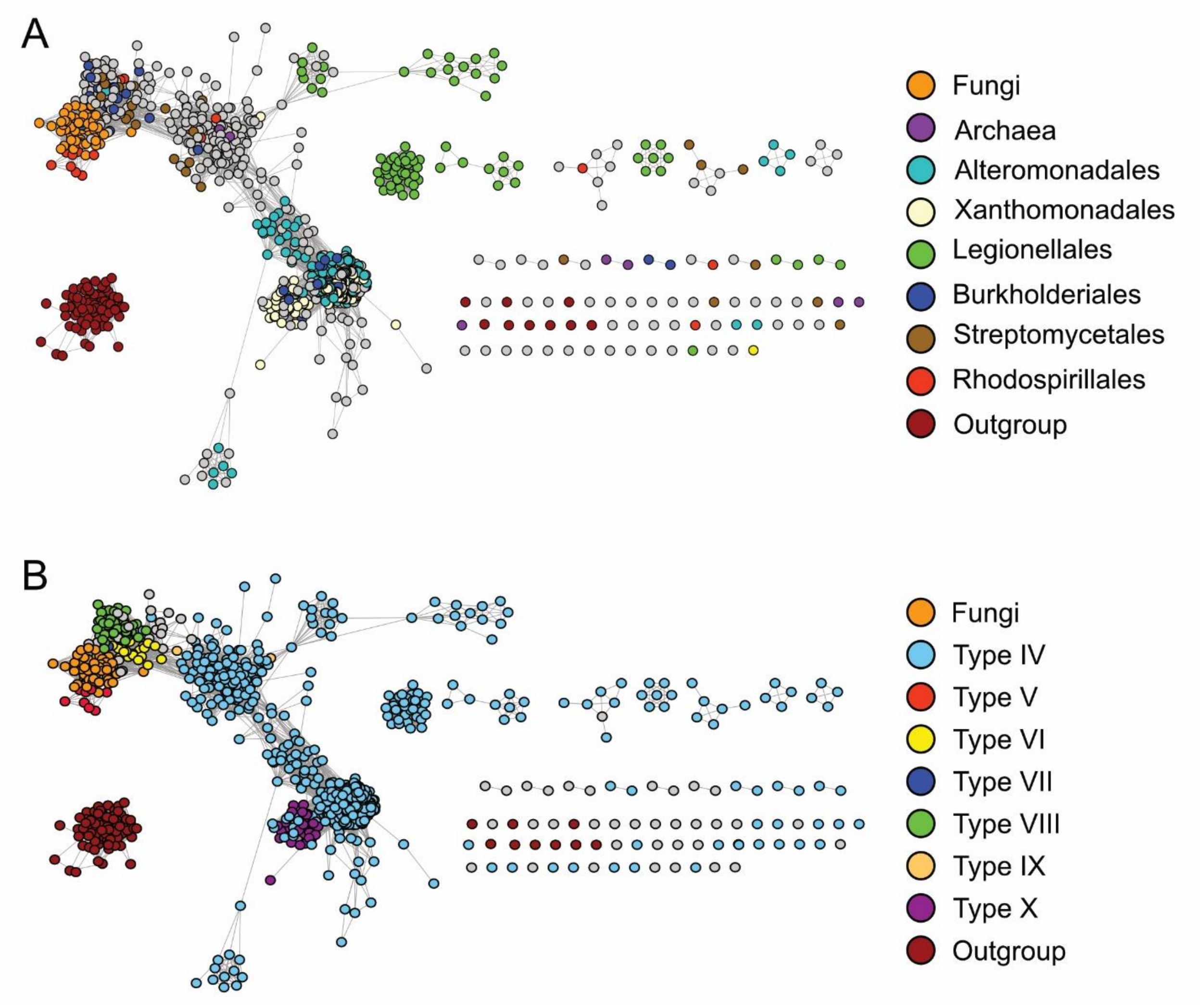

**Figure S8. BiG-SCAPE network of putative borosin BGCs colored by taxonomic group**. Genome assemblies in the RefSeq database corresponding to protein accession IDs of the putative borosins from the BLASTP search were analyzed using a local copy of antiSMASH containing the BorosinMT rule to identify putative borosin BGCs. The identified BGC regions were clustered in BiG-SCAPE then imported into and visualized in Cytoscape. A cutoff value of 0.50 was applied. A select set of nodes are colored by taxonomic group. Letters denote the designated BGC family for the clusters. Only families with more than eight members are designated. This cutoff was used based off the observed decrease in diversity of BGC domain composition in families with fewer members.

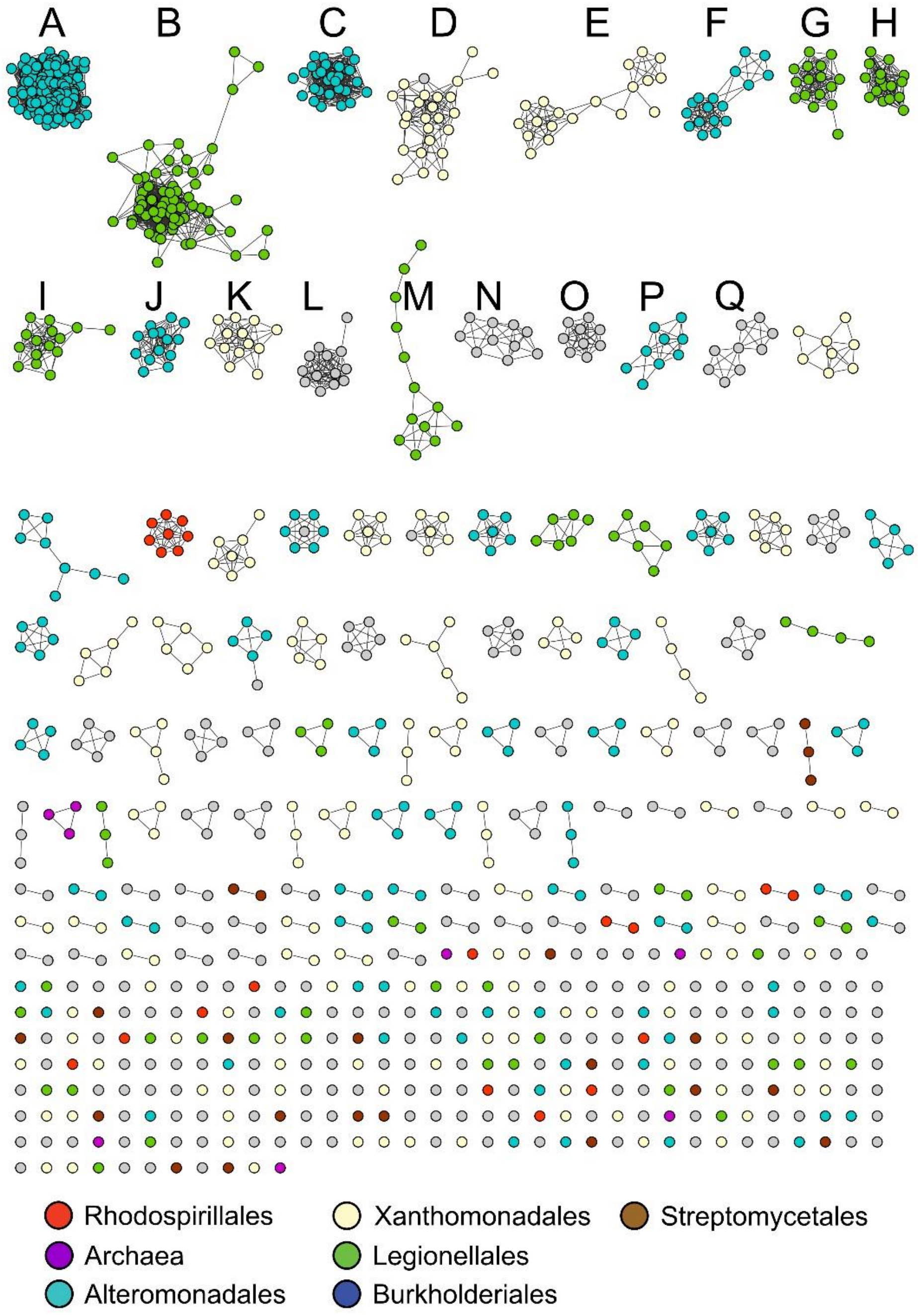

**Figure S9. BiG-SCAPE network of putative borosin BGCs colored by borosin type**. Genome assemblies in the RefSeq database corresponding to protein accession IDs of the putative borosins from the BLASTP search were analyzed using a local copy of antiSMASH containing the BorosinMT rule to identify putative borosin BGCs. The identified BGC regions were clustered in BiG-SCAPE then imported into and visualized in Cytoscape. A cutoff value of 0.50 was applied. Nodes are colored by the identified borosin type.

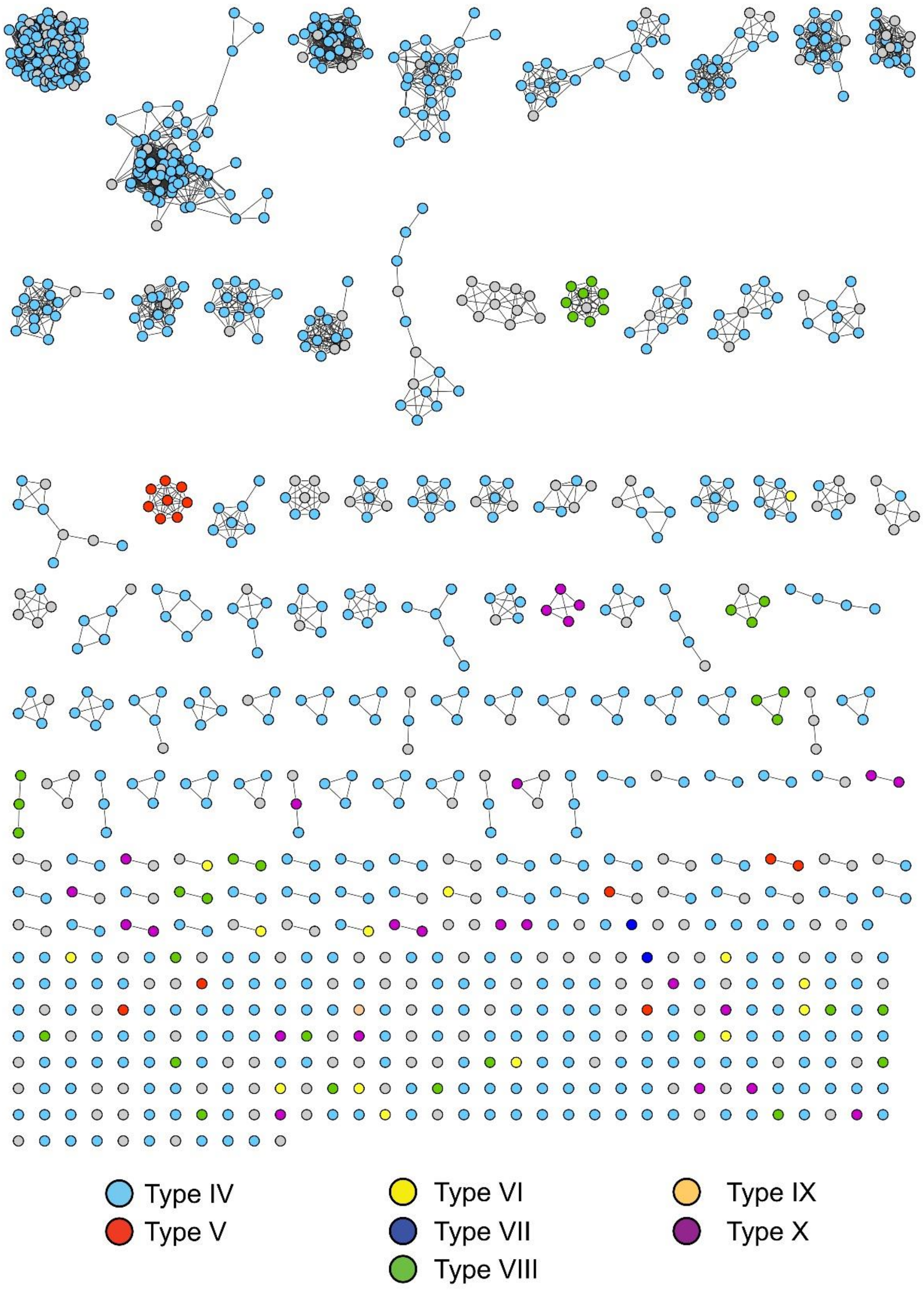

**Fig S10. Putative BGCs of characterized 'split' borosins.** ORFs identified by NCBI are included. ORF names reflect NCBI annotations, and accession numbers have been provided in parentheses.

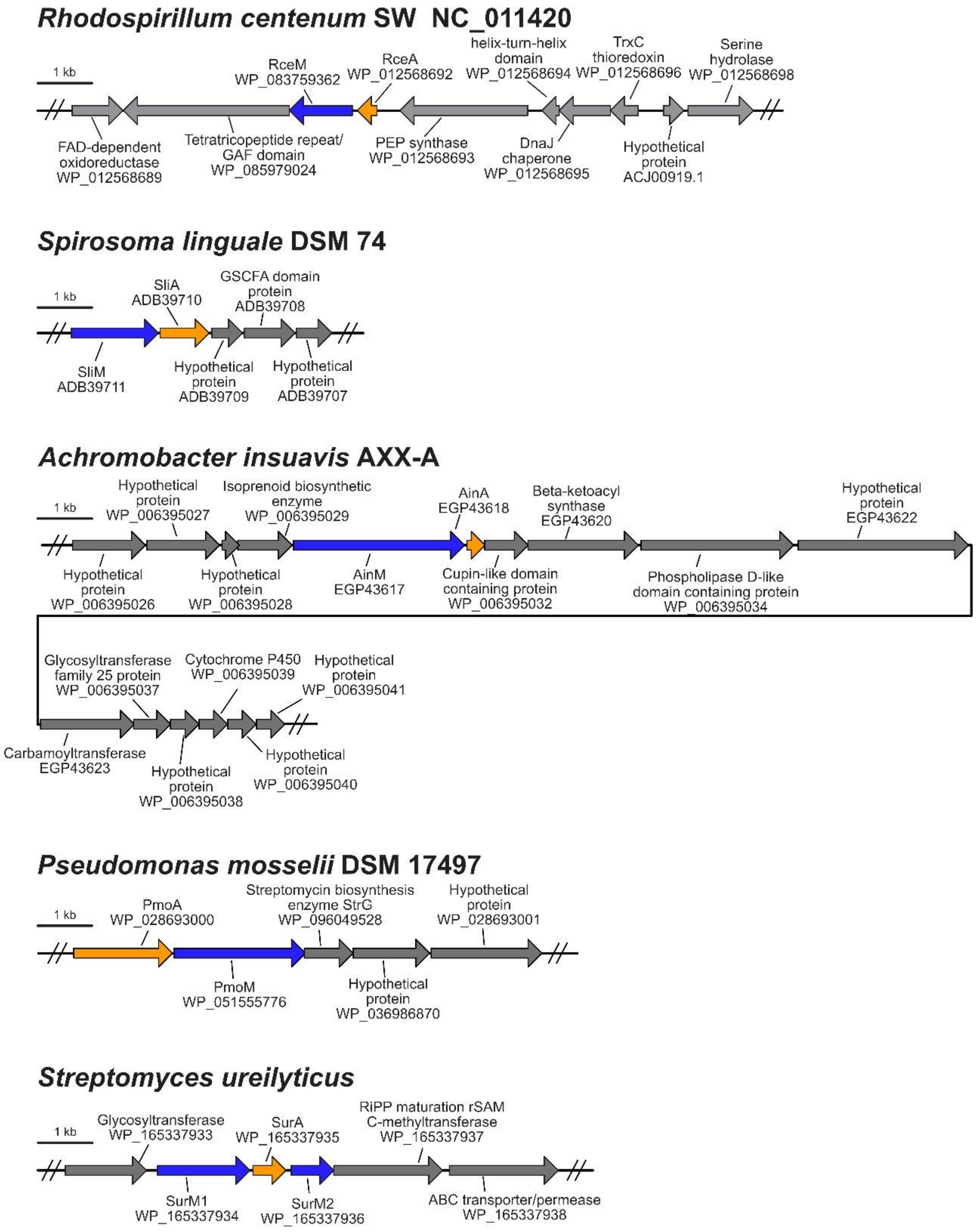

**Fig. S11. SDS-PAGE gel of purified α-*N*-methyltransferase and precursor proteins from *E. coli*.** 5 µg of each purified protein was run on a 4-12% NuPAGE Bis-Tris gel and stained with Coomassie brilliant blue. The name of the protein is followed by the duration of *E. coli* expression. Protein-of-interest is marked by "◀". Lane 1: RceM, 24 hrs; Lane 2: RceA, 24 hrs; Lane 3: RceA S78F, 48 hrs; Lane 4: SliM, 48 hrs; Lane 5: SliA, 24 hrs; Lane 6: AinM, 24 hrs; Lane 7: AinA, 24 hrs; Lane 8: PmoM, 24 hrs; Lane 9: PmoA, 24 hrs; Lane 10: SurM1, 48 hrs; Lane 11: SurM2, 24 hrs; Lane 12: SurA.

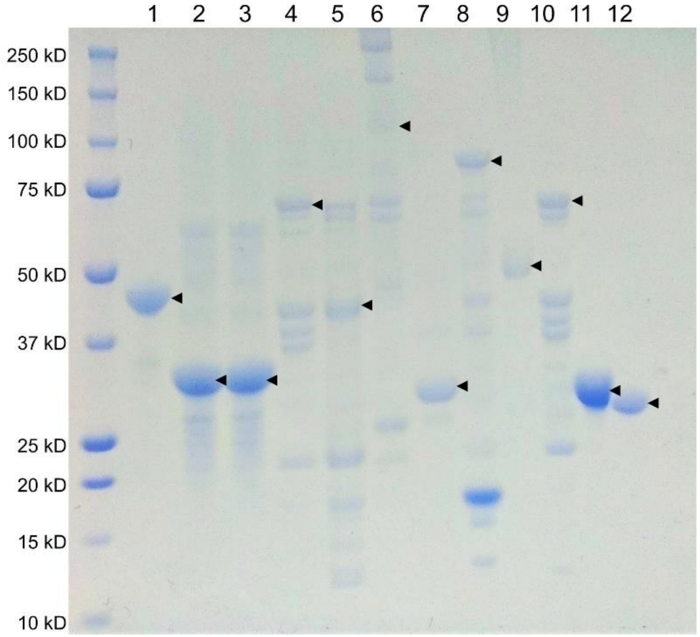
